# Tumor removal limits prostate cancer cell dissemination in bone and osteoblasts induce cancer cell dormancy through focal adhesion kinase

**DOI:** 10.1101/2022.09.02.506436

**Authors:** Ruihua Liu, Shang Su, Jing Xing, Ke Liu, Yawei Zhao, Mary Stangis, Diego P. Jacho, Eda D. Yildirim-Ayan, Cara M. Gatto-Weis, Bin Chen, Xiaohong Li

## Abstract

**Background:** Disseminated tumor cells (DTCs) can enter a dormant state and cause no symptoms in cancer patients. On the other hand, the dormant DTCs can reactivate and cause metastases progression and lethal relapses. In prostate cancer (PCa), relapse can happen after curative treatments such as primary tumor removal. The impact of surgical removal on PCa dissemination and dormancy remains elusive. Furthermore, as dormant DTCs are asymptomatic, dormancy-inducing can be an operational cure for preventing metastases and relapse of PCa patients.

**Methods:** We used a PCa subcutaneous xenograft model and species-specific PCR to survey the DTCs in various organs at different time points of tumor growth and in response to tumor removal. We developed *in vitro* 2D and 3D co-culture models to recapitulate the dormant DTCs in the bone microenvironment. Proliferation assays, fluorescent cell cycle reporter, qRT-PCR, and Western Blot were used to characterize the dormancy phenotype. We performed RNA sequencing to determine the dormancy signature of PCa. A drug repurposing algorithm was applied to predict dormancy-inducing drugs and a top candidate was validated for the efficacy and the mechanism of dormancy induction.

**Results:** We found DTCs in almost all mouse organs examined, including bones, at week 2 post-tumor cell injections. Surgical removal of the primary tumor reduced the overall DTC abundance, but the DTCs were enriched only in the bones. We found that osteoblasts, but not other cells of the bones, induced PCa cell dormancy. RNA-Seq revealed the suppression of mitochondrial-related biological processes in osteoblast-induced dormant PCa cells. Importantly, the mitochondrial-related biological processes were found up-regulated in both circulating tumor cells and bone metastases from PCa patients’ data. We predicted and validated the dormancy-mimicking effect of PF-562271, an inhibitor of focal adhesion kinase (FAK) *in vitro*. Decreased FAK phosphorylation and increased nuclear translocation were found in both co-cultured and PF-271-treated C4-2B cells, suggesting that FAK plays a key role in osteoblast-induced PCa dormancy.

**Conclusions:** Our study provides the first insights into how primary tumor removal enriches PCa cell dissemination in the bones, defines a unique osteoblast-induced PCa dormancy signature, and identifies FAK as a PCa cell dormancy gatekeeper.

## Background

Prostate cancer (PCa) is the most frequently diagnosed cancer and the second leading cause of cancer-related deaths in men in the United States [1]. Up to 90% of patients with advanced-stage PCa develop bone metastases, but only 10% are diagnosed with bone metastases at initial diagnosis [2–4]. Primary PCa cells can disseminate early and remain dormant in distant organs before reactivation, causing metastasis and recurrence [5–8]. Despite various therapies, including prostatectomy and enzalutamide treatment, 20-45% of these patients relapse years later [9–14]. The mechanisms of PCa cell dissemination, dormancy, and reactivation, particularly the impact of treatments on these processes, are poorly understood.

Tumor dormancy can be categorized as tumor mass dormancy and cellular dormancy [5, 6, 15]. Tumor mass dormancy is when a small, undetectable tumor reaches an equilibrium between proliferation and cell death, resulting in no progression or overt metastases. Cellular dormancy occurs when a cancer cell enters a state of quiescence with significantly reduced proliferation. Tumor dormancy is often a combination of both, and the mechanisms of tumor dormancy can be attributed to cancer cell-intrinsic and acquired quiescence resulting from external stimuli, including the microenvironment and drug treatment [16–26]. Dormant cancer cells are under cell cycle arrest, are neither actively proliferative nor apoptotic, and express reduced levels of Ki67 and cell cycle-related genes but higher levels of dormancy makers, such as the nuclear receptor subfamily 2 group F member 1 (NR2F1) [17, 22, 27–30]. Due to cancer cell heterogeneity, distinct dormancy markers are expected and identified under different microenvironment settings and for various cancers. Therefore, our study aims to investigate the effects of tumor removal on PCa cell dissemination, determine clinically relevant dormant PCa profiles, and explore approaches to harness PCa cell dormancy.

We conducted a comparative analysis of the abundance of the disseminated tumor cells (DTCs) in various organs of mice with or without tumor removal. Our findings using an *in vitro* mixed co-culture model revealed that only osteoblasts induced and maintained PCa cell dormancy. We profiled the gene expression signature of the dormant C4-2B cells and identified that the top six enriched biological processes from the significantly down-regulated genes were all mitochondria-related. Interestingly, we also discovered through PCa patient dataset analyses that mitochondria-related biological processes were enriched in the up-regulated genes of circulating tumor cells. This indicates that dormancy could hinder PCa progression in patients and, conversely, a relapse could result from dormancy reactivation. However, direct mitochondrial inhibition failed to induce PCa cell dormancy. To identify a drug that mimics PCa cell dormancy in vitro, we used a novel drug prediction artificial intelligence (AI) platform and predicted PF-562271 (PF-271), a potent ATP-competitive and reversible focal adhesion kinase (FAK) inhibitor [31–33]. We validated the dormancy-mimicking effect of PF-271 by inhibiting FAK phosphorylation and inducing FAK nuclear translocation in vitro. Overall, our study provides the first evidence of the effects of primary tumor removal on PCa cell dissemination, identified the signature of the osteoblast-induced C4-2B cell dormancy, and discovered a drug that induces PCa cell dormancy *in vitro*.

## Materials and Methods

### Cell cultures

Unless specified, cell lines were purchased from ATCC (American Type Culture Collection, Manassas, VA). Mouse osteoblast MC3T3-E1 cells (gift from Dr. Bart Williams, Van Andel Institute) were maintained in α-MEM supplemented with 10% fetal bovine serum (FBS). Mouse macrophage Raw264.7 cells (gift from Dr. Cindy Miranti, Arizona State University), mouse fibroblast NIH3T3 cells, and human embryonic kidney 293FT cells were maintained in DMEM/HG supplemented with 10% FBS. Mouse mesenchymal progenitor OP-9 cells (gift from Dr. Tim Triche, Van Andel Institute) were cultured in MEM-α supplemented with 20% FBS. Murine Osteocyte-like Cell Line MLO-Y4 were grown in α-MEM supplemented with 5% FBS and 5% calf serum (CS) on the plate pre-coated with rat tail type 1 collagen (0.15 mg/mL, Corning, Corning, NY). Human PCa cell lines C4-2B (gift from Dr. Leland Chung and purchased from ATCC), PC-3, DU-145, and 22Rv1 were cultured in RPMI-1640 supplemented with 10% FBS. Human primary umbilical vein endothelial cells (HUVEC) were cultured in vascular cell basal media supplemented with BBE endothelial cell growth kit components. All the cells were maintained in a humidified 5% CO_2_ incubator at 37°C except for hFOB1.19 (34°C + 5% CO_2_). All the cells were authenticated using STR profiling in LabCorps (Burlington, NC) and maintained mycoplasma-free. Stable cell lines were generated via lentivirus infection, as described in the sections below.

### Lentivirus packaging and stable cell line generation

Plasmid pLenti-GFP-Blast (gift from Dr. Wei Wu, Tsinghua University) was used for general GFP labeling, and pLenti-PGK-Neo-PIP-FUCCI (Fluorescent Ubiquitination-based Cell Cycle Indicator) (Addgene # 118616) was used for accurate cell cycle phase indicator [34]. GFP-mouse FAK plasmid was a gift from Dr. Steve Lim, University of Alabama at Birmingham. We subcloned it into pLenti-GFP-blast (replacing the original GFP cassette) to get pLenti-GFP-mFAK plasmid for overexpressing GFP-mFAK in C4-2B cells. Luciferase-expressing lentiviral particles (puromycin resistance) were purchased from GeneCopoeia (Rockville, MD). All the other lentiviral supernatants for stable line generation were prepared in 293FT cells using ViraPower™ Lentiviral Gateway™ Expression Kit (Thermo Fisher) following the manual. C4-2B were infected with virus particles/supernatants, selected with blasticidin, puromycin, or neomycin for 7-10 days to get stable cell populations, and later maintained at lower antibiotic concentrations. C4-2B/GFP and C4-2B/PIP-FUCCI cells were confirmed via fluorescence microscopy.

### Reagents, Chemicals, Kits, and plasmids

Listed in **Supplementary Table S1**.

### Mouse models

NSG-SCID mice were originally purchased from Jackson Laboratory (Bar Harbor, Maine, USA) and bred and maintained in our animal facility. The research protocols were approved by the Institutional Animal Care and Use Committee (IACUC) under protocol numbers 400066/400131 at the University of Toledo. NSG males, six to eight weeks old, were used for xenografting human PCa cells. For subcutaneous xenograft models, 1 million C4-2B/Luc cells were suspended in 50 μL PBS, mixed with 50 μL Matrigel (Corning, USA), and subcutaneously injected into the left flank of the NSG mice. Luminescent imaging (IVIS, Perkin Elmer, Waltham, Massachusetts, USA) was performed in a blinded manner once per week to monitor tumor growth. D-Luciferin (GoldBio, St Louis, MO) was injected intraperitoneally at a dose of 150 μg/g and incubated for 10 min before scanning in the IVIS® Spectrum In Vivo Imaging System (PerkinElmer, Waltham, MS).

At 2 weeks (wks) post tumor xenografting, the xenografted mice with similar sizes of tumors were randomized into three groups for DTC detection at different time points. In Group 1, the mice were euthanized for organ harvest and DTC detection at 2 wks. Group 2 served as the control group. In Group 3, the mice received surgical removal of the tumors at 2 wks. The mice in Group 2/3 were monitored, imaged, and euthanized at different time points, i.e., 6, 8, and 18 wks post PCa injections. Note that there are no time points beyond 8 wks for mice in Group 2 because xenografted tumors reached the euthanization criteria according to the IACUC protocol. Various organs and tissues were harvested at endpoints for genomic DNA extraction and species-specific PCR to detect the disseminated human PCa cells.

For determination of DTC distribution in long bones such as tibiae and femurs, we first removed the skeletal muscles, then cut the bones open at one end and set it into a 0.5 mL open-bottomed centrifuge tube, which was placed inside a 1.5 mL centrifuge tube. The bone marrows were centrifuged out at the maximum speed in a tabletop Eppendorf centrifuge and collected in the 1.5 mL centrifuge tube for further uses, either DNA extraction or immunofluorescence (IF) staining.

### Detection of the disseminated tumor cells (DTCs)

Total DNA was extracted from various mouse organs using the NucleoSpin Tissue kit (Macherey-Nagel Inc., Allentown, PA, USA). The DNAs were used for species-specific PCR with primers against human α-satellite and mouse plakoglobin genomic regions [35]. To semi-quantify the DTCs, a standard curve was first established by performing species-specific PCR from total DNA extracted from the mixtures of C4-2B, at 0, 1, 10, 100, 1,000, 10 thousand (k), or 100 k of cells, and 3 million MC3T3-E1 cells. 2 µg DNA for soft organs and 600 ng for tibia or femur were used in the PCR reactions for human α-satellite; 500 ng DNA was used for mouse plakoglobin. Primers are listed in **Supplementary Table S2**. Gel images for verification of species specificity of primers were provided in **Supplementary Data**. The final PCR products were loaded on 2% agarose gel, and the band intensities on raw TIFF images without oversaturation were quantified using Image J (FIJI 1.53c). Human cell numbers were relatively quantified via linear regression based on the intensity ratios of human α-satellite to mouse plakoglobin. This standard is used to semi-quantify the human cell DTC from the harvested mouse tissues. All DTC quantifications were performed in a blinded manner, and the estimated DTC numbers were listed in **Supplementary Table S3**. Representative images were shown in **Figure 1** and **Supplementary Figure S1.**

**Figure 1.**
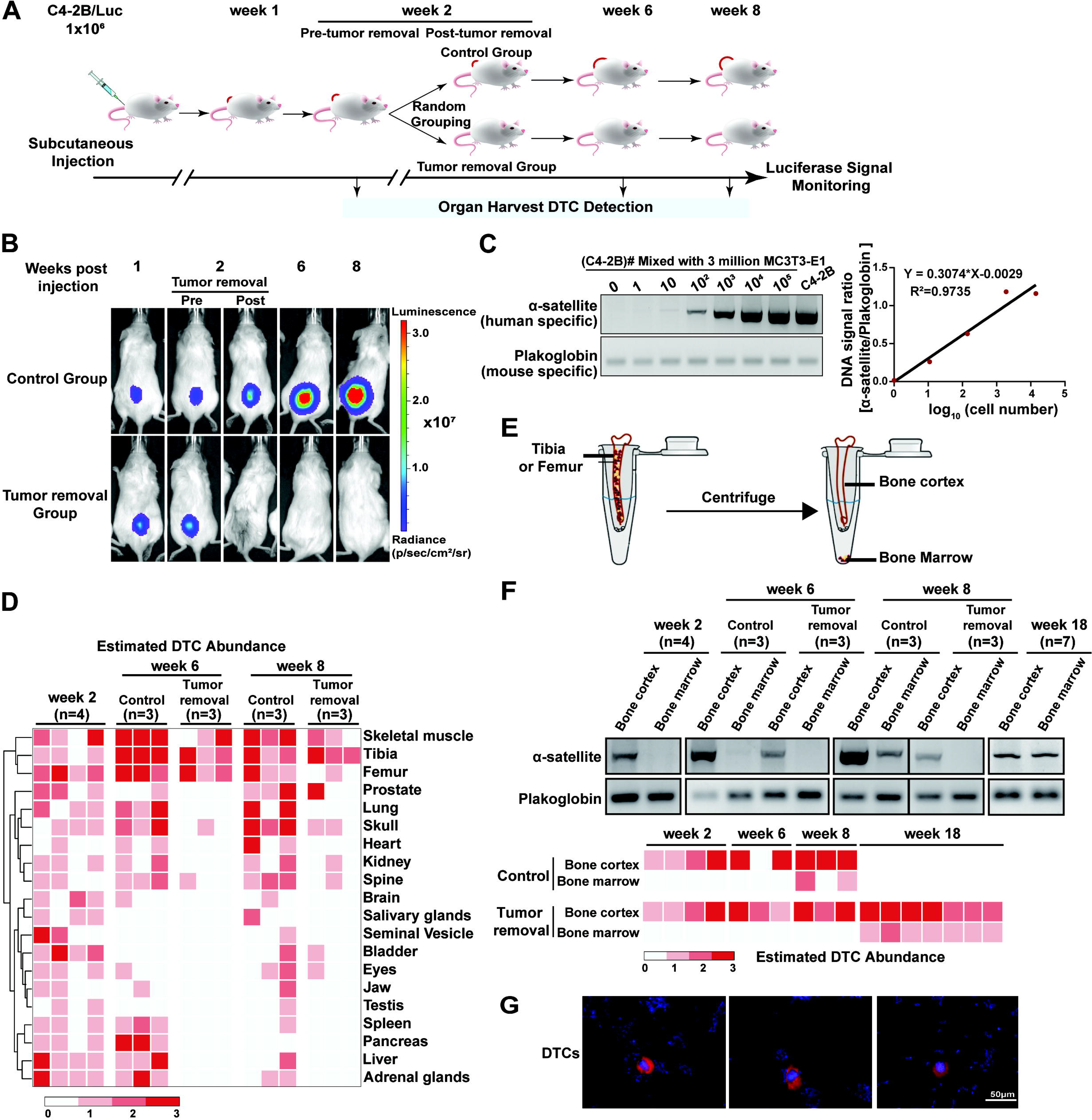
Primary tumor removal limited PCa DTC in the bone. **A**. The schematic diagram of experimental design. **B**. Representative weekly luminescent images of subcutaneous C4-2B/Luc tumors in mice. Pre- and Post-tumor removal indicate the time points right before the surgery and after recovery in the same week. **C**. The standard curve for DTC number estimation is based on band intensities from the species-specific PCR of genomic DNA. Plakoglobin was used here to normalize the genomic input. **D**. The heatmap for estimated abundance of DTCs in different tissues. Color scale: 0, not detected; 1, 1-10 DTCs; 2, 10-100 DTCs; 3, >100 DTCs. **E**. Diagram of separation of the bone cortex from the bone marrow of mouse tibiae or femurs. **F**. DTC abundance in mice’s bone cortex and bone marrow post xenografting. Upper panel, representative species-specific PCR gels; lower panel, heatmap for estimated DTC abundance. For genomic DNA loading, 2 μg total genomic DNA for α-satellite PCR of bone marrow, 0.6 μg for α-satellite PCR of bone cortex, and 0.5 μg for all plakoglobin PCR were used. **G**. Representative pictures of IF staining of the bone marrow cells using a human species-specific mitochondrial antibody (red). DAPI (blue) was used to show the nucleus. Scale bar, 50 μm. Experiments were repeated at least twice with triplicates.

To validate that the bone marrow DTC is intact, we resuspended the bone marrow cells in the 1.5 mL centrifuge tube, lysed red blood cells with ACK Lysing Buffer (ThermoFisher), and further mounted them onto the slides using Cytospin. Cells on the slides were fixed with 4% Paraformaldehyde (PFA), permeated with 0.2% Triton, and blocked by 3% BSA. Slides were further incubated overnight at 4^◦^C with primary antibody against human mitochondrial marker. On the second day, the slides were washed three times and further incubated with Alexa Fluor 568 conjugated goat-anti-mouse IgG secondary antibody. Then the cells were stained with DAPI solution and washed three times before mounting with Fluoromount-G® Anti-Fade Mounting Medium (Southern Biotech, Birmingham, AL). Images were captured using Zeiss Observer Z1 confocal microscope (Carl Zeiss, Oberkochen, Germany) and processed using Image J.

### *In vitro* co-culture models

For the mixed co-cultures, stromal cells, MC3T3-E1 (3×10^5^/well), RAW264.7 (1×10^5^/well), NIH3T3 (5×10^4^/well), OP-9 cells (3×10^5^/well), MLO-Y4 (3×10^5^/well), hFOB1.19 (3×10^5^/well), or HUVEC (3×10^5^/well) were seeded in a 6-well plate and allowed for adherence for 20 to 24 hours at 37°C (hFOB1.19 at 34°C). PCa cells (C4-2B, PC-3, 22Rv1, or DU-145) labeled with GFP (1×10^4^/well) were then added. For the transwell co-culture, MC3T3-E1 cells (3×10^5^/well) were seeded in the low chamber, while C4-2B cells (1×10^4^/well) were added in the upper chamber. The culture media for corresponding stromal cells were used for co-culture unless specified. Cells were harvested at certain times for further analysis.

To separate C4-2B/GFP prostate cancer cells from co-cultures with osteoblast (MC3T3-E1 or hFOB1.19), we aspirated the media from the 3-day co-culture, and gently rinsed the cells once with 1 mL PBS. Subsequently, we detached the C4-2B/GFP cells into 1 mL PBS by carefully pipetting with a 1 mL pipette. The cell suspension in PBS was transferred to a new well in a 6-well plate, while an additonal 1 mL PBS was added to the remaining osteoblasts. To ensure the successful separation between C4-2B (GFP-positive in all cells) and osteoblasts (no GFP expression), we examined both fractions under fluorescence microscopy. This separation method consistently yielded near 100% efficiency, as validated by fluorescence microscopy and genomic DNA PCR (shown in **Figure 2I** for MC3T3-E1 co-culture separation and **Supplementary Figure S3A** for hFOB1.19 co-culture separation).

**Figure 2.**
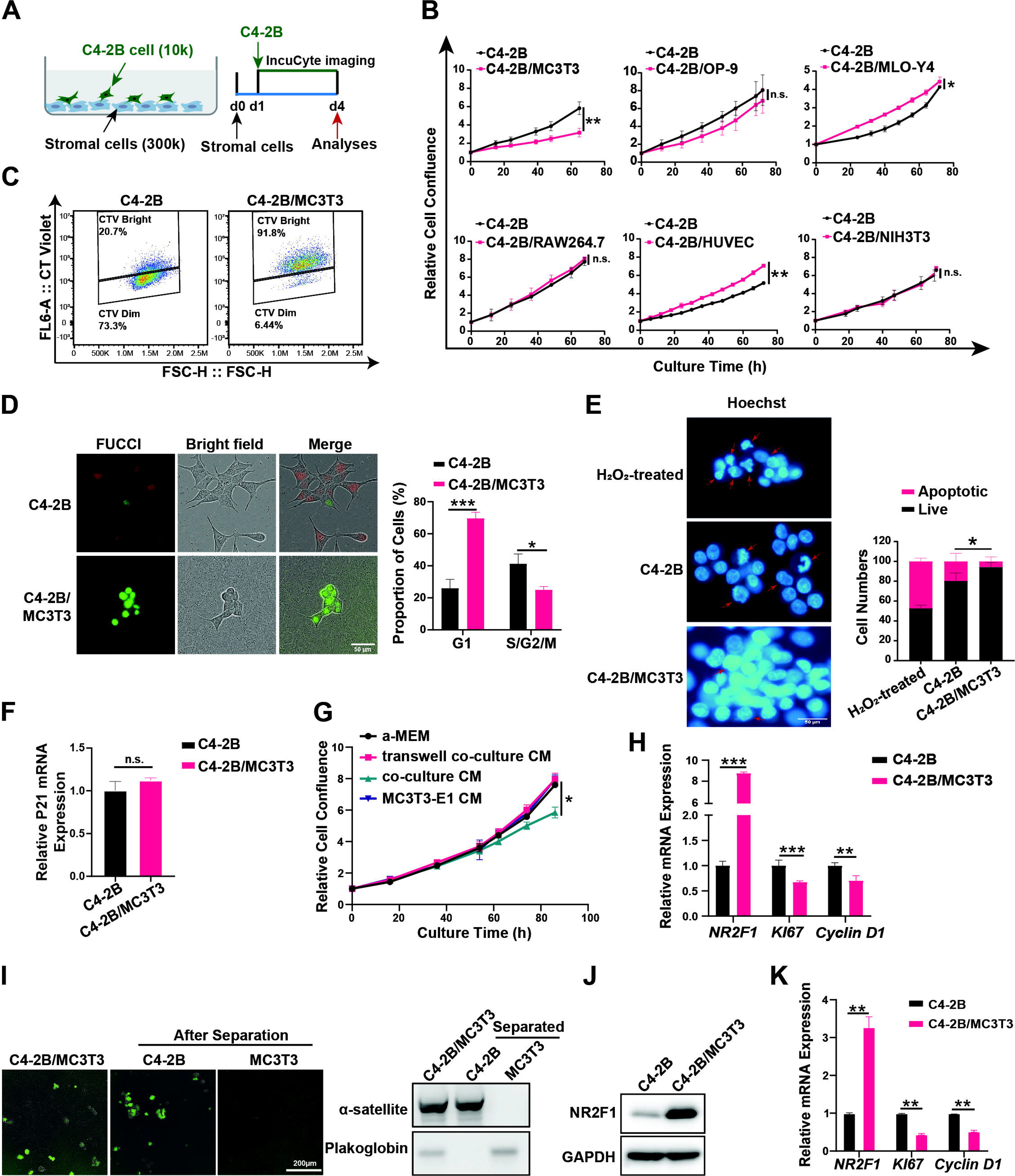
Osteoblasts induced PCa cell dormancy via physical contacts. **A**. The schematic diagram of the co-culture model. Relative cell confluence of GFP-labelled C4-2B cells. GFP-positive areas of each time point were normalized to the ones at the co-culture seeding. **C**. Cell division of C4-2B cells cultured alone or co-cultured with MC3T3-E1 cells. C4-2B cells were stained with CellTrace™ Violet (CT Violet) before seeding. FACS analyses of the CT Violet signal intensities were conducted 72 h. **D**. Cell cycle analyses with PIP-FUCCI reporter. Cells with only green color, G1; Cells in red, late S/G2/M. Representative fluorescent and bright field pictures were shown. **E**. C4-2B cells cultured alone or co-cultured with MC3T3-E1 for 72 h or treated with H_2_O_2_ for 2h (positive control for apoptosis) were stained with Hoechst 33342 for examination of apoptotic cells (red arrows) based on the nuclear morphology. **F/H**. C4-2B cells were cultured alone or co-cultured with MC3T3-E1 for 72 h. The mRNA expressions of marker genes p21 (**F**), NR2F1, Ki67, and Cyclin D1 (**H**) were examined using qRT-PCR. Human-specific primers were used, and relative expressions were calculated and plotted after normalization with GAPDH expressions. **G**. C4-2B cells were treated with different conditioned media (CM) as indicated for 72 h. **I**. Separation of C4-2B cells (GFP labeled) from co-culture of MC3T3-E1 cells (not labeled) was confirmed using both fluorescence microscopy (left) and species-specific PCR (right). **J**. Immunoblotting of NR2F1 in C4-2B cultured alone or separated from the co-cultures. **K**. The mRNA expression changes of marker genes in the 3D co-culture model. All the experiments were independently repeated at least twice. The data in curves and bar plots are presented as the mean ± SD of each set of triplicate samples. Scale bars, 50 μm (**D&E**), 200 μm (**I**). * p < 0.05, ** p < 0.01, *** p < 0.001.

For conditioned media (CM) treatment, we collected CM at day 3 from C4-2B, MC3T3-E1, the mixed co-culture of C4-2B and MC3T3-E1 cells, or the transwell co-cultures (MC3T3-E1 cells in the well and C4-2B cells in the inserts with 0.4 μm pore membrane). CM was mixed with fresh α-MEM media at a 1:1 ratio to treat C4-2B cells. Cell proliferation was then monitored using Incucyte S3 based on the GFP signals for 3 days.

The 3D co-culture protocol was modified based on previous studies [36–39]. Briefly, C4-2B cells or the mixture of MC3T3-E1 and C4-2B (30:1) were encapsulated at 10^6^ cells/mL seeding density within collagen type I solution (Corning, USA) and neutralized to pH 7∼8 with chilled 1N NaOH to a final collagen concentration of 3 mg/mL. The cell-collagen solutions were added to the silicon chambers and polymerized at 37°C for 1 hour, proceeded with adding 1 mL complete media and incubated for 3 days at 37°C and 5% CO_2_ before harvest for gene expression analyses.

### Cell division and cell cycle analyses

To monitor cell divisions, the C4-2B or PC-3 cells were labeled by CellTrace™ Violet Cell Proliferation Kit (ThermoFisher) according to the manufacturer’s manual. This chemical dye can be saturated in cells and is diluted into half every time the cell divides, thus indicating the cell division [40]. The cells with various dye intensities (suspended at a density of 1 x 10^6^ cells/mL in PBS) were examined using flow cytometry on a FACS Cytek Aurora flow cytometer. Data were analyzed with FlowJo software. Gating parameters were determined using non-stained negative control samples.

C4-2B/PIP-FUCCI (PCNA-Interacting Protein degron version of Fluorescent Ubiquitination-based Cell Cycle Indicator) cells were used to monitor the cell cycle [1]. Note that PIP-FUCCI is different from original FUCCI, i.e, cells with pure green and red fluorescence are in the G0/G1 and late S/G2/M phases, respectively. We counted the total cells and cells with either green or red color under 20x magnitude at 6 random areas per group. Cell counting was performed in a blinded manner.

### Cell Apoptosis Assay

For cell apoptosis, cells were stained with Hoechst 33342 (5 μg/mL, ThermoFisher) at 37°C for 30 min, refreshed with fresh media, and captured under fluorescence microscopy. The cell nuclear morphologies from multiple views were recorded and apoptotic nuclei were counted in a blinded manner. C4-2B cells treated with 1 mM H2O2 for 2 hours were used as the positive control for apoptosis [41].

### Mitochondrial gene copy analysis

After 3 days’ cultures, the cultured alone C4-2B cells or the mixed co-cultured C4-2B/MC3T3-E1 were harvested for DNA extraction using the NucleoSpin Tissue kit (Macherey-Nagel Inc.). Primers against the nuclear and mitochondrial genomes were used for qRT-PCR (sequences provided in **Supplementary Table S2**) and the Ct values were used for calculating the mitochondrial gene copies per cell [42].

### RNA extraction and qRT-RCR

Total RNA extractions from cells were performed using TRIzol (ThermoFisher). RNA was further reversely transcribed into cDNA using the High-Capacity cDNA Reverse Transcription Kit (ThermoFisher) following the manufacturer’s protocol. The quantitative real-time PCR (qRT-PCR) was performed using SYBR Select Master Mix (Bio-Rad). Relative quantities of target mRNA expressions were performed on cycle threshold (Ct) values of target mRNAs and the reference gene GAPDH, using the 2^−ΔΔCt^ method. Primers are validated for species specificity and the sequences are listed in **Supplementary Table S2**.

### RNA-Sequencing

For RNA-Seq, total RNAs were extracted from cells cultured alone or the mixed co-culture using the RNeasy kit and were decontaminated of the DNA using RNase-Free DNase Set (Qiagen, Hilden, Germany). Libraries were prepared by the Van Andel Institute Genomics Core from 500 ng of total RNA using the KAPA stranded mRNA kit (v4.16) (Kapa Biosystems, Wilmington, MA). RNA was sheared to 300-400 bp. Before PCR amplification, cDNA fragments were ligated to IDT for Illumina UDI adapters (IDT, Coralville, IA). The quality and quantity of the libraries were assessed using a combination of Agilent DNA High Sensitivity chip (Agilent Technologies, Santa Clara, CA) and QuantiFluor® dsDNA System (Promega). Individually indexed libraries were pooled and 50 bp, paired-end sequencing was performed on an Illumina NovaSeq 6000 sequencer using 100 cycle sequencing kit (Illumina Inc., San Diego, CA) to a minimum read depth of 40 M reads/library. Base calling was done by Illumina RTA3 and the output of NCS was demultiplexed and converted to FastQ format with Illumina Bcl2fastq v1.9.0. The Raw read FASTQ files can be accessed via GEO (GSE210751).

We applied three pipelines, Xenome [43], XenofilteR [44], and Kallisto [45], to process the human-specific gene expression changes from the RNA-Seq data (FastQ) from the mixed co-culture samples. For Xenome, the raw reads were first filtered to keep the human-only reads and then aligned to the human reference genome (GRCh38 for humans) via HISAT2 [46], and read counts were called using FeatureCounts in the subread package [47]. For XenofilteR, the FastQ files were first split into files of 50 million sequences via the split2 function from the SeqKit toolkit [48], then aligned via HISAT2 to human (GRCh38) and mouse (GRCm38) reference genomes respectively. The sorted bam files for both human and mouse alignment were loaded into XenofilteR to filter out mouse reads and merged for read counts calling via FeatureCounts. For Kallisto, we merged human and mouse transcriptome reference sequences (downloaded from Gencode) [49] and created a cross-species transcriptome index for human-specific gene expression quantification. Raw reads from C4-2B cells cultured alone were also applied to the same three algorithms with the same parameter settings for comparative DE analyses. DESeq2 was used in all three pipelines to perform DE analyses (a gene with |log2FC| >=1 & padj < 0.05 was considered as a DE gene) based on the FeatureCounts files (Xenome and XenofilteR) or gene counts (Kallisto). Genes identified to be DE by at least two pipelines were considered for Gene Ontology (GO) and KEGG enrichment analyses in a blinded manner via EnrichR [50] or ShinyGO v0.61 [51]. The DE lists from all three pipelines were provided in **Supplementary Table S5**. Codes are available upon request.

### Western blotting

Western blotting experiments were performed as previously described [52]. Antibodies were purchased from various companies. The application, the concentration, and the sources of the antibodies were listed in **Supplementary Table S4**. An antibody was selected based on its validations, including positive and negative controls, and references. Original blot images were provided in **Supplementary Data**.

### Statistical analysis

The normality of data was determined before performing statistical analyses. For experiments without repeated measures, data were analyzed using a two-way analysis of variance (ANOVA) when there were two independent variables. Otherwise, a two-tailed Student’s t-test was used. The analyses on the statistical significance of all data sets were performed using GraphPad Prism 9. If not mentioned, a two-tailed p-value was computed, and statistical significance was set at p <0.05.

## Results

### Tumor removal limits disseminated tumor cells (DTCs) to the bones

We subcutaneously injected human C4-2B/Luc cells (stably expressing firefly luciferase) into immunodeficient NSG male mice and monitored tumor growth weekly using bioluminescent imaging and caliper measurements. At week 2 (the week numbers hereafter indicate weeks post PCa cell injection), all subcutaneous tumors reached similar sizes in host mice (within the range of 144+/-37 mm^3^). The mice were then randomly assigned to three groups (**Figure 1A**): Group 1 was sacrificed at the 2-week point, Group 2 (termed the Control group) maintained the tumors, and Group 3 (termed the Tumor removal group) underwent tumor removal surgery. We monitored the mice in Groups 2 and 3 until week 8 (when the tumor size reached the euthanasia criteria) for organ harvest. The luminescent signals of tumors increased in mice from the Control group but remained absent in mice from the Tumor removal group (**Figure 1B**).

To profile PCa cell dissemination, species-specific PCRs (α-satellite gene for humans and plakoglobin for mice) were performed. We used 2 µg or 600 ng DNA extracted from each dissected mouse tissue and semi-quantified the numbers of DTCs (**Figure 1C**). We detected DTCs at week 2 post-tumor cell injections in various organs examined, such as the liver, lung, and bones (**Figure 1D & Supplementary Figure S1**). The estimated DTC numbers were 1-10 cells/2 µg total DNA (approximately 0.001% to 0.01% of the total cells) in most organs (**Figure 1D & Supplementary Figure S1**). In mice from the Control group, an overall increasing trend of DTCs was observed in most of the organs examined at week 6 and week 8 compared to week 2, implying that the growing PCa tumors continue disseminating cancer cells into these organs. In mice of the Tumor removal group, fewer DTCs were detected compared to those in the Control group at the same week age and the respective organs in week 2, supporting the notion that tumor removal reduces the overall cancer burden and the major source of dissemination [26, 53]. However, in mice from the Tumor removal group, DTCs were only detected in bone and skeletal muscle, i.e., the tibiae, femurs, skull, spine, and skeletal muscles of the hind limbs at week 6. At week 8, DTCs were also detected in the prostate and kidney in some mice, but the most abundant DTCs were still detected in the tibiae and femurs (**Figure 1D & Supplementary Figure S1**). These data revealed that: 1) PCa cells disseminate to various organs when the tumors reach ∼100 mm^3^; 2) Along with the growth of subcutaneous tumors, DTCs increase in some organs such as the lung and skull but decrease in other organs such as the pancreas and liver; 3) After tumor removal, the DTCs reduced to an undetectable level in most organs, suggesting that the DTCs detected prior to the tumor removal were eliminated; 4) DTCs in these organs depend on the presence of a primary tumor and its continuing dissemination. However, DTCs were steadily detected in bones and skeletal muscles, and the amounts were maintained no more than those prior to tumor removal. These data suggest that bones are a protective reservoir, where DTCs are maintained but prevented from active proliferation.

To further delineate the localization of DTCs in bones, we separated the bone marrow (BM) from the bone cortex (BC) of the tibiae or femur via centrifugation (**Figure 1E**). DTCs were detected in the bone cortex at all the times examined but were undetectable in the bone marrow at week 2 and week 6 from mice of both the Control and Tumor removal groups. At week 8, DTCs were detected in the bone marrow of mice from the Control group but not the Tumor removal group (**Figure 1F**). The presence of intact DTCs in the bone marrow was confirmed through immunofluorescence (IF) staining of the human mitochondria marker (**Figure 1G**).

To further determine the dynamics of DTCs in the bones, we examined mice from the Tumor removal group at week 18 and were able to detect the bone marrow DTCs (**Figure 1F**). The bone marrow DTCs detected could either result from the migration of the bone cortex DTCs or from the proliferation of DTC arriving early in the bone marrow and maintaining at an undetectable level. Since DTCs and tumor mass found in the bone marrow in both patients and animal models correlate with overt bone metastases [54], bone marrow DTCs are possibly more proliferative than bone cortex DTCs. Therefore, we hypothesize that DTCs are dormant in the bone cortex before their proliferation and progression into the bone marrow.

### Osteoblasts induce PCa cell dormancy and physical contacts are required

To test our hypothesis and determine how PCa cells become dormant in the bone microenvironment, we mixed and co-cultured PCa C4-2B/GFP cells (denoted as C4-2B in the following co-culture context if not specified) with different stromal cells that compose the bone microenvironment and tracked C4-2B growth by monitoring GFP signals via IncuCyte imaging (**Figure 2A**). Our results showed that only osteoblasts (both mouse MC3T3-E1 cells and human hFOB1.19 cells tested) inhibited C4-2B cell proliferation (**Figure 2B and Supplementary Figure S3A**). In contrast, mesenchymal stem cell OP-9, osteoclast/macrophage RAW264.7, and fibroblast NIH3T3 showed no significant effect on C4-2B cell proliferation; osteocyte MLO-Y4 displayed a statistically significant stimulation but the effect size was neglectable; endothelia cell HUVEC showed a significant stimulation of C4-2B proliferation (**Figure 2B**). We found that C4-2B cells with high membrane dye intensity (indicating fewer cell divisions) accounted for at least 90% of total C4-2B cells in the mixed co-culture, compared to 20% in C4-2B cultured alone (**Figure 2C**). Using the PIP-FUCCI (PCNA-Interacting Protein degron version of Fluorescent Ubiquitination-based Cell Cycle Indicator), a dual fluorescence indicator designed for cell cycle phase labeling [34], we found that in the mixed co-culture, 70% of total C4-2B cells were in G0/G1 (green only) and 25% were in late S/G2/M (red), while in the C4-2B cultured alone, 25% of total C4-2B cells were in G0/G1 and 40% in late S/G2/M (**Figure 2D**), suggesting that MC3T3-E1 cells induced G0/G1 arrest. Furthermore, we found that MC3T3-E1 cells inhibited C4-2B cell apoptosis (**Figure 2E**) and had no effect on the expression of senescence markers such as p21 (*CDKN1A*) [55, 56] (**Figure 2F**). Similarly, the proliferation of other PCa cells, such as PC-3 or DU-145 cells, was also inhibited by MC3T3-E1 cells (**Supplementary Figure S2**). We then investigated whether physical contact between PCa cells and osteoblasts is essential for proliferation inhibition. Conditioned media (CM) from the mixed co-culture, CM from either C4-2B cells or MC3T3-E1 cells cultured alone, and CM from trans-well co-culture (C4-2B cells on top insert) were applied to C4-2B cells. We found that only CM from the mixed co-culture significantly inhibited C4-2B proliferation (**Figure 2G**), suggesting that physical contact is required; unknown secreted factors unique to the mixed co-culture can partially inhibit the C4-2B cell proliferation.

To further verify the dormancy status of the co-culture C4-2B cells, we examined the expression of a dormancy marker, NR2F1(Nuclear Receptor Subfamily 2 Group F Member 1)[19]. We found that, compared to C4-2B cultured alone, the mixed co-cultured C4-2B cells exhibited a significant increase in NR2F1 expression at both mRNA and protein levels (**Figure 2H, J**). Concurrently, we observed a decrease in the expression of Ki67 (MKI67) and cyclin D1 (CCND1) (**Figure 2H**). It is important to note that the gene expressions pertain to C4-2B cells, as we utilized human-specific primers for qPCR analysis of mRNA expression. The primer sequences are provided in **Table S2**, and we have included gels in the **Supplementary Data** to verify the species specificity of these primers. For the protein expressions, we successfully separated the co-cultured C4-2B cells (labeled with GFP) from osteoblasts (non-labeled). This separation was highly efficient, as supported by the fluorescence microscopy and genomic DNA PCR (shown in **Figure 2I** for MC3T3-E1 co-culture separation and **Supplementary Figure S3A** for hFOB1.19 co-culture separation).

To better mimic the bone microenvironment, we advanced our 2D mixed co-culture to 3D by embedding C4-2B and MC3T3-E1 cells in type I collagen, as type I collagen forms 90% of the organic mass of bone [57, 58]. We observed increased NR2F1 expression and decreased Ki67 and cyclin D1 expression in this 3D mixed co-culture compared to C4-2B cultured alone at the same 3D condition (**Figure 2K**), which suggests consistency between the 2D and 3D mixed co-cultures. For convenience, we continued to use the 2D mixed co-culture in our studies. These data suggested that osteoblasts induce C4-2B cells into a dormancy-like status through direct physical contact.

We also investigated the effects of cell number ratio and co-culture time on the dormancy induction in the mixed co-cultured of C4-2B and MC3T3-E1 cells. We observed concurrent increases of NR2F1 and decreases of Ki67 and Cyclin D1 at various C4-2B: MC3T3-E1 ratios ranging from 1:30 to 1:6 (**Supplementary Figure S4A**). Similar decreases of Ki67 and increases of NR2F1 were detected in co-cultures ranging from day 3 to day 6 (**Supplementary Figure S4B**). Increases of NR2F1 at the protein level were further confirmed from day 3 to day 6 at a ratio of 1:30 of the C4-2B: MC3T3-E1 cells (**Supplementary Figure S4C**).

To test whether the osteoblast-induced C4-2B dormancy is caused by nutrient deficiency because of the competition with the co-cultured osteoblasts, we cultured C4-2B cells in 1, 5, or 20% serum-containing media to compare with the C4-2B cells cultured in the standard 10% serum-containing media. We found that the proliferation of C4-2B cells was only inhibited by 1% serum-containing media among the groups (**Supplementary Figure S5A**); the NR2F1 mRNA expressions were modestly increased to 2.4 and 2.1 fold in 1% and 5% serum-containing media, respectively, but decreased to 1/2 in 20% serum-containing media; Ki67 and cyclin D1 were slightly decreased by about 1/4 in 1% and 20% serum-containing media, respectively (**Supplementary Figure S5B**). Together with the data that co-cultures with any other type of stromal cells could not inhibit C4-2B cell proliferation (**Figure 2B**), these data suggest that nutrient competition, such as reduced serum played little role, if any at all, in osteoblast-induced PCa cell dormancy. On the other hand, we also found that enzalutamide treatments, although inhibiting C4-2B proliferation (**Supplementary Figure S5C**), increased the expression of NR2F1 and decreased Ki67 but increased cyclin D1 (**Supplementary Figure S5D**). Altogether, these data showed uncoupled proliferation changes and dormancy markers, thus, suggesting unique key determinants to be explored for osteoblast-induced PCa cell dormancy.

### Osteoblasts decrease mitochondria-related genes in the mixed co-cultured C4-2B cells

To gain further insights into the molecular mechanisms for the PCa dormancy induced in the mixed co-culture, we conducted RNA-sequencing (RNA-Seq) to identify dysregulated genes compared to C4-2B cultured alone. We utilized three independent bioinformatic pipelines, Xenome [43], XenofilteR [44], and Kallisto [45], to isolate human-specific reads from the total reads of mixed human PCa cells and mouse osteoblasts. The common significantly dysregulated genes from the three pipelines were defined as dormant PCa signatures and further annotated for the top biological process enrichments using Gene Ontology (GO) (**Figure 3A**). The top 6 enriched biological processes from the significant down-regulated dysregulated genes were all mitochondria-related, including oxidative phosphorylation, energy-coupled proton transmembrane transport, mitochondrial ATP (adenosine triphosphate) synthesis coupled electron transport (**Figure 3B**). We observed the decreases of all 13 mitochondrial protein-coding genes (localized in the mitochondrial genome) at the mRNA level in the mixed co-cultured C4-2B cells in RNA-seq results and the decreases of 6 representative genes, i.e., MT-ATP6 (ATP synthase membrane subunit 6), MT-ND1,2&4 (NADH:Ubiquinone oxidoreductase core subunit), MT-CYB (cytochrome B), and MT-CO1 (cytochrome C oxidase I), were confirmed using qRT-PCR (**Figure 3C**). Additionally, we found that the mitochondrial DNA (mtDNA) copy number was downregulated in the mixed co-cultured C4-2B cells (**Figure 5D**). These findings suggest that decreased mitochondrial genes and suppression of mitochondria-related biological processes may serve as novel phenotypic dormant cell markers. Notably, these decreased mitochondrial genes were increased in the proliferation-inhibited C4-2B cells resulting from either serum limitation or AR antagonist enzalutamide **(Supplementary Figure S5B, D**), further supporting the unique dormant nature of C4-2B cells induced by the osteoblast co-culture.

**Figure 3.**
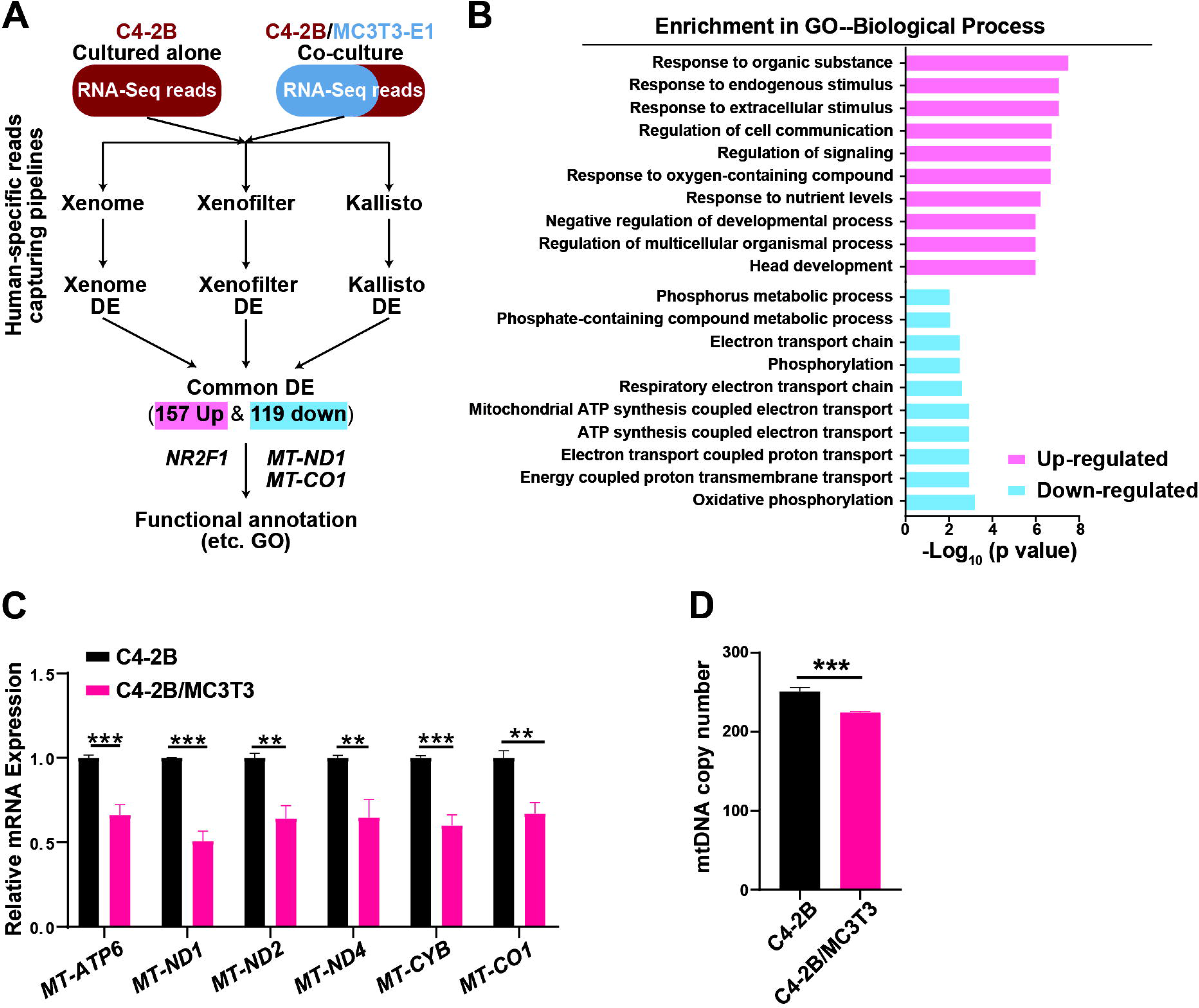
Mitochondrial gene expressions were decreased in dormant C4-2B cells. **A**. Workflow of RNA-sequencing analyses. **B**. Gene Ontology (GO) annotation of biological process in dormant C4-2B cells via ShinyGO v0.61. **C**. qRT-PCR confirmation of representative mitochondrial gene expression changes in the co-cultured C4-2B cells relative to the cells cultured alone. **D**. Analyses of mitochondrial DNA (mtDNA) copy numbers via qRT-PCR. C4-2B cells were cultured alone or co-cultured with MC3T3 cells for 72 h. All the experiments except for the RNA-Seq were independently repeated at least three times. The data of bar plots are presented as the mean ± SD of each set of triplicate samples. ** p < 0.01, *** p < 0.001.

**Figure 4.**
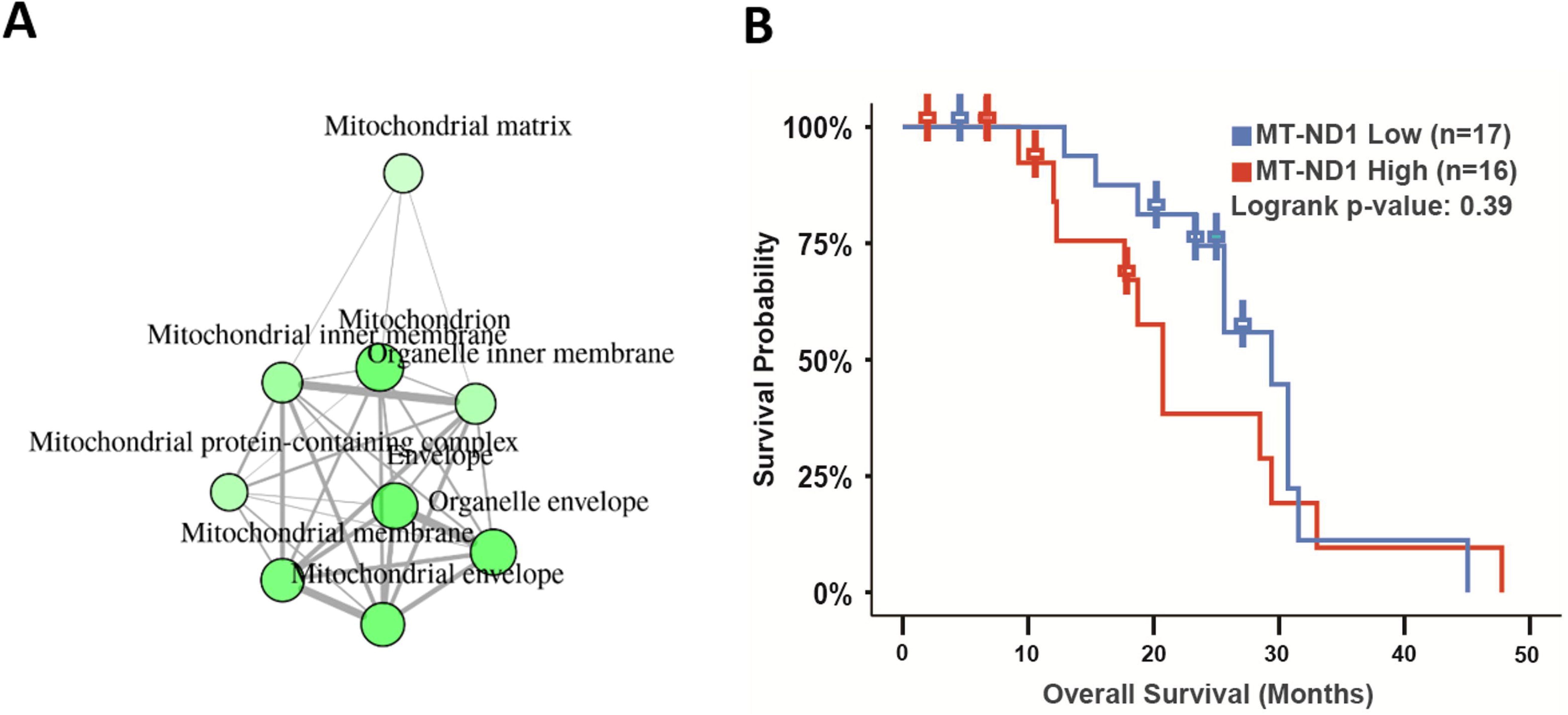
Mitochondrial gene elevation in circulating tumor cells and bone metastases of poor survival in PCa patients. **A.** Significantly up-regulated DE genes in circulating tumor cells from PCa patients were enriched to the mitochondrial structural components. Data were processed from the previously published study [59], and the plot was generated using ShinyGO v0.61. **B.** Patients with *MT-ND1* gene expression at a low level, compared to patients expressed *MT-ND1* at a high level, has a favorable prognostic trend in PCa patients with bone metastases (n=16 high vs n= 17 low, P= 0.39, from cBioportal study ID: prad_su2c_2019).

**Figure 5.**
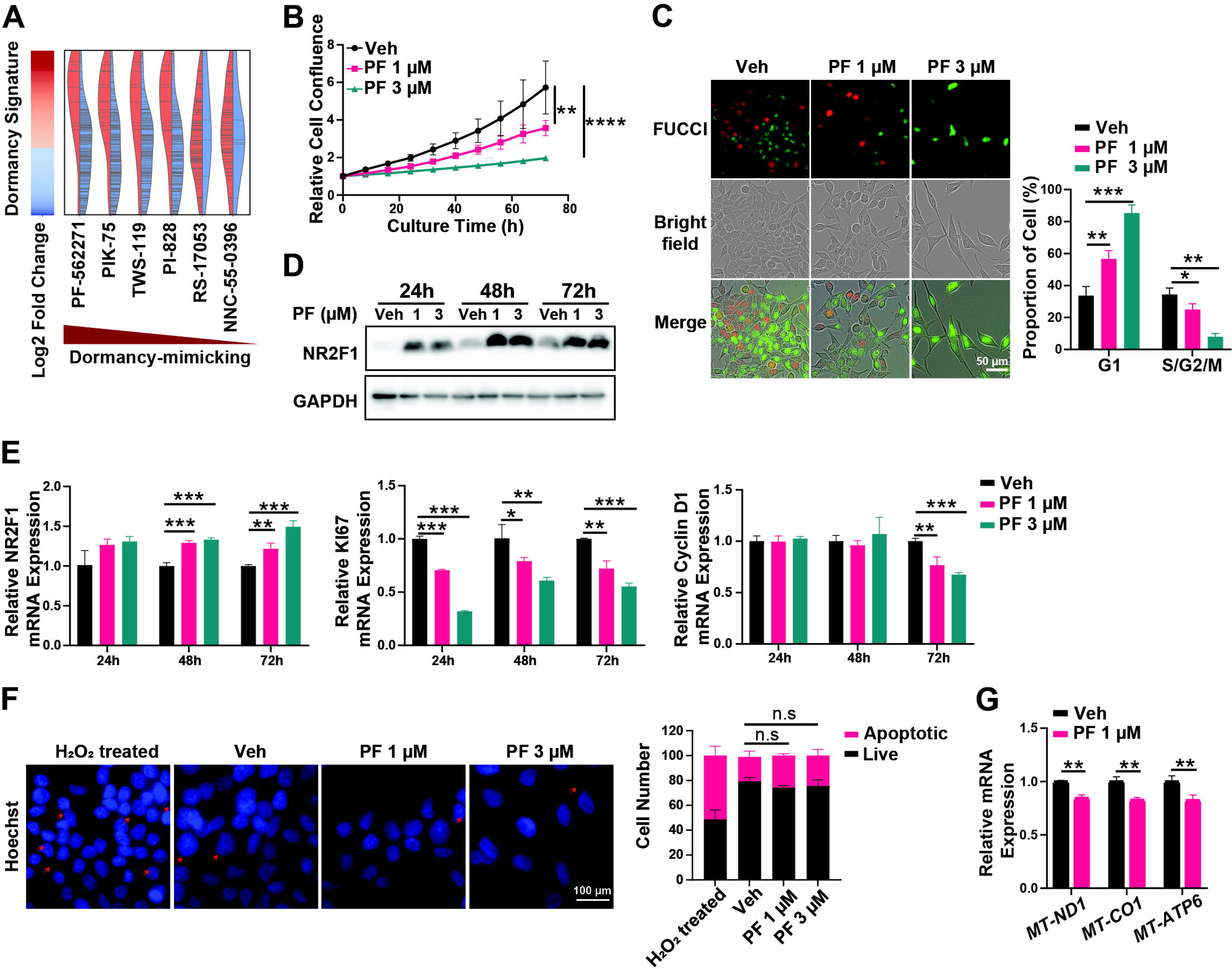
PF-271 treatments mimicked the osteoblast-induced dormancy in C4-2B cells. **A**. Top list of AI-predicted potential dormancy-mimicking drugs based on the dormant gene signature of C4-2B cells in the mixed co-culture. C4-2B cells were treated with 1 μM/ 3 μM of PF-271 (PF) or Vehicle (Veh, DMSO) for 72 h. Cell proliferation monitoring by Incucyte (**B**), cell cycle phase distribution by PIP-FUCCI (**C**), Time-lapse NR2F1 protein expressions by immunoblotting (**D**), and mRNA expressions of NR2F1/Ki67/Cyclin D1 (**E**) or MT-ND1/MT-CO1/MT-ATP6 (**G**) by qRT-PCR, were performed in the same manner as described in Figure 2. **F**. Analyses of apoptotic cells in C4-2B cells treated with 1 μM/ 3 μM of PF-271 (PF) or Vehicle (Veh, DMSO) for 72 h. H_2_O_2_ treatment for positive control. Scale bars, 50 μm (**C**) and 100 μm (**F**). All the in vitro experiments were independently repeated at least three times. The data in curves and bar plots are presented as the mean ± SD of each set of triplicate samples. * p < 0.05, ** p < 0.01, *** p < 0.001, **** p < 0.0001, and n.s. non-significant.

Furthermore, we analyzed published patient datasets and found that up-regulated dysregulated genes in circulating PCa tumor cells from patients are enriched to the mitochondrial structural components (data processed from a previously published study [59] (**Figure 4A**). Low MT-ND1 gene expression has a favorable prognosis in PCa patients with bone metastases (**Figure 4B**, from cBioportal study ID: prad_su2c_2019). The mining of these patient datasets suggests that reversing activated mitochondrial genes and enrichment of mitochondria-related biological processes, namely inducing PCa dormancy, could impede PCa metastases or progression.

To test whether mitochondrial inhibition could induce dormancy in C4-2B cells, we examined the effects of well-established mitochondrial inhibitors. We found that rotenone or IACS-010759 (IACS), both the respiration complex I inhibitors [60, 61], and FCCP, a potent uncoupler of oxidative phosphorylation in mitochondria [62], inhibited C4-2B proliferation in a dose-dependent manner (**Supplementary Figure S6A**). However, despite inhibiting C4-2B proliferation, various doses of rotenone and IACS significantly decreased the expression of dormancy marker NR2F1 at 24 to 72 hours (**Supplementary Figure S6B)**. At 72 hours, FCCP also decreased NR2F1 and increased the expressions of Ki67, Cyclin D1, MT-ND1, and MT-CO1 (**Supplementary Figure S6C**). These data suggested that direct mitochondrial inhibition could not induce PCa cell dormancy.

Altogether, our data demonstrated that osteoblasts induced C4-2B cells into dormancy, defined with known dormancy features such as increased NR2F1 and decreased Ki67 and Cyclin D1 mRNA expressions. Importantly, we identified a novel dormant PCa cell feature, mitochondria-related gene suppression, possibly reversed in CTCs, and the increased mitochondrial genes could serve as poor survival markers. However, mitochondrial inhibitors could not recapitulate the dormancy-like phenotype, suggesting the enrichment of the decreased mitochondria-related biological processes is likely a result, rather than a cause, of the osteoblast-induced PCa cell dormancy.

### PF-271 mimics the osteoblast-induced C4-2B dormancy

To investigate how to induce dormancy in C4-2B cells, we utilized a novel AI platform (developed and currently unpublished by Dr. Bin Chen and colleagues) to predict drugs that mimic the dormant PCa cell gene signature. This approach allowed us to compare the dormancy gene signature with fold changes against a database of drug-induced gene expression matrices and score drugs based on the similarities between drug-induced gene expression profiles and the dormant signature (**Figure 5A**). Among the top mimicking drugs, PF-562271 (PF-271) and PIK-75 were the top 2 drugs with better prediction scores and growth inhibition potency tested in C4-2B cells (**Supplementary Figure S7**).

PF-271 is a reversible and ATP-competitive inhibitor of focal adhesion kinase (FAK) [31–33]. We showed that PF-271 inhibited cell C4-2B proliferation in a dose-dependent manner (**Figure 5B**), induced G0/G1 arrest (**Figure 5C**), increased NR2F1 expression at both mRNA and protein levels, decreased Ki67 and cyclin D1 mRNA expression (**Figure 5D&E**), but had no effect on apoptosis (**Figure 5F**). PF-271 also decreased the mRNA expressions of the mitochondrial genes, for example, MT-ND1, MT-CO1, and MT-ATP6 (**Figure 5G**). Taken together, these results suggest that PF-271 mimics the osteoblast-induced C4-2B cell dormancy.

In contrast, PIK-75, currently known as an inhibitor of the p110α PI3K[63], inhibited C4-2B cell proliferation in a dose-dependent manner but did not induce C4-2B dormancy (**Supplementary Figure S8**). PIK-75 did not affect cell cycle distribution (**Supplementary Figure S8B**) or NR2F1 protein expression at 48 and 72 hours (**Supplementary Figure S8C**). Although PIK-75 did induce the expression of NR2F1 at the mRNA level at 48 and 72 hours, it increased the expressions of Ki67 and Cyclin D1 (**Supplementary Figure S8D**), and induced apoptosis (**Supplementary Figure S8E**). Altogether, these data indicate that PIK-75 does not have a dormancy-mimicking effect.

### PF-271 inhibits FAK phosphorylation and promotes FAK nuclear translocation in C4-2B cells

To investigate how PF-271 induced the dormancy-mimicking effect, we first tested its known effect on FAK and included two other FAK inhibitors, Y15 and defactinib [64, 65]. We observed growth inhibitions and NR2F1 protein accumulation in C4-2B cells treated with Y15 or defactinib (**Supplementary Figure S9**), suggesting FAK is an essential mediator in this dormancy induction. In both C4-2B and PC-3 cells, PF-271 treatment resulted in significant decreases in phosphorylated FAK at Y397 and Cyclin D1 (**Figure 6A** & **Supplementary Figure S10**). We further observed PF-271 resulted in an increase of nuclear FAK in C4-2B/GFP-FAK cells (overexpressing GFP-tagged FAK) (**Figure 6B & Supplementary Movie S1, S2**). Notably, compared with the C4-2B/GFP-FAK cells cultured alone or co-cultured with OP-9 mesenchymal stem cells, C4-2B/GFP-FAK cells co-cultured with MC3T3-E1 cells had decreased phospho-FAK at Y397 and increased nuclear translocation of GFP-FAK, similar to PF-271 treated C4-2B cells (**Figure 6C**, **D & Supplementary Movies S3-6**). These data suggest that PF-271 treatment mimics osteoblast-induced C4-2B dormancy by inhibiting FAK phosphorylation and promoting FAK nuclear translocation and that FAK is a key mediator of PCa dormancy.

**Figure 6.**
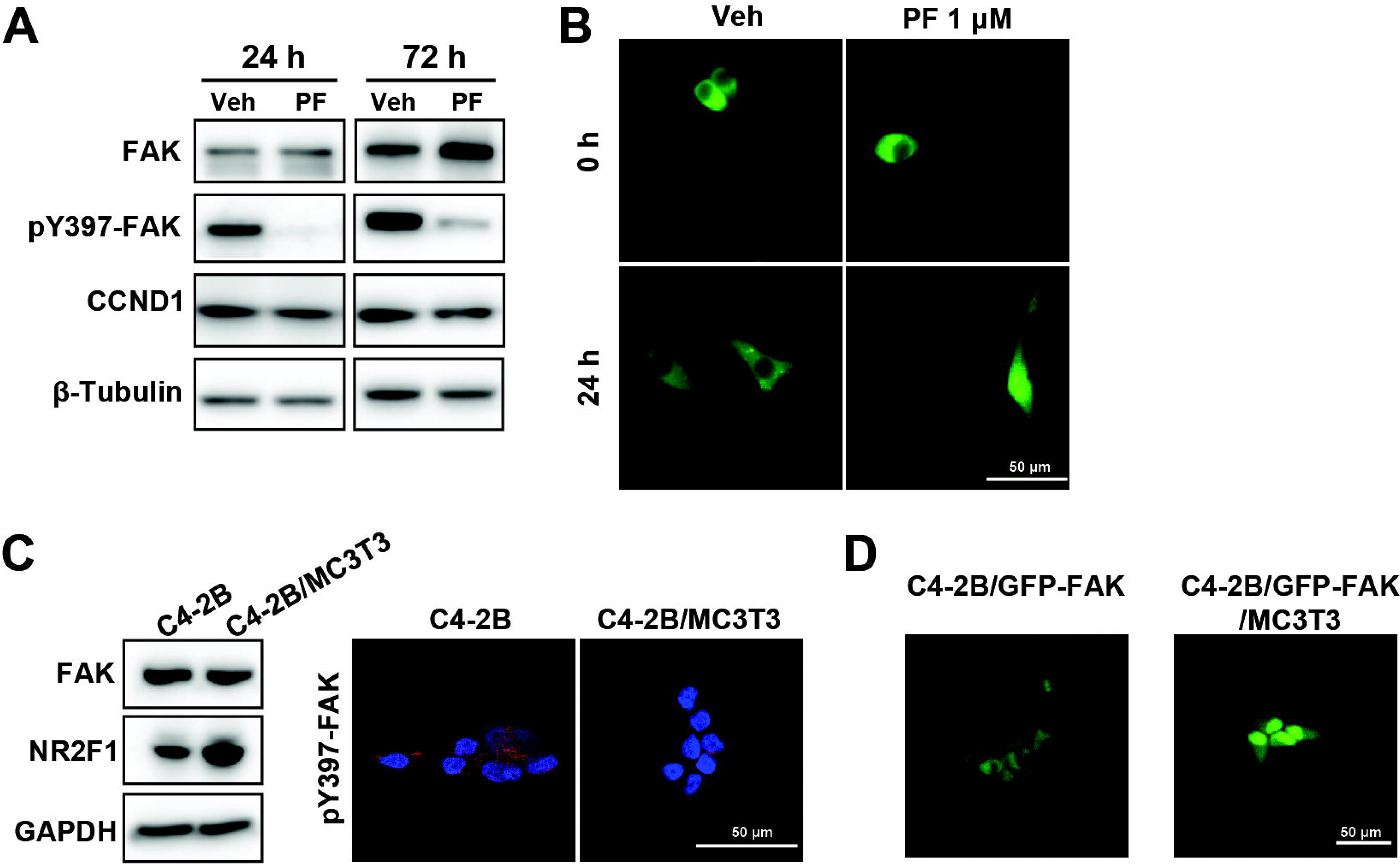
PF-271 mimicked C4-2B dormancy by blocking FAK phosphorylation and promoting FAK nuclear translocation. **A**. C4-2B/GFP-FAK cells were treated with 1 μM PF or Veh (Vehicle, DMSO) for 24 h or 72 h. The protein levels of pY397-FAK, total FAK, and CCND1 (cyclin D1) were examined by immunoblotting. **B**. C4-2B/GFP-FAK cells were treated with 1 μM PF or Veh (Vehicle, DMSO) for 24 h and monitored every 2h on IncuCyte S3. Representative pictures at 24 h are shown. **C**. C4-2B cells were co-cultured with MC3T3-E1 or cultured alone for 72 h. The protein levels of total FAK and NR2F1 were examined by immunoblotting (left). Phospho-FAK levels were examined with immunostaining and captured with confocal microscopy. Red, pY397-FAK; blue, DAPI, nucleus. **D**. C4-2B/GFP-FAK cells were co-cultured with MC3T3-E1 or cultured alone for 72 h. Representative pictures at 72 h are shown. Scale bars, 50 μm. All the experiments were independently repeated at least three times.

## Discussion

Metastatic progression and recurrence, despite curative intended treatments, are among the leading causes of cancer-related mortality [66, 67]. PCa cases increased by nearly 300 thousand annually in the US and more than 1.4 million worldwide [68]. Early dissemination, dormancy, and reactivation of cancer cells have been implicated as the key factors driving these events. However, the impact of clinical treatments on cancer dissemination and dormancy, as well as the underlying mechanisms of dormancy, remain poorly understood [69–71]. In this study, we investigated the effects of primary tumor removal on PCa cell dissemination in a mouse xenograft model. We found that tumor removal decreased DTC numbers to levels below the detection limits in all the tissues examined, except for the bones. Furthermore, we observed a dynamic progression of detectable DTCs from the bone cortex to the bone marrow. The *in vitro* experiments revealed that osteoblasts induced PCa cell dormancy. Moreover, we identified a novel and unique gene signature in the osteoblast-induced dormant PCa cells, defined by the increased expression of dormancy markers, suppression of proliferation, and decreased mitochondrial protein-coding genes. Using an AI-facilitated drug repurposing approach, we discovered and validated the clinical drug PF-271 mimicking the osteoblast-induced C4-2B cell dormancy by inhibiting FAK phosphorylation and increasing FAK nuclear translocation. Therefore, our data suggest that FAK is a key mediator of PCa dormancy and a promising target for inhibiting and preventing PCa metastatic recurrence. The summary and hypothesis of our study is illustrated in **Figure 7**.

**Figure 7.**
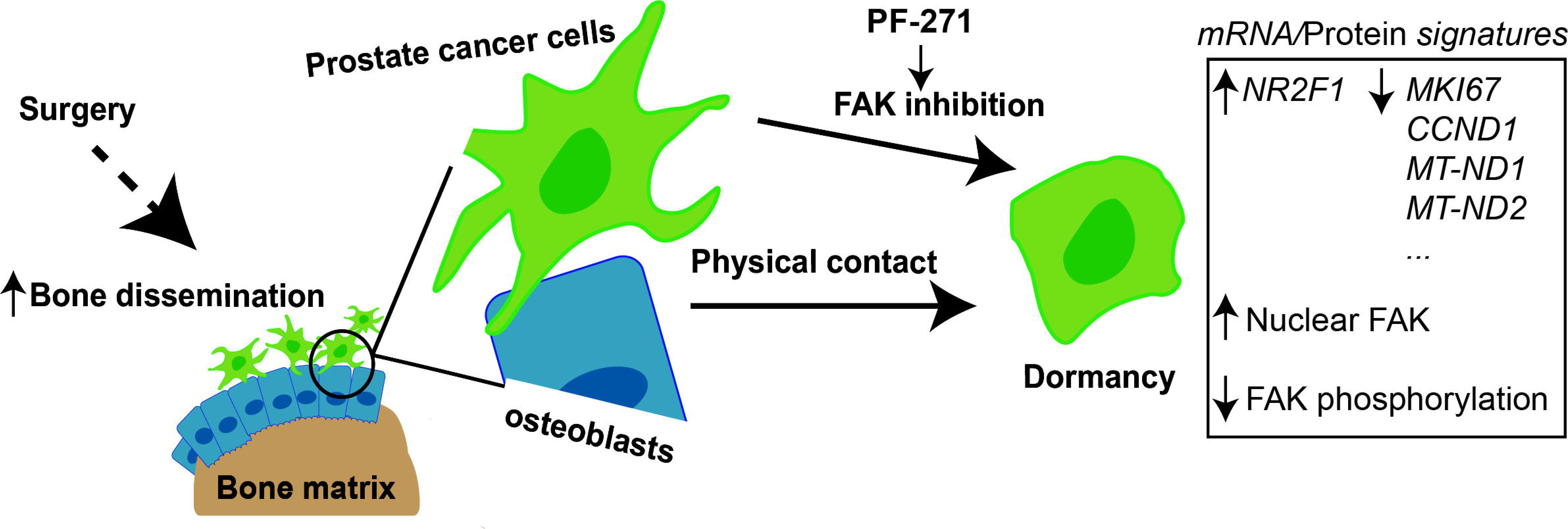
The summary and hypothesis. Physical contact between osteoblasts and cancer cells is required for the dormancy induction by inactivating FAK, which enhances its nuclear translocation and reduces its phosphorylation. Dormant prostate cancer cells exhibit increased *NR2F1* expression, decreased expression of *MKI67* and *CCND1*, and reduced levels of OXPHOS-related genes such as *MT-ND1 and MT-ND2*. PF-271, a small molecule inhibitor of FAK, can induce dormancy in the prostate cancer cells.

Correlations between surgery and cancer progression have been observed and debated in clinical and animal models [26]. Generally, primary tumor resection contributed to longer survival, slower PSA increase, and lower metastatic risks, reported in two patient cohorts and *in vivo* studies [72, 73]. However, a non-negligible 25% of biochemical recurrence and a further 10% of metastases were recently reported among patients receiving prostatectomy [74]. To our knowledge, the DTC quantification in mouse xenografts conducted in this study is the first longitudinal examination of organ-specific PCa cell dissemination. DTCs were detected in various organs, such as bone, liver, lung, and kidney, consistent with the clinically observed sites for PCa metastases [3]. Interestingly, our data revealed that with the presence and growth of primary tumors, DTCs increase in some organs but decrease in others. Whether this discrepancy has true clinical relevance or is merely the variance of individual mice needs to be further determined. The primary tumor is likely exerting an influence on the dissemination of DTCs in the bone marrow. It could either continue to spread to various organs or trigger the activation of DTCs within the bone cortex, leading to their migration into the bone marrow. Another possibility is that the detection of DTCs in the bone marrow may have resulted from their proliferation from undetectable levels prior to week 8.

After the tumor removal, DTCs were reduced to undetectable levels in most organs. However, DTCs were detectable in bones and skeletal muscles, and the amounts were lower than before tumor removal, suggesting bones are the protecting reservoir for DTCs to maintain dormant. Furthermore, DTCs were detected in the bone cortex prior to the bone marrow, implying that surgical removal of primary tumor prolonged PCa DTCs in the bone cortex. The mechanisms underlying the detection of DTCs in the bone marrow at week 18 after the removal of the primary tumor are intriguing and will require further elucidation in future studies. During this period, there was no input from the primary tumor for 16 weeks, spanning from week 2 to week 18. We hypothesize that the source of these marrow DTCs could either be DTCs from the bone cortex that proliferated and migrated into the bone marrow, or DTCs that were actively proliferating within the bone marrow itself.

Clinically, metastatic PCa tumors were found in the marrows of bone metastatic patients and the presence of DTCs in the marrow was associated with poor prognosis [9–13, 70, 75, 76]. Since bone marrow DTCs were only identified until later stages, i.e., week 8 in the Control and week 18 in the Tumor removal group, we hypothesize that DTCs are dormant in the bone cortex. The semi-quantitative PCR method that we have optimized provides a cost-effective approach and accessible platform to evaluate the trends of PCa cell dissemination and relapse. It will be interesting to apply this method to investigate how other therapies, such as chemotherapy, first-line androgen-deprivation therapy drugs, or androgen receptor antagonist enzalutamide, affect PCa cell dissemination and dormancy.

Signatures of dormant cancer cells, including PCa, have been reported in previous studies but few of these studies characterized dormant PCa cells in the bone microenvironment [22, 25, 40, 77–80]. Our study contributed to PCa dormant signature in the context of osteoblasts, which shared some common markers genes with previous studies but highlighted some unique changes. Consistently, NR2F1 was shown as a marker for dormant cancer cells [27, 81–83]. Stem cell markers ALDH1A1 and BCL11B showed significant up-regulation in our dormant C4-2B DE gene list **(Supplementary Table S5)**, but BMI1 and SOX2 were not changed [84, 85]. SOX9 increase was reported as a dormant head and neck squamous cancer cell marker [27]. However, SOX9 expression was significantly decreased in our osteoblast-induced dormant C4-2B cells (**Supplementary Figure S11A & Table S5**). Consistently, SOX9 overexpression promotes the development of invasive carcinoma [86, 87], suggesting a distinct role of SOX9 in prostate cancer. An increased ratio of the phospho-p38 to the phospho-ERK was described as a dormancy marker [5]. In our system, we noticed a decreased ratio of phospho-p38 to the phospho-ERK in co-cultured dormant C4-2B cells (**Supplementary Figure S11B).** AXL receptor tyrosine kinase was reported for inducing AR-negative PCa cells PC-3 and DU-145 in the MC3T3-E1 co-culture [25], but AR-positive C4-2B cells do not express AXL proteins although we observed an increase of AXL expression in the RNA-Seq. The unique gene signature of dormant PCa cells was possibly a result of response to different tumor microenvironments. Specifically, osteoblasts might dictate the signatures specific to bone metastatic dissemination, dormancy, progression, and relapse. The mitochondrial functional enrichment was identified in the down-regulated DE of the dormant C4-2B gene signature. In contrast, in a recent publication, mitochondrial functional enrichment in up-regulated DE was found in PCa metastases compared to primary tumors [88]. These data support and validate the clinical significance of the dormant C4-2B signature, i.e., the reversal of the bone metastatic progression and relapse. The signature of decreased mitochondrial gene expressions along with increased NR2F1 and decreased Ki67/Cyclin D1 helped distinguish PCa cells of bona fide dormancy from proliferation inhibition by serum starvations, enzalutamide treatment, or PIK-75 treatment. Therefore, this signature can be used to discover drugs that could mimic or reverse PCa cell dormancy, could treat or prevent bone metastasis and relapse, as well as to understand the basic biology of PCa cells. Because the dormant PCa cells cause no symptoms in patients, maintaining cancer cell dormancy could be an operational cure for cancer patients.

Using both 2D and 3D *in vitro* co-culture of PCa and various types of cells of the bone microenvironment, we found that osteoblasts’ direct physical contact with PCa cells is necessary and sufficient to induce PCa cells into a dormancy-like status. This is consistent with previous reports using different approaches [80, 89]. Furthermore, the reverse correlation between the osteoblast-induced PCa dormancy signature with the clinical bone metastatic and CTC signatures supports that the direct co-culture of PCa cells and osteoblasts is an easy and powerful platform for rapid screening of drug candidates that can modulate PCa dormancy. Co-culture of C4-2B and MC3T3-E1 without direct contact via transwell resulted in a modest increase in NR2F1 but no changes in Ki67 and cyclin D1 mRNA expressions (**Supplementary Figure S2F**). The conditioned media (CM) from directly mixed co-culture induced partial growth inhibition of C4-2B cells but CM from non-contact transwell co-culture did not affect C4-2B growth at all, suggesting that the secreted factors stimulated by physical contact co-culture and signaling at the C4-2B/MC3T3-E1 interface were responsible for dormancy induction. Both need to be determined by further studies.

Using a novel AI tool (unpublished), we predicted the dormancy-mimicking drugs. The *in vitro* testing suggested the dormancy-mimicking effect of PF-271, an FAK inhibitor. Coincidently, using the human epidermoid carcinoma HEp3 cells that were passaged 120 to 170 times in chicken embryos as a dormancy model, a previous study found that genetic inhibition of FAK could mimic the tumor dormancy induced by the specific microenvironment, in which FAK was activated by the urokinase plasminogen activator receptor (uPAR) and mediating the expression of ERK [90]. In our osteoblast-induced dormant PCa cells, neither uPAR nor ERK was changed, suggesting a role of FAK in PCa dormancy with a distinct mechanism. Inhibition of FAK was also reported to be sufficient to prevent tight junction disruption [91], validating that cell-cell interaction such as adhesion and tight junction play a vital role in establishing PCa dormancy. We noticed the functional enrichment of KEGG pathway “cell adhesion molecules” in up-regulated DE in dormant cells (**Supplementary Table S6**). We revealed, for the first time in C4-2B cells, that PF-271 blocked FAK phosphorylation and increased FAK nuclear translocation, through which increased NR2F1 expression and inhibited cyclin D1 expression, in turn, induced cancer cell dormancy. The efficacy of PF-271 *in vivo* models of metastases and relapse will be tested in future studies.

We recognized the limitations of our study. Co-culture of relatively homogeneous PCa cells with osteoblasts over-simplified the complicated bone microenvironment. Other regulators such as mechanical stresses constantly received and changed in the bone microenvironment may also impact the tumor dormancy. Although C4-2B and PC-3 displayed the same dormancy phenotype in proliferation inhibition and molecular markers (increased NR2F1 and decreased Ki67/Cyclin D1), we also found that DU-145 displayed proliferation inhibition but NR2F1 was decreased; another PCa cell line, 22Rv1, did not show proliferation inhibition although NR2F1 was modestly increased (**Supplementary Figure S2E**). These data suggested that certain PCa cell intrinsic features play roles in responding to osteoblasts and need to be further defined. For example, DU-145 is a brain metastatic cell line and 22Rv1 is featured with the elevated expression of AR variant ARv7. We haven’t been able to show the presence of DTCs in the bone cortex other than PCR. Bone CLARITY may be a better approach to visualize the localization of the DTCs *ex vivo* [92]. We also wondered whether the detected DTC are CTCs trapped in harvested organs. However, the DTCs in mice of the tumor removal group clearly showed a preference for DTC in the bone over the liver, one of the blood-rich organs. No positive human cells were detectable by PCR in the peripheral blood of tumor-bearing mice (**Supplementary Figure S1D**). Together, these data suggest that the detected DTC in mice without perfusion is unlikely CTCs and the bone preference of dissemination is consistent with the frequently observed bone-tropism of PCa [93].

## Conclusions

Our study shed light on the effect of tumor removal on PCa cell dissemination with clinically relevant focus on the bones. We determined a unique osteoblast-induced PCa cell dormancy signature with significant suppression of mitochondrial gene expression, which was reported to be activated in bone metastases of PCa. The reverse correlation of dormant PCa cells and overt metastases in the bone microenvironment supported the rationale that inducing PCa dormancy prevents bone metastasis. Therefore, through an AI-powered algorithm, we identified a dormancy-mimicking drug, which induces PCa cell dormancy via FAK inhibition. Next, we will test and optimize the FAK inhibitors on dormancy induction and DTC suppression in preclinical mouse models. We will also address the molecular mechanisms by which osteoblasts induce PCa cell dormancy in the near future.

## Supporting information

Supplementary Figures and raw pictures for western blotting and agarose gel

## Abbreviation

AI: Artificial Intelligence;
ALDH1A1: Aldehyde Dehydrogenase 1 Family Member A1;
AR: Androgen Receptor;
AXL: AXL Receptor Tyrosine Kinase;
BC: Bone Cortex;
BCL11B: BCL11 Transcription Factor B;
BM: Bone Marrow;
CM: Conditioned Media;
CTC: Circulating Tumor Cell;
DE: Differentially Expressed genes;
DTC: Disseminated Tumor Cell;
ERK: extracellular signal-regulated kinase;
FAK: Focal Adhesion Kinase;
FUCCI: Fluorescent Ubiquitination-based Cell Cycle Indicator;
GFP: Green Fluorescence Protein;
GO: Gene Ontology;
KEGG: Kyoto Encyclopedia of Genes and Genomes;
MT-ATP6: Mitochondrially Encoded ATP Synthase Membrane Subunit 6;
MT-ND1/2/4: Mitochondrially Encoded NADH:Ubiquinone Oxidoreductase Core Subunit 1/2/4;
MT-CO1: Mitochondrially Encoded Cytochrome C Oxidase I;
MT-CYB: Mitochondrially Encoded Cytochrome B;
mtDNA: mitochondrial DNA;
NR2F1: Nuclear Receptor Subfamily 2 Group F Member 1;
PCa: Prostate Cancer;
PIP: PCNA-Interacting Protein;
PSA: Prostate-Specific Antigen;
qRT-PCR: quantitative real-time polymerase chain reaction;
RNA-Seq: RNA sequencing;
SOX2: SRY-Box Transcription Factor 2;
SOX9: SRY-Box Transcription Factor 9;
uPAR: urokinase plasminogen activator receptor;

## Declarations

### Ethics approval and consent to participate

All animal experiments were approved by the Institutional Animal Care and Use Committee (IACUC) under protocol numbers 400066/400131 at the University of Toledo. No human subjects were involved.

### Consent for publication

Not applicable

### Availability of data and materials

The datasets generated, i.e., raw RNA-Seq read files (FastQ) for C4-2B cultured alone and C4-2B/MC3T3-E1 mixed co-culture are available in the GEO repository under accession GSE210751. The sources for the public or previously published RNA-Seq files processed in this manuscript are included in the context. Heatmaps for DTC were generated using the “pheatmap” package in R. All codes on RNA-Seq analyses and heatmap drawings are available from the corresponding author on reasonable request.

### Authors’ contributions

XL conceived and directed the project. XL, RL, and SS wrote the manuscript. XL, RL, and SS designed and performed the experiments with help from YZ, MS, and CMG. JX performed the AI drug prediction. SS and KL performed the bioinformatic analyses under the support and guidance of BC. RL and SS performed the statistical analyses. DJ and EYA developed the 3D co-culture model and DJ and RL performed the related experiments. All authors have read and agreed to the final version of the manuscript.

## Acknowledgments

We thank Dr. Songpeng Zu from the University of California, San Diego, for his advice on statistical analyses and Mr. Shubhra Kanti Dey and Mr. Augustine Kwabil from the University of Toledo for technical support.

## References

1. Siegel RL, Miller KD, Wagle NS, Jemal A: Cancer statistics, 2023. CA Cancer J Clin 2023, 73:17-48.

2. Berruti A, Dogliotti L, Bitossi R, Fasolis G, Gorzegno G, Bellina M, Torta M, Porpiglia F, Fontana D, Angeli A: Incidence of skeletal complications in patients with bone metastatic prostate cancer and hormone refractory disease: predictive role of bone resorption and formation markers evaluated at baseline. J Urol 2000, 164:1248–1253.

3. Bubendorf L, Schopfer A, Wagner U, Sauter G, Moch H, Willi N, Gasser TC, Mihatsch MJ: Metastatic patterns of prostate cancer: an autopsy study of 1,589 patients. Hum Pathol 2000, 31:578–583.

4. Sartor O, de Bono JS: Metastatic Prostate Cancer. N Engl J Med 2018, 378:1653–1654.

5. Aguirre-Ghiso JA: Models, mechanisms and clinical evidence for cancer dormancy. Nat Rev Cancer 2007, 7:834–846.

6. Phan TG, Croucher PI: The dormant cancer cell life cycle. Nat Rev Cancer 2020, 20:398–411.

7. Ruppender NS, Morrissey C, Lange PH, Vessella RL: Dormancy in solid tumors: implications for prostate cancer. Cancer Metastasis Rev 2013, 32:501–509.

8. Uhr JW, Pantel K: Controversies in clinical cancer dormancy. Proc Natl Acad Sci U S A 2011, 108:12396–12400.

9. Pfitzenmaier J, Ellis WJ, Arfman EW, Hawley S, McLaughlin PO, Lange PH, Vessella RL: Telomerase activity in disseminated prostate cancer cells. BJU Int 2006, 97:1309–1313.

10. Thomas C, Wiesner C, Melchior SW, Schmidt F, Gillitzer R, Thuroff JW, Pfitzenmaier J: Urokinase-plasminogen-activator receptor expression in disseminated tumour cells in the bone marrow and peripheral blood of patients with clinically localized prostate cancer. BJU Int 2009, 104:29–34.

11. Murray NP: Minimal residual disease in prostate cancer patients after primary treatment: theoretical considerations, evidence and possible use in clinical management. Biol Res 2018, 51:32.

12. Murray NP, Aedo S, Fuentealba C, Reyes E, Salazar A: Minimum Residual Disease in Patients Post Radical Prostatectomy for Prostate Cancer: Theoretical Considerations, Clinical Implications and Treatment Outcome. Asian Pac J Cancer Prev 2018, 19:229–236.

13. Murray NP, Aedo S, Fuentealba C, Reyes E, Salazar A, Lopez MA, Minzer S, Orrego S, Guzman E: Minimal Residual Disease Defines the Risk and Time to Biochemical Failure in Patients with Pt2 and Pt3a Prostate Cancer Treated With Radical Prostatectomy: An Observational Prospective Study. Urol J 2020, 17:262–270.

14. Murray NP, Fuentealba C, Reyes E, Salazar A, Guzman E, Orrego S: The Epstein criteria predict for organ-confined prostate cancer but not for minimal residual disease and outcome after radical prostatectomy. Turk J Urol 2020, 46:360–366.

15. Aguirre-Ghiso JA: How dormant cancer persists and reawakens. Science 2018, 361:1314–1315.

16. Datta SR, Dudek H, Tao X, Masters S, Fu H, Gotoh Y, Greenberg ME: Akt phosphorylation of BAD couples survival signals to the cell-intrinsic death machinery. Cell 1997, 91:231–241.

17. Abida W, Cyrta J, Heller G, Prandi D, Armenia J, Coleman I, Cieslik M, Benelli M, Robinson D, Van Allen EM, et al: Genomic correlates of clinical outcome in advanced prostate cancer. Proc Natl Acad Sci U S A 2019, 116:11428–11436.

18. Giancotti FG: Mechanisms governing metastatic dormancy and reactivation. Cell 2013, 155:750–764.

19. Sosa MS, Bragado P, Aguirre-Ghiso JA: Mechanisms of disseminated cancer cell dormancy: an awakening field. Nat Rev Cancer 2014, 14:611–622.

20. Albrengues J, Shields MA, Ng D, Park CG, Ambrico A, Poindexter ME, Upadhyay P, Uyeminami DL, Pommier A, Kuttner V, et al: Neutrophil extracellular traps produced during inflammation awaken dormant cancer cells in mice. Science 2018, 361.

21. Kurppa KJ, Liu Y, To C, Zhang T, Fan M, Vajdi A, Knelson EH, Xie Y, Lim K, Cejas P, et al: Treatment-Induced Tumor Dormancy through YAP-Mediated Transcriptional Reprogramming of the Apoptotic Pathway. Cancer Cell 2020, 37:104–122 e112.

22. Owen KL, Gearing LJ, Zanker DJ, Brockwell NK, Khoo WH, Roden DL, Cmero M, Mangiola S, Hong MK, Spurling AJ, et al: Prostate cancer cell-intrinsic interferon signaling regulates dormancy and metastatic outgrowth in bone. EMBO Rep 2020, 21:e50162.

23. Vera-Ramirez L: Cell-intrinsic survival signals. The role of autophagy in metastatic dissemination and tumor cell dormancy. Semin Cancer Biol 2020, 60:28–40.

24. Jung Y, Cackowski FC, Yumoto K, Decker AM, Wang Y, Hotchkin M, Lee E, Buttitta L, Taichman RS: Abscisic acid regulates dormancy of prostate cancer disseminated tumor cells in the bone marrow. Neoplasia 2021, 23:102–111.

25. Yumoto K, Eber MR, Wang J, Cackowski FC, Decker AM, Lee E, Nobre AR, Aguirre-Ghiso JA, Jung Y, Taichman RS: Axl is required for TGF-beta2-induced dormancy of prostate cancer cells in the bone marrow. Sci Rep 2016, 6:36520.

26. Demicheli R, Retsky MW, Hrushesky WJ, Baum M, Gukas ID: The effects of surgery on tumor growth: a century of investigations. Ann Oncol 2008, 19:1821–1828.

27. Sosa MS, Parikh F, Maia AG, Estrada Y, Bosch A, Bragado P, Ekpin E, George A, Zheng Y, Lam HM, et al: NR2F1 controls tumour cell dormancy via SOX9- and RARbeta-driven quiescence programmes. Nat Commun 2015, 6:6170.

28. Sharma S, Pei X, Xing F, Wu SY, Wu K, Tyagi A, Zhao D, Deshpande R, Ruiz MG, Singh R, et al: Regucalcin promotes dormancy of prostate cancer. Oncogene 2021, 40:1012–1026.

29. Murray NP, Reyes E, Tapia P, Badinez L, Orellana N, Fuentealba C, Olivares R, Porcell J, Duenas R: Redefining micrometastasis in prostate cancer - a comparison of circulating prostate cells, bone marrow disseminated tumor cells and micrometastasis: Implications in determining local or systemic treatment for biochemical failure after radical prostatectomy. Int J Mol Med 2012, 30:896–904.

30. Cackowski FC, Eber MR, Rhee J, Decker AM, Yumoto K, Berry JE, Lee E, Shiozawa Y, Jung Y, Aguirre-Ghiso JA, Taichman RS: Mer Tyrosine Kinase Regulates Disseminated Prostate Cancer Cellular Dormancy. J Cell Biochem 2017, 118:891–902.

31. Sun H, Pisle S, Gardner ER, Figg WD: Bioluminescent imaging study: FAK inhibitor, PF-562,271, preclinical study in PC3M-luc-C6 local implant and metastasis xenograft models. Cancer Biol Ther 2010, 10:38–43.

32. Bagi CM, Roberts GW, Andresen CJ: Dual focal adhesion kinase/Pyk2 inhibitor has positive effects on bone tumors: implications for bone metastases. Cancer 2008, 112:2313–2321.

33. Roberts WG, Ung E, Whalen P, Cooper B, Hulford C, Autry C, Richter D, Emerson E, Lin J, Kath J, et al: Antitumor activity and pharmacology of a selective focal adhesion kinase inhibitor, PF-562,271. Cancer Res 2008, 68:1935–1944.

34. Grant GD, Kedziora KM, Limas JC, Cook JG, Purvis JE: Accurate delineation of cell cycle phase transitions in living cells with PIP-FUCCI. Cell Cycle 2018, 17:2496–2516.

35. Becker M, Nitsche A, Neumann C, Aumann J, Junghahn I, Fichtner I: Sensitive PCR method for the detection and real-time quantification of human cells in xenotransplantation systems. Br J Cancer 2002, 87:1328–1335.

36. Baylan N, Bhat S, Ditto M, Lawrence JG, Lecka-Czernik B, Yildirim-Ayan E: Polycaprolactone nanofiber interspersed collagen type-I scaffold for bone regeneration: a unique injectable osteogenic scaffold. Biomed Mater 2013, 8:045011.

37. Subramanian G, Elsaadany M, Bialorucki C, Yildirim-Ayan E: Creating homogenous strain distribution within 3D cell-encapsulated constructs using a simple and cost-effective uniaxial tensile bioreactor: Design and validation study. Biotechnol Bioeng 2017, 114:1878–1887.

38. Elsaadany M, Harris M, Yildirim-Ayan E: Design and Validation of Equiaxial Mechanical Strain Platform, EQUicycler, for 3D Tissue Engineered Constructs. Biomed Res Int 2017, 2017:3609703.

39. Elsaadany M, Yan KC, Yildirim-Ayan E: Predicting cell viability within tissue scaffolds under equiaxial strain: multi-scale finite element model of collagen-cardiomyocytes constructs. Biomech Model Mechanobiol 2017, 16:1049–1063.

40. Ren Q, Khoo WH, Corr AP, Phan TG, Croucher PI, Stewart SA: Gene expression predicts dormant metastatic breast cancer cell phenotype. Breast Cancer Res 2022, 24:10.

41. Troyano A, Sancho P, Fernandez C, de Blas E, Bernardi P, Aller P: The selection between apoptosis and necrosis is differentially regulated in hydrogen peroxide-treated and glutathione-depleted human promonocytic cells. Cell Death Differ 2003, 10:889–898.

42. Venegas V, Halberg MC: Measurement of mitochondrial DNA copy number. Methods Mol Biol 2012, 837:327–335.

43. Conway T, Wazny J, Bromage A, Tymms M, Sooraj D, Williams ED, Beresford-Smith B: Xenome--a tool for classifying reads from xenograft samples. Bioinformatics 2012, 28:i172–178.

44. Kluin RJC, Kemper K, Kuilman T, de Ruiter JR, Iyer V, Forment JV, Cornelissen-Steijger P, de Rink I, Ter Brugge P, Song JY, et al: XenofilteR: computational deconvolution of mouse and human reads in tumor xenograft sequence data. BMC Bioinformatics 2018, 19:366.

45. Bray NL, Pimentel H, Melsted P, Pachter L: Near-optimal probabilistic RNA-seq quantification. Nat Biotechnol 2016, 34:525–527.

46. Kim D, Paggi JM, Park C, Bennett C, Salzberg SL: Graph-based genome alignment and genotyping with HISAT2 and HISAT-genotype. Nat Biotechnol 2019, 37:907–915.

47. Liao Y, Smyth GK, Shi W: featureCounts: an efficient general purpose program for assigning sequence reads to genomic features. Bioinformatics 2014, 30:923–930.

48. Shen W, Le S, Li Y, Hu F: SeqKit: A Cross-Platform and Ultrafast Toolkit for FASTA/Q File Manipulation. PLoS One 2016, 11:e0163962.

49. Frankish A, Diekhans M, Jungreis I, Lagarde J, Loveland JE, Mudge JM, Sisu C, Wright JC, Armstrong J, Barnes I, et al: Gencode 2021. Nucleic Acids Res 2021, 49:D916–D923.

50. Xie Z, Bailey A, Kuleshov MV, Clarke DJB, Evangelista JE, Jenkins SL, Lachmann A, Wojciechowicz ML, Kropiwnicki E, Jagodnik KM, et al: Gene Set Knowledge Discovery with Enrichr. Curr Protoc 2021, 1:e90.

51. Ge SX, Jung D, Yao R: ShinyGO: a graphical gene-set enrichment tool for animals and plants. Bioinformatics 2020, 36:2628–2629.

52. Su S, Cao J, Meng X, Liu R, Vander Ark A, Woodford E, Zhang R, Stiver I, Zhang X, Madaj ZB, et al: Enzalutamide-induced and PTH1R-mediated TGFBR2 degradation in osteoblasts confers resistance in prostate cancer bone metastases. Cancer Lett 2022, 525:170–178.

53. Rashid OM, Nagahashi M, Ramachandran S, Graham L, Yamada A, Spiegel S, Bear HD, Takabe K: Resection of the primary tumor improves survival in metastatic breast cancer by reducing overall tumor burden. Surgery 2013, 153:771–778.

54. Vashist YK, Effenberger KE, Vettorazzi E, Riethdorf S, Yekebas EF, Izbicki JR, Pantel K: Disseminated tumor cells in bone marrow and the natural course of resected esophageal cancer. Ann Surg 2012, 255:1105–1112.

55. Gonzalez-Gualda E, Baker AG, Fruk L, Munoz-Espin D: A guide to assessing cellular senescence in vitro and in vivo. FEBS J 2021, 288:56–80.

56. Hernandez-Segura A, Nehme J, Demaria M: Hallmarks of Cellular Senescence. Trends Cell Biol 2018, 28:436–453.

57. Paschalis EP, Recker R, DiCarlo E, Doty SB, Atti E, Boskey AL: Distribution of collagen cross-links in normal human trabecular bone. J Bone Miner Res 2003, 18:1942–1946.

58. Tzaphlidou M: Bone architecture: collagen structure and calcium/phosphorus maps. J Biol Phys 2008, 34:39–49.

59. Miyamoto DT, Zheng Y, Wittner BS, Lee RJ, Zhu H, Broderick KT, Desai R, Fox DB, Brannigan BW, Trautwein J, et al: RNA-Seq of single prostate CTCs implicates noncanonical Wnt signaling in antiandrogen resistance. Science 2015, 349:1351–1356.

60. Molina JR, Sun Y, Protopopova M, Gera S, Bandi M, Bristow C, McAfoos T, Morlacchi P, Ackroyd J, Agip AA, et al: An inhibitor of oxidative phosphorylation exploits cancer vulnerability. Nat Med 2018, 24:1036–1046.

61. Li N, Ragheb K, Lawler G, Sturgis J, Rajwa B, Melendez JA, Robinson JP: Mitochondrial complex I inhibitor rotenone induces apoptosis through enhancing mitochondrial reactive oxygen species production. J Biol Chem 2003, 278:8516–8525.

62. Munoz F, Martin ME, Salinas M, Fando JL: Carbonyl cyanide p-trifluoromethoxyphenylhydrazone (FCCP) induces initiation factor 2 alpha phosphorylation and translation inhibition in PC12 cells. FEBS Lett 2001, 492:156–159.

63. Huang S, Liu Y, Chen Z, Wang M, Jiang VC: PIK-75 overcomes venetoclax resistance via blocking PI3K-AKT signaling and MCL-1 expression in mantle cell lymphoma. Am J Cancer Res 2022, 12:1102–1115.

64. O’Brien S, Golubovskaya VM, Conroy J, Liu S, Wang D, Liu B, Cance WG: FAK inhibition with small molecule inhibitor Y15 decreases viability, clonogenicity, and cell attachment in thyroid cancer cell lines and synergizes with targeted therapeutics. Oncotarget 2014, 5:7945–7959.

65. Lin HM, Lee BY, Castillo L, Spielman C, Grogan J, Yeung NK, Kench JG, Stricker PD, Haynes AM, Centenera MM, et al: Effect of FAK inhibitor VS-6063 (defactinib) on docetaxel efficacy in prostate cancer. Prostate 2018, 78:308–317.

66. Fares J, Fares MY, Khachfe HH, Salhab HA, Fares Y: Molecular principles of metastasis: a hallmark of cancer revisited. Signal Transduct Target Ther 2020, 5:28.

67. Al-Moraissi EA, Dahan AA, Alwadeai MS, Oginni FO, Al-Jamali JM, Alkhutari AS, Al-Tairi NH, Almaweri AA, Al-Sanabani JS: What surgical treatment has the lowest recurrence rate following the management of keratocystic odontogenic tumor?: A large systematic review and meta-analysis. J Craniomaxillofac Surg 2017, 45:131–144.

68. Wang L, Lu B, He M, Wang Y, Wang Z, Du L: Prostate Cancer Incidence and Mortality: Global Status and Temporal Trends in 89 Countries From 2000 to 2019. Front Public Health 2022, 10:811044.

69. Boire A, Coffelt SB, Quezada SA, Vander Heiden MG, Weeraratna AT: Tumour Dormancy and Reawakening: Opportunities and Challenges. Trends Cancer 2019, 5:762–765.

70. Kollermann J, Heseding B, Helpap B, Kollermann MW, Pantel K: Comparative immunocytochemical assessment of isolated carcinoma cells in lymph nodes and bone marrow of patients with clinically localized prostate cancer. Int J Cancer 1999, 84:145–149.

71. Risson E, Nobre AR, Maguer-Satta V, Aguirre-Ghiso JA: The current paradigm and challenges ahead for the dormancy of disseminated tumor cells. Nat Cancer 2020, 1:672–680.

72. Linxweiler J, Hajili T, Zeuschner P, Menger MD, Stockle M, Junker K, Saar M: Primary Tumor Resection Decelerates Disease Progression in an Orthotopic Mouse Model of Metastatic Prostate Cancer. Cancers (Basel) 2022, 14.

73. Sow Y, Sow O, Fall B, Sine B, Sarr A, Ze Ondo C, Diao B, Ndoye AK, Ba M: Impact of tumor cytoreduction in metastatic prostate cancer. Res Rep Urol 2019, 11:137–142.

74. Stensland KD, Caram MV, Burns JA, Sparks JB, Shin C, Zaslavsky A, Hollenbeck BK, Tsodikov A, Skolarus TA: Recurrence, metastasis, and survival after radical prostatectomy in the era of advanced treatments. Journal of Clinical Oncology 2022, 40:1.

75. Weckermann D, Muller P, Wawroschek F, Harzmann R, Riethmuller G, Schlimok G: Disseminated cytokeratin positive tumor cells in the bone marrow of patients with prostate cancer: detection and prognostic value. J Urol 2001, 166:699–703.

76. Weckermann D, Polzer B, Ragg T, Blana A, Schlimok G, Arnholdt H, Bertz S, Harzmann R, Klein CA: Perioperative activation of disseminated tumor cells in bone marrow of patients with prostate cancer. J Clin Oncol 2009, 27:1549–1556.

77. Khoo WH, Ledergor G, Weiner A, Roden DL, Terry RL, McDonald MM, Chai RC, De Veirman K, Owen KL, Opperman KS, et al: A niche-dependent myeloid transcriptome signature defines dormant myeloma cells. Blood 2019, 134:30–43.

78. Chery L, Lam HM, Coleman I, Lakely B, Coleman R, Larson S, Aguirre-Ghiso JA, Xia J, Gulati R, Nelson PS, et al: Characterization of single disseminated prostate cancer cells reveals tumor cell heterogeneity and identifies dormancy associated pathways. Oncotarget 2014, 5:9939–9951.

79. Quayle LA, Spicer A, Ottewell PD, Holen I: Transcriptomic Profiling Reveals Novel Candidate Genes and Signalling Programs in Breast Cancer Quiescence and Dormancy. Cancers (Basel) 2021, 13.

80. Yu-Lee LY, Yu G, Lee YC, Lin SC, Pan J, Pan T, Yu KJ, Liu B, Creighton CJ, Rodriguez-Canales J, et al: Osteoblast-Secreted Factors Mediate Dormancy of Metastatic Prostate Cancer in the Bone via Activation of the TGFbetaRIII-p38MAPK-pS249/T252RB Pathway. Cancer Res 2018, 78:2911–2924.

81. Borgen E, Rypdal MC, Sosa MS, Renolen A, Schlichting E, Lonning PE, Synnestvedt M, Aguirre-Ghiso JA, Naume B: NR2F1 stratifies dormant disseminated tumor cells in breast cancer patients. Breast Cancer Res 2018, 20:120.

82. Gao XL, Zheng M, Wang HF, Dai LL, Yu XH, Yang X, Pang X, Li L, Zhang M, Wang SS, et al: NR2F1 contributes to cancer cell dormancy, invasion and metastasis of salivary adenoid cystic carcinoma by activating CXCL12/CXCR4 pathway. BMC Cancer 2019, 19:743.

83. Khalil BD, Sanchez R, Rahman T, Rodriguez-Tirado C, Moritsch S, Martinez AR, Miles B, Farias E, Mezei M, Nobre AR, et al: An NR2F1-specific agonist suppresses metastasis by inducing cancer cell dormancy. J Exp Med 2022, 219.

84. Jakob M, Sharaf K, Schirmer M, Leu M, Kuffer S, Bertlich M, Ihler F, Haubner F, Canis M, Kitz J: Role of cancer stem cell markers ALDH1, BCL11B, BMI-1, and CD44 in the prognosis of advanced HNSCC. Strahlenther Onkol 2021, 197:231–245.

85. Federer-Gsponer JR, Muller DC, Zellweger T, Eggimann M, Marston K, Ruiz C, Seifert HH, Rentsch CA, Bubendorf L, Le Magnen C: Patterns of stemness-associated markers in the development of castration-resistant prostate cancer. Prostate 2020, 80:1108–1117.

86. Cai C, Wang H, He HH, Chen S, He L, Ma F, Mucci L, Wang Q, Fiore C, Sowalsky AG, et al: ERG induces androgen receptor-mediated regulation of SOX9 in prostate cancer. J Clin Invest 2013, 123:1109–1122.

87. Thomsen MK, Ambroisine L, Wynn S, Cheah KS, Foster CS, Fisher G, Berney DM, Moller H, Reuter VE, Scardino P, et al: SOX9 elevation in the prostate promotes proliferation and cooperates with PTEN loss to drive tumor formation. Cancer Res 2010, 70:979–987.

88. Whitburn J, Rao SR, Morris EV, Tabata S, Hirayama A, Soga T, Edwards JR, Kaya Z, Palmer C, Hamdy FC, Edwards CM: Metabolic profiling of prostate cancer in skeletal microenvironments identifies G6PD as a key mediator of growth and survival. Sci Adv 2022, 8:eabf9096.

89. Shiozawa Y, Pedersen EA, Havens AM, Jung Y, Mishra A, Joseph J, Kim JK, Patel LR, Ying C, Ziegler AM, et al: Human prostate cancer metastases target the hematopoietic stem cell niche to establish footholds in mouse bone marrow. J Clin Invest 2011, 121:1298–1312.

90. Aguirre Ghiso JA: Inhibition of FAK signaling activated by urokinase receptor induces dormancy in human carcinoma cells in vivo. Oncogene 2002, 21:2513–2524.

91. Ivey NS, Renner NA, Moroney-Rasmussen T, Mohan M, Redmann RK, Didier PJ, Alvarez X, Lackner AA, MacLean AG: Association of FAK activation with lentivirus-induced disruption of blood-brain barrier tight junction-associated ZO-1 protein organization. J Neurovirol 2009, 15:312–323.

92. Greenbaum A, Chan KY, Dobreva T, Brown D, Balani DH, Boyce R, Kronenberg HM, McBride HJ, Gradinaru V: Bone CLARITY: Clearing, imaging, and computational analysis of osteoprogenitors within intact bone marrow. Sci Transl Med 2017, 9.

93. Manna F, Karkampouna S, Zoni E, De Menna M, Hensel J, Thalmann GN, Kruithof-de Julio M: Metastases in Prostate Cancer. Cold Spring Harb Perspect Med 2019, 9.

