## Supplementary Figures and raw pictures for western blotting and agarose gel for "Tumor removal limits prostate cancer cell dissemination in bone and osteoblasts induce cancer cell dormancy through focal adhesion kinase"

[1]Department of Cell and Cancer Biology, College of Medicine and Life Sciences, the University of Toledo, Toledo, OH 43614; [2]Department of Pediatrics and Human Development, College of Human Medicine, Michigan State University, Grand Rapids, MI 49503; [3]Bioengineering Department, the University of Toledo, Toledo, OH 43606; [4]Department of Pathology, College of Medicine and Life Sciences, the University of Toledo, Toledo, OH 43614.

Contents:

- **Supplementary Tables S1-S6 (provided as a separate xlsx excel file)**
- **Supplementary Movies 1-6 (provided as separate files)**
- **Supplementary Figures S1-S11**
- **Supplementary Data (original western blot and DNA gel images)**

**Supplementary Tables**

**Table S1. Sources of reagents, chemicals and plasmids.**

**Table S2. Primers used in the study.**

**Table S3. Estimated DTC numbers in various mouse tissues post tumor xenografts with or without tumor removal.**

**Table S4. Antibodies used in the study.**

**Table S5. DE genes in co-cultured dormant C4-2B cells by three different pipelines.**

**Table S6. Functional annotation of DE genes indicates enrichment of cell adhesion pathway in dormant cells.**

**Supplementary Movies**

**Movie S1 C4-2B/GFP-FAK cells treated with DMSO.**

**Movie S2 C4-2B/GFP-FAK cells treated with 1 μM PF-562271 (PF-271).**

**Movie S3 C4-2B/GFP-FAK cells cultured alone (in regular media for MC3T3-E1).**

**Movie S4 C4-2B/GFP-FAK cells co-cultured with MC3T3-E1 cells.**

**Movie S5 C4-2B/GFP-FAK cells treated with DMSO (in regular media for OP-9).**

**Movie S6 C4-2B/GFP-FAK cells co-cultured with OP-9 cells.**

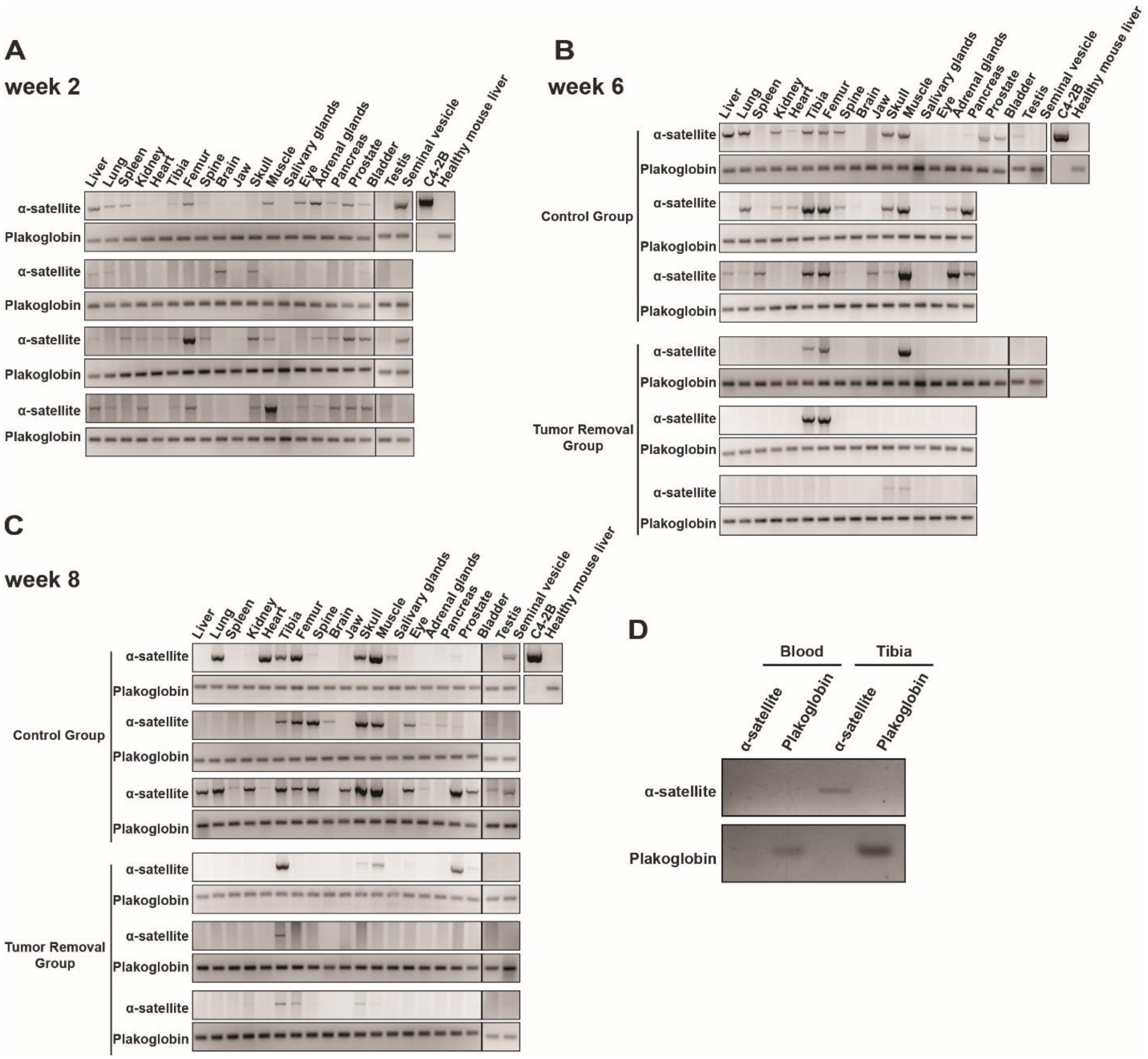


**Figure S1. Species-specific PCR for DTC detection from various tissues of each individual mouse**

The original DNA gels for the species-specific PCR. 2 µg total genomic DNA for α-satellite PCR and 0.5 µg total genomic DNA for plakoglobin PCR were loaded. Post-tumor cell injections, wk 2 (**A**), wk 6 (**B**), and wk 8 (**C**). **D**. Representative picture of the PCRs on the genomic DNA from the peripheral blood and the tibia of the same tumor-bearing mouse. All PCR experiments were performed at least twice.

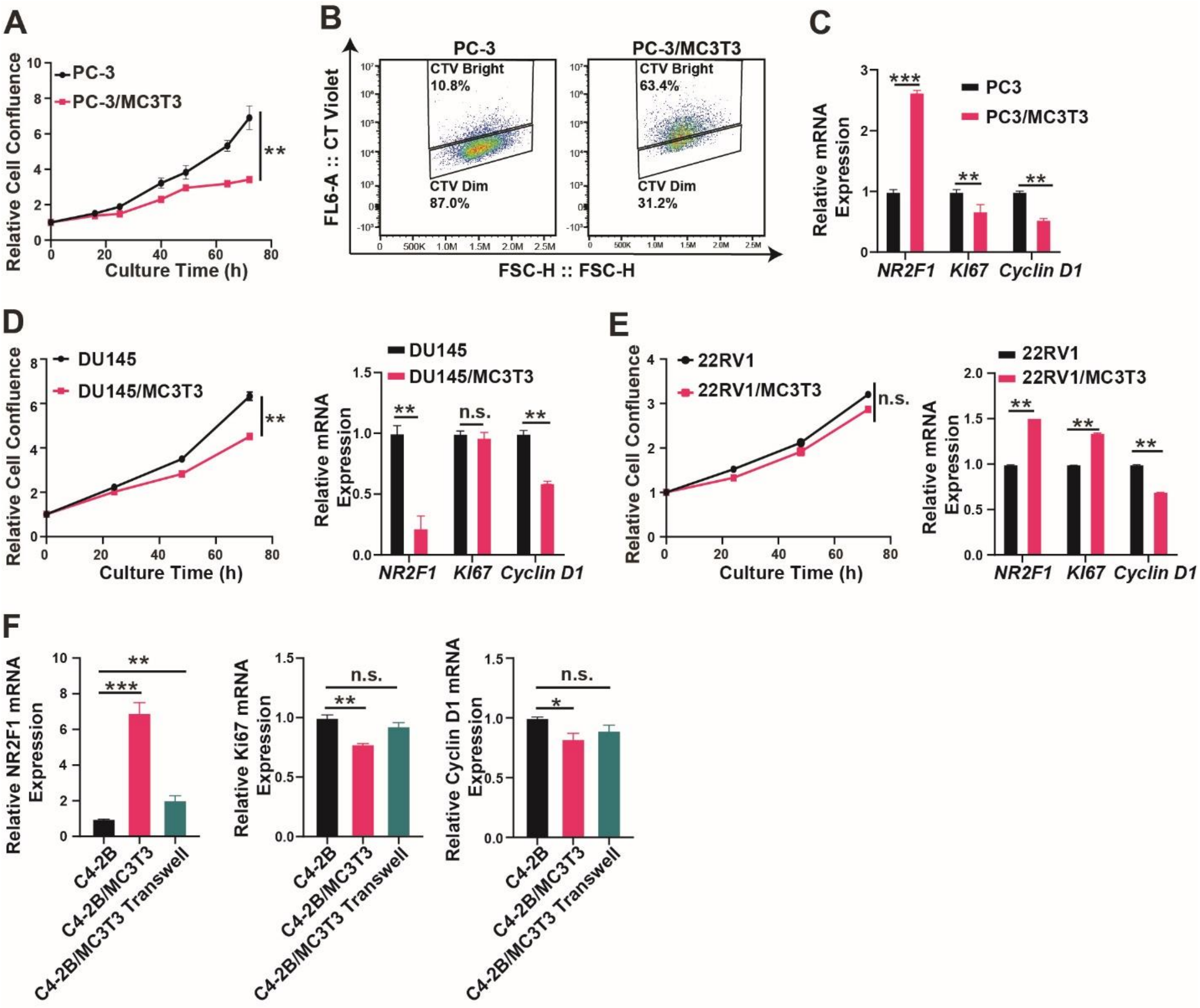


**Figure S2. MC3T3-E1 inhibited the cell proliferation of PC-3 and DU145 cells**

**A-C.** PC-3/GFP cells were cultured alone or co-cultured with MC3T3-E1 cells for 72 h. Cell proliferation by Incucyte imaging (**A**), cell division by CT Violet dye(**B**), and the marker gene mRNA expression assay by qRT-PCR (**C**) were performed in the same manner as described in **Figure 2**. **D/E**. DU145/GFP cells (**D**) and 22Rv1/GFP cells (**E**) were cultured alone or co-cultured with MC3T3-E1 cells for 72 h. Cell proliferation curves (Incucyte, left panel) and mRNA expression changes for NR2F1/Ki67/Cyclin D1 (qRT-PCR, right panel) were shown. **F**. The mRNA expression changes of NR2F1/Ki67/Cyclin D1 in C4-2B cells co-cultured directly or indirectly (in transwell) with MC3T3-E1 for 72 h, compared to C4-2B cells cultured alone, were examined by qRT-PCR. All the experiments were independently repeated at least twice. The data in curves and bar plots are presented as the mean ± SD of each set of triplicate samples. * $p < 0.05$, ** $p < 0.01$, *** $p < 0.001$, and n.s. non-significant.

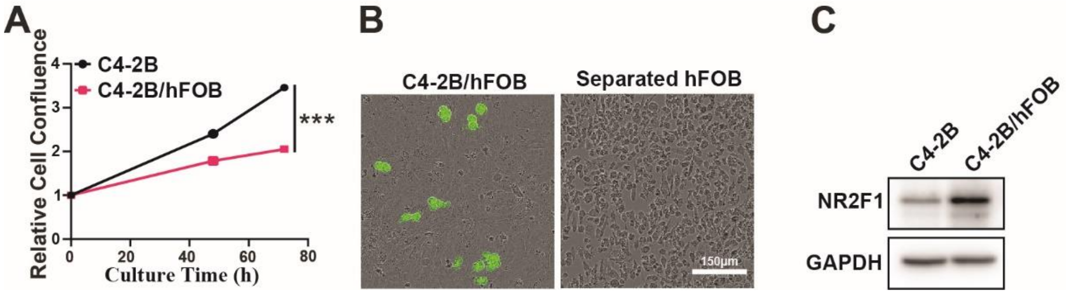


**Figure S3. Human osteoblast hFOB1.19 induced C4-2B cell dormancy in the mixed co-culture**

**A**. C4-2B/GFP cells were cultured alone or co-cultured with hFOB1.19 cells for 72 h under 34℃. Cell proliferation was plotted as the relative confluence to the start point and the data are presented as the mean ± SD of each set of triplicate samples. **B**. Separation of C4-2B cells (GFP labeled) from hFOB cells (no label). **C**. The NR2F1 protein levels in C4-2B cells cultured alone and separated from hFOB co-culture were examined by immunoblotting. All the experiments were independently repeated at least twice. *** $p < 0.001$.

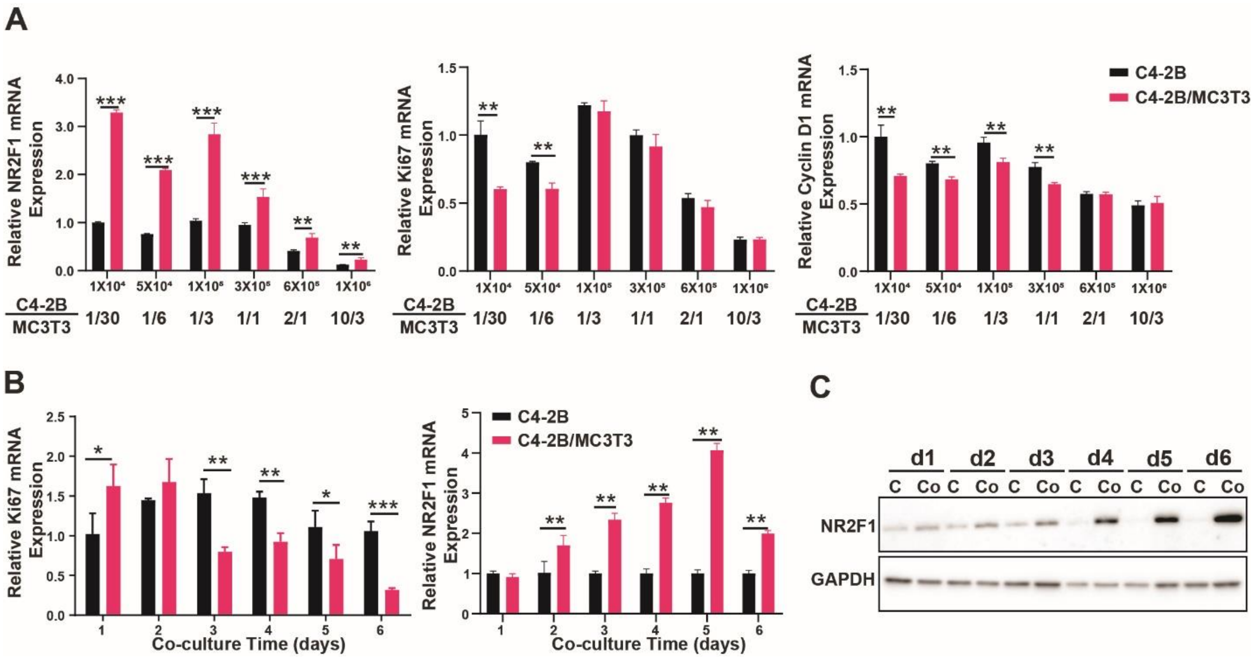


**Figure S4. Evaluation of cell number ratio and culture time for the mixed co-culture**

C4-2B cells were cultured alone or co-cultured with MC3T3-E1 cells at the indicated ratios (**A**) or times (**B**). The mRNA expression of NR2F1/Ki67/Cyclin D1 were examined by qRT-PCR. The data are presented as the mean ± SD of each set of triplicate samples. **C**. NR2F1 proteins levels in C4-2B cells co-cultured (Co) or cultured alone (C) for indicated days were examined by immunoblotting. All the experiments were independently repeated twice. * $p < 0.05$, ** $p < 0.01$, *** $p < 0.001$.

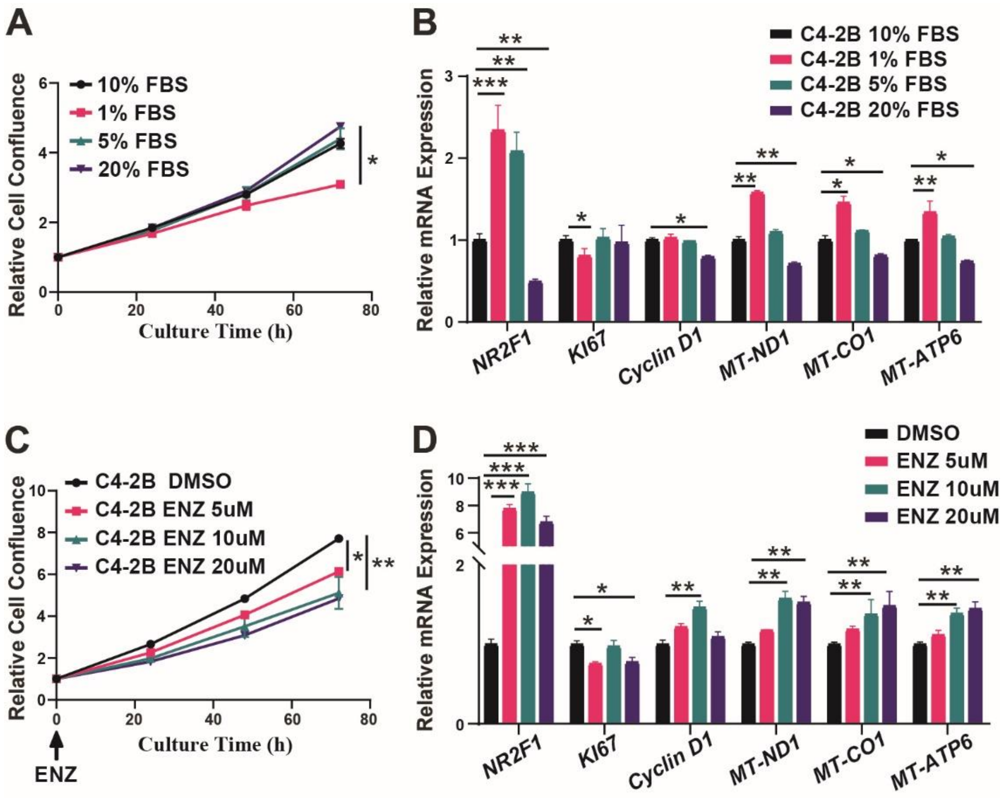


**Figure S5. Serum starvation or enzalutamide treatments inhibited C4-2B cell proliferation but did not induce dormancy**

**A/B**. C4-2B cells were treated with media containing different concentrations of FBS as indicated for 72 h. Proliferation (**A**) and marker gene expression (**B**) were examined. **C/D**. C4-2B cells were treated with DMSO (as vehicle) and enzalutamide at 5/10/20 μM as indicated for 72 h. Proliferation (**C**) and marker gene expression (**D**) were examined. All the experiments were independently repeated at least twice. The data in curves and bar plots are presented as the mean ± SD of each set of triplicate samples. * $p < 0.05$, ** $p < 0.01$, *** $p < 0.001$.

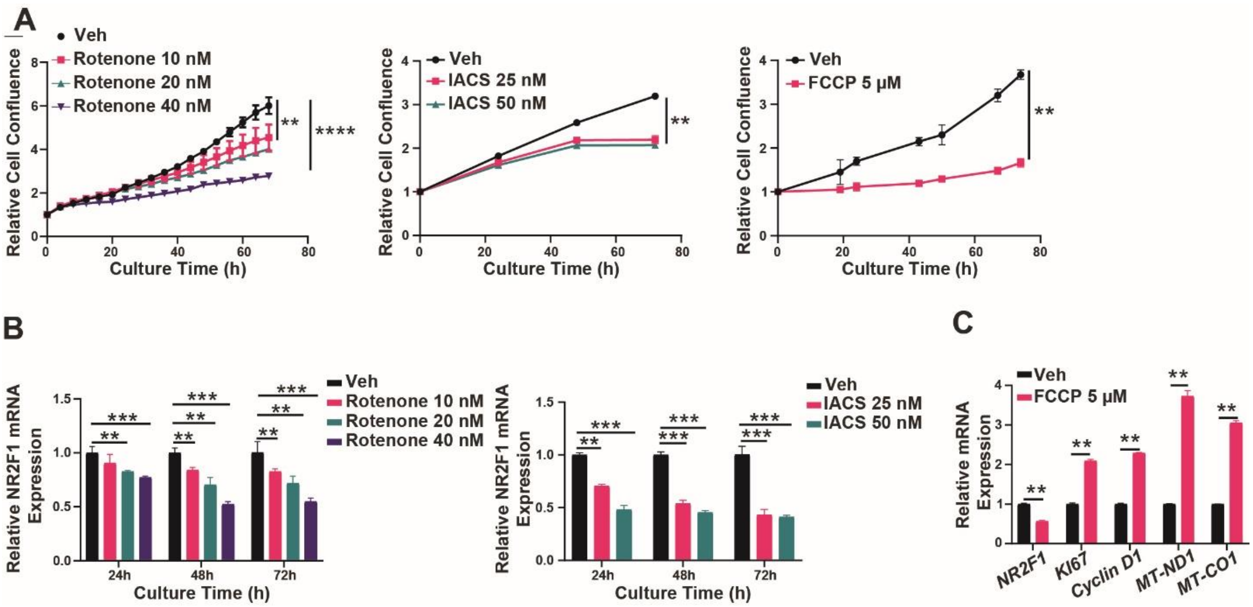


**Figure S6. Mitochondrial inhibitors inhibited C4-2B cell proliferation but did not induce dormancy**

C4-2B cells were treated with rotenone, IACS-010759 (IACS), and FCCP at the indicated concentrations for 24/48/72 h. on cell proliferation. Proliferation was monitored by Incucyte and plotted in **A**. The mRNA expressions of NR2F1 as well as other marker genes by qRT PCR were shown in **B** and **C.** All the experiments were independently repeated at least twice. The data in curves and bar plots are presented as the mean ± SD of each set of triplicate samples. * $p < 0.05$, ** $p < 0.01$, *** $p < 0.001$, **** $p < 0.0001$.

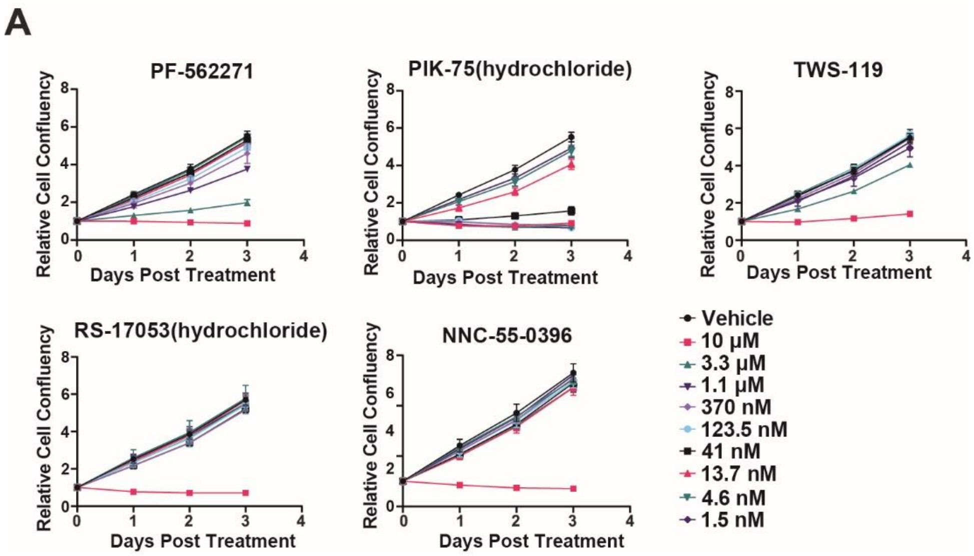


**Figure S7. Growth curves of C4-2B cells in response to the top drugs from the virtual screen**

C4-2B cells were treated with the top drugs from the virtual prediction (**Figure 5A**) at the indicated concentrations for 72 h. The proliferation was measured by relative cell confluence from the Incucyte imaging. All the experiments were independently repeated twice. The data are presented as the mean ± SD of each set of triplicate samples.

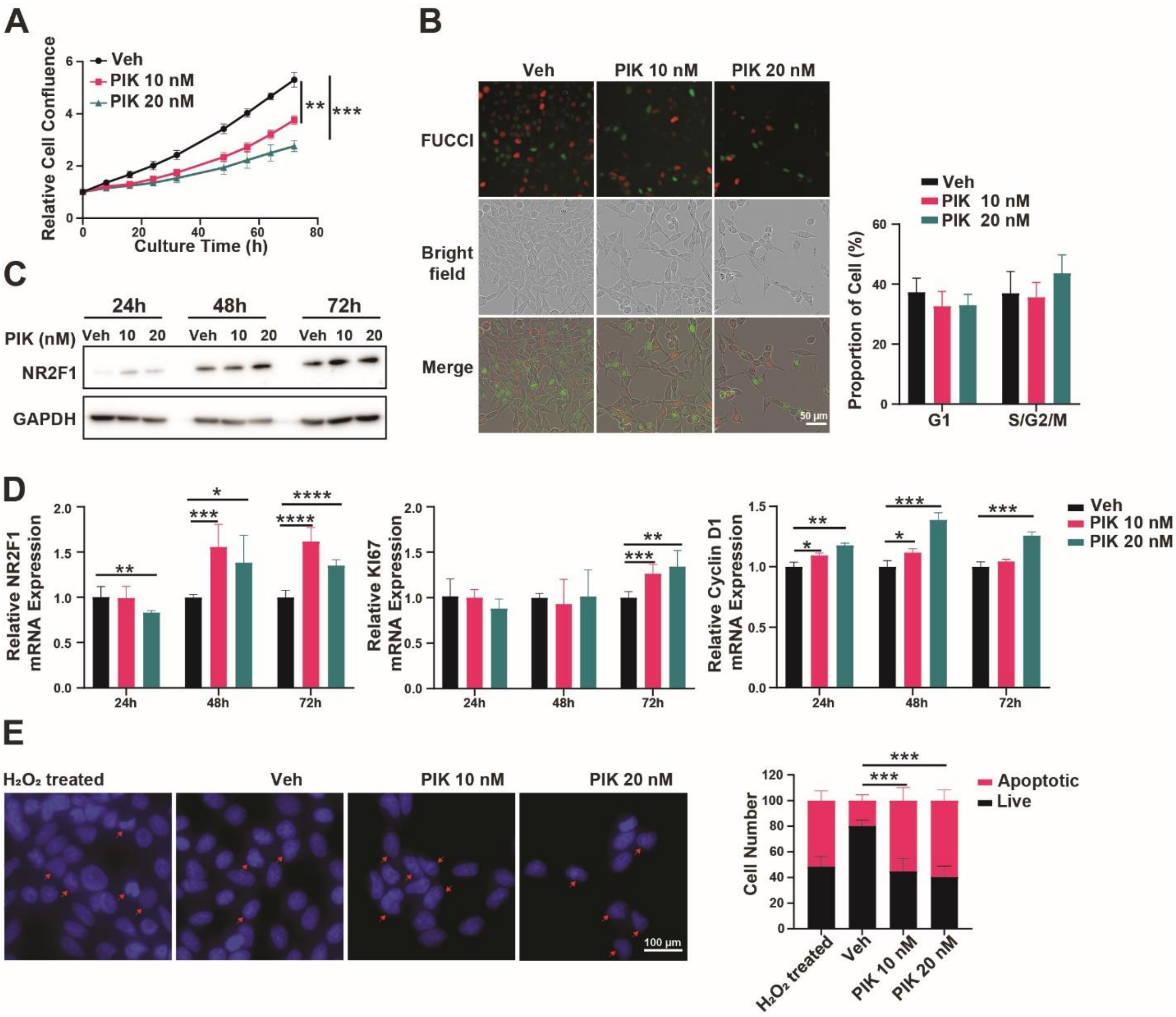


**Figure S8. PIK-75 inhibited C4-2B cells proliferation but could not mimic dormancy**

C4-2B cells were treated with PIK-75 (PIK in short) at 10/20 nM or Vehicle (Veh, DMSO) for 72 h as indicated. Cell proliferation monitoring by Incucyte (**A**), cell cycle phase distribution by FUCCI (**B**), Time-lapse NR2F1 protein expressions by immunoblotting (**C**), and mRNA expressions of NR2F1/Ki67/Cyclin D1 (**D**) by qRT-PCR, were performed in the same manner as described in **Figure 2**. **E**. Analyses of apoptotic cells (red arrow) in C4-2B cells treated with 10/20 nM of PIK or Vehicle (Veh, DMSO) for 72 h. $H_2O_2$ treatment for positive control. Scale bars, 50 μm (**B**) and 100 μm (**E**). All the experiments were independently repeated twice. The data in curves and bar plots are presented as the mean ± SD of each set of triplicate samples.
* $p < 0.05$, *** $p < 0.01$, *** $p < 0.001$.

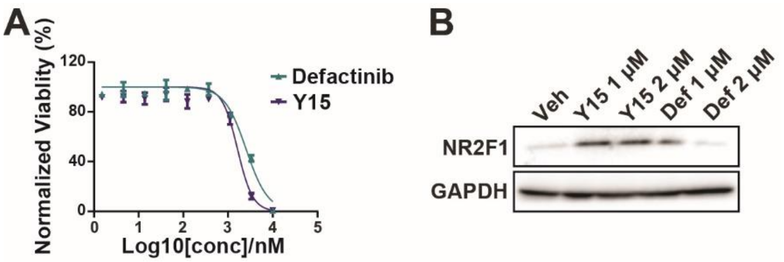


**Figure S9. Defactinib and Y15, two other FAK inhibitors, induced dormancy-mimicking effects in C4-2B cells.**

**A**. C4-2B cells were treated with FAK inhibitors defactinib or Y15 at various concentrations for 72 h as indicated. Cell viability was measured via CCK-8 assay in 96-well plates. Relative viability was calculated by normalizing to the DMSO-treated group. The data are presented as the mean ± SD of each set of triplicate samples. **B**. FAK inhibitors increased the NR2F1 expression at the protein level in C4-2B cells. Both experiments were repeated twice.

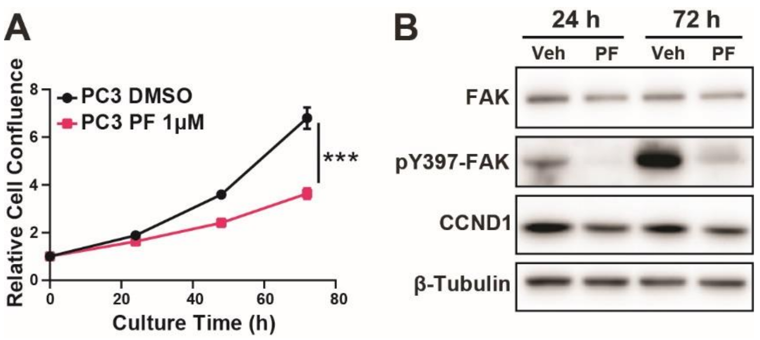


**Figure S10. PF-271 induced dormancy-mimicking effects in PC-3 cells.**

PC-3 cells were treated PF-271 (1 μM) or Veh (Vehicle, DMSO) for 24/72 h as indicated. **A.** Proliferation was monitored by Incucyte imaging and plotted as relative confluence (mean ± SD of each set of triplicate samples). **B.** The protein levels of total FAK, pY397-FAK, and CCND1 were examined by the immunoblotting.  Both experiments were repeated twice.

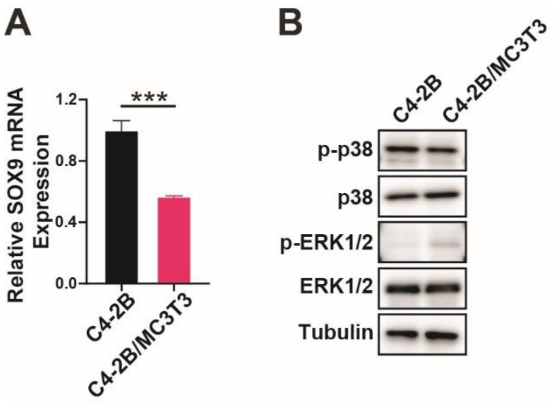


**Figure S11. Unique features of the osteoblast-induced C4-2B cell dormancy**

C4-2B cells were cultured alone or co-cultured with MC3T3-E1 cells for 72 h. **A**. SOX9 mRNA expressions were examined by qRT-PCR with human-specific primers. Data are presented as the mean ± SD of each set of triplicate samples. *** $p < 0.001$. **B**. The protein levels of total p38, phospho-p38 (p-p38), total ERK1/2, and phospho-ERK1/2 were examined by immunoblotting. The experiments were repeated twice.

### Supplementary Data

Original Images for DNA gels, Western blots and primer specificity verification gels

Figure 1

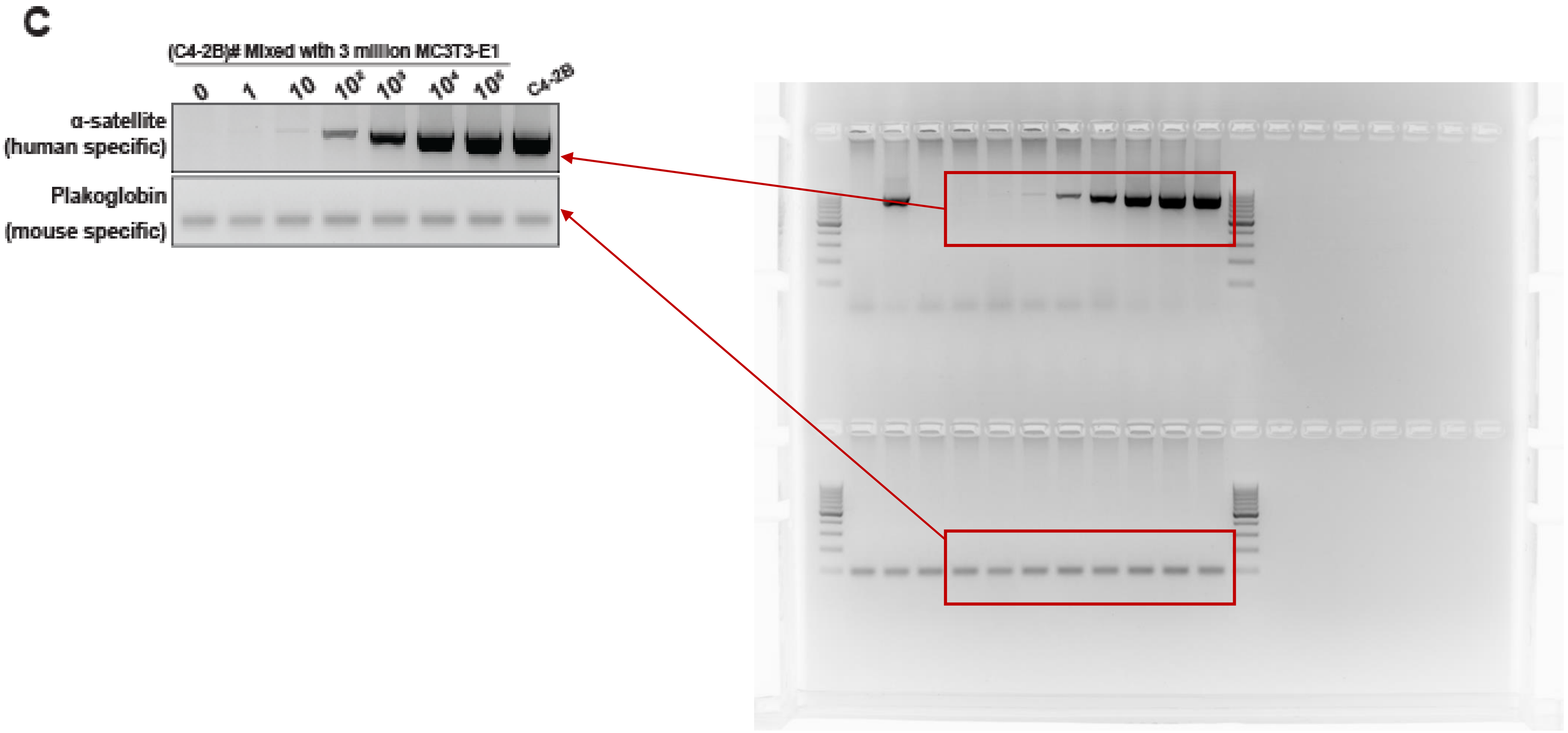
C
(C4-2B)# Mixed with 3 million MC3T3-E1
0
1
10
$10^2$
$10^3$
$10^4$
$10^5$
C4-2B
α-satellite (human specific)
Plakoglobin (mouse specific)

Figure 1

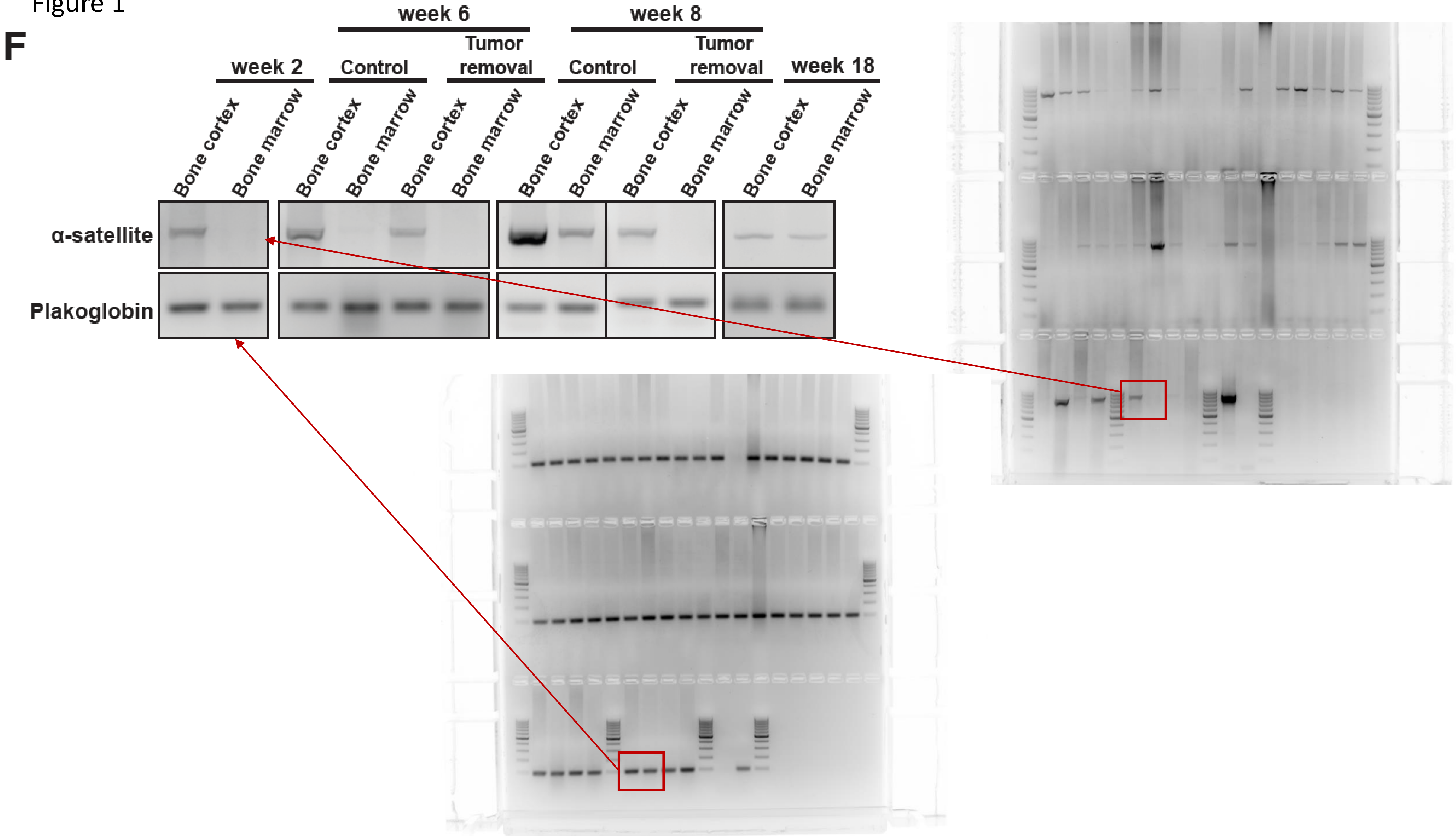
F
week 2
week 6
week 8
Control
Tumor removal
Control
Tumor removal
week 18
Bone cortex
Bone marrow
Bone cortex
Bone marrow
Bone cortex
Bone marrow
Bone cortex
Bone marrow
Bone cortex
Bone marrow
Bone cortex
Bone marrow
α-satellite
Plakoglobin

Figure 1

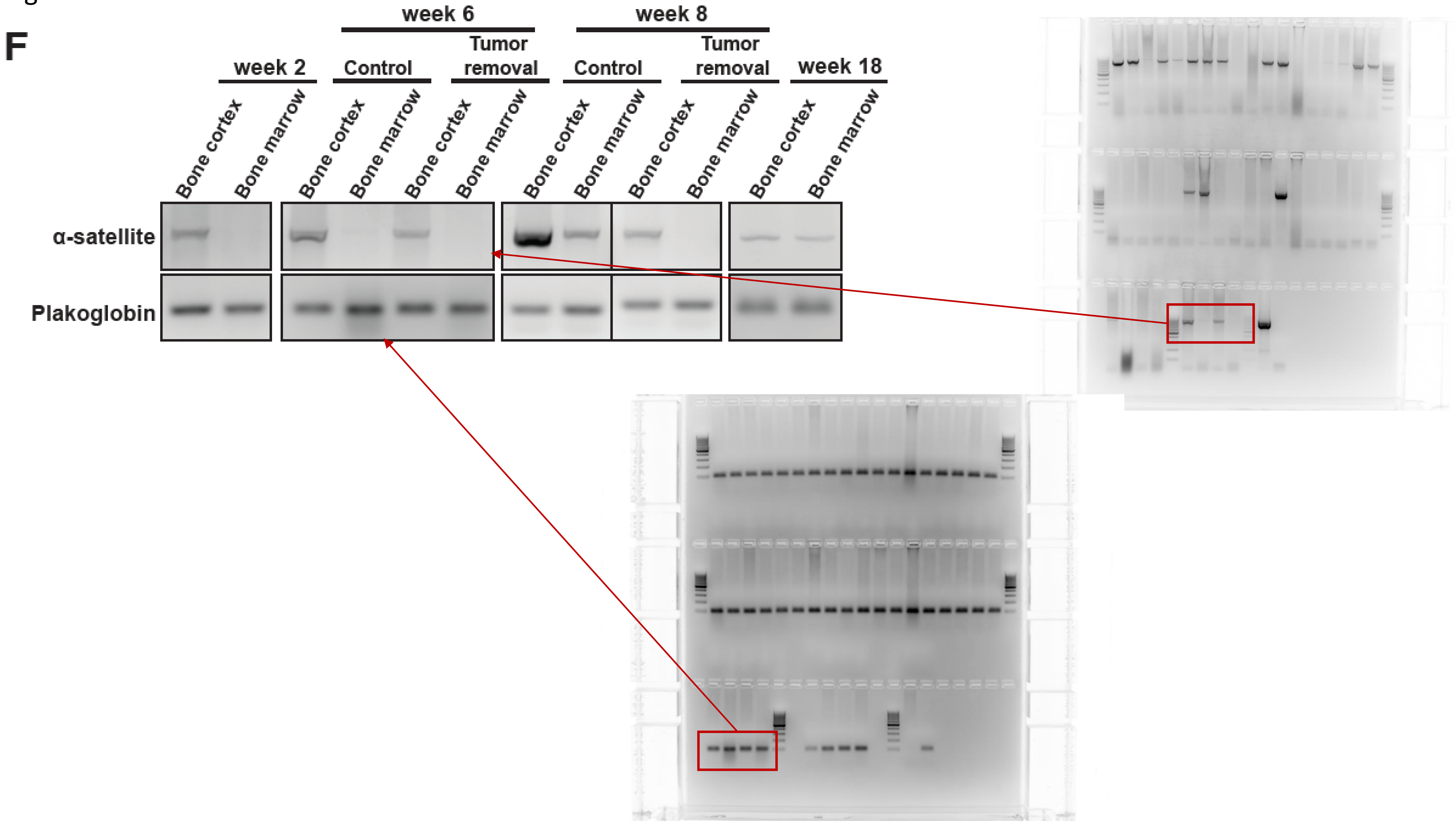
F
week 6
week 8
Tumor removal
Tumor removal
week 2
Control
Control
week 18
Bone cortex
Bone marrow
Bone cortex
Bone marrow
Bone cortex
Bone marrow
Bone cortex
Bone marrow
Bone cortex
Bone marrow
Bone cortex
Bone marrow
α-satellite
Plakoglobin

Figure 1

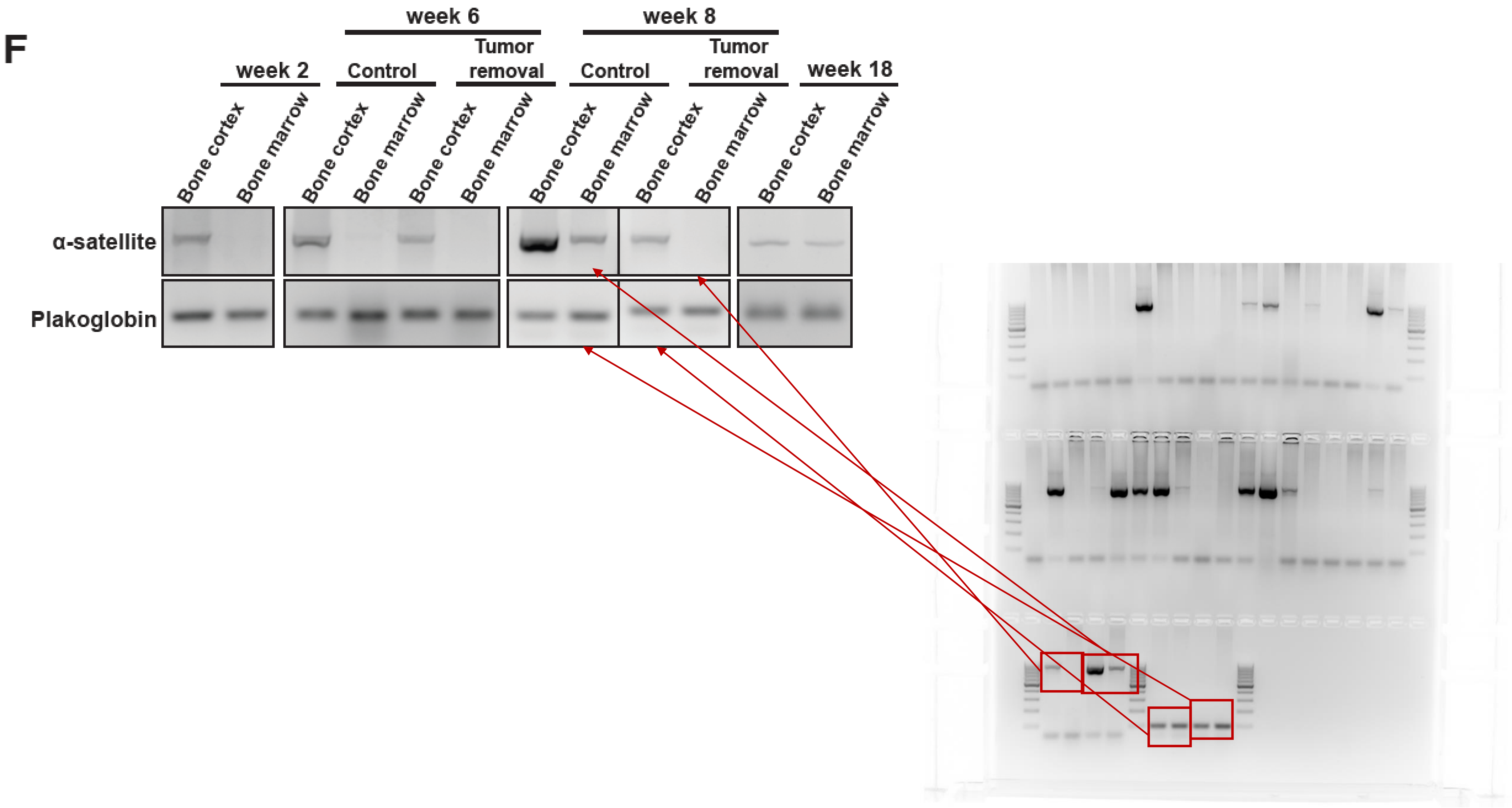
F
week 6
week 8
Tumor removal
Tumor removal
week 2
Control
Control
week 18
Bone cortex
Bone marrow
Bone cortex
Bone marrow
Bone cortex
Bone marrow
Bone cortex
Bone marrow
Bone cortex
Bone marrow
Bone cortex
Bone marrow
α-satellite
Plakoglobin

Figure 1

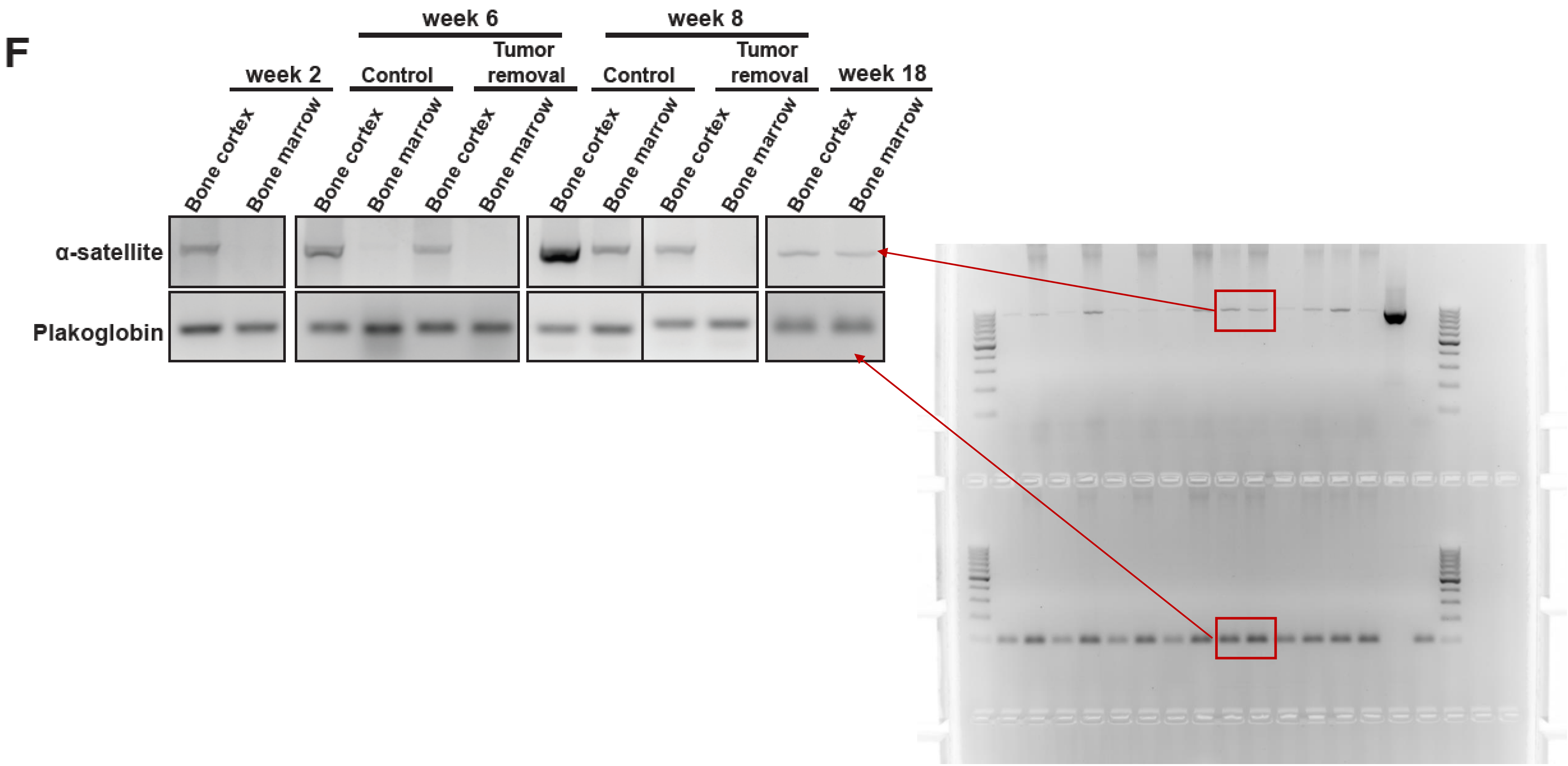
F
week 2
week 6
Control
Tumor removal
week 8
Control
Tumor removal
week 18
Bone cortex
Bone marrow
Bone cortex
Bone marrow
Bone cortex
Bone marrow
Bone cortex
Bone marrow
Bone cortex
Bone marrow
Bone cortex
Bone marrow
α-satellite
Plakoglobin

Figure 2

I

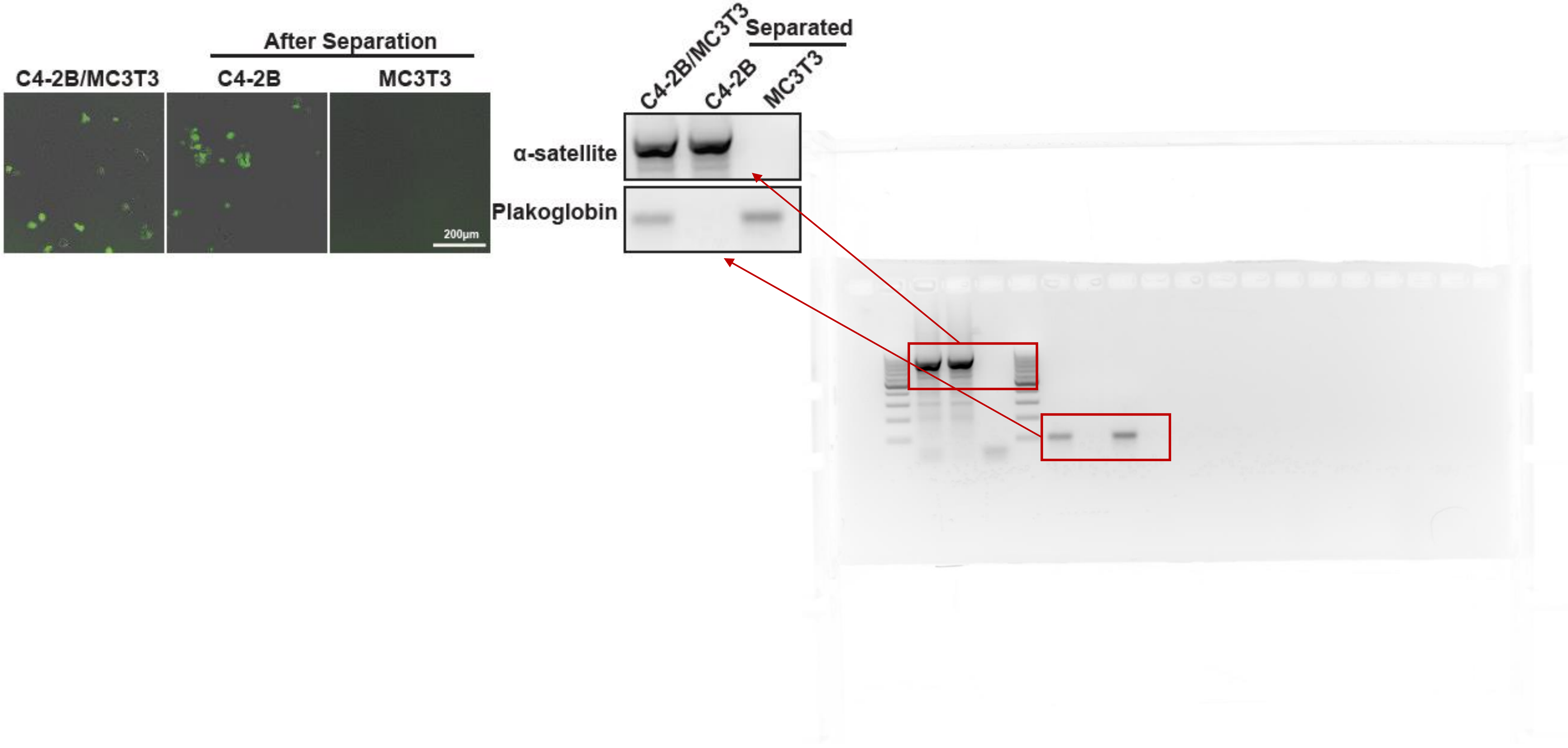
After Separation
C4-2B/MC3T3
C4-2B
MC3T3
200μm
C4-2B/MC3T3
C4-2B
MC3T3
Separated
α-satellite
Plakoglobin

Figure 2

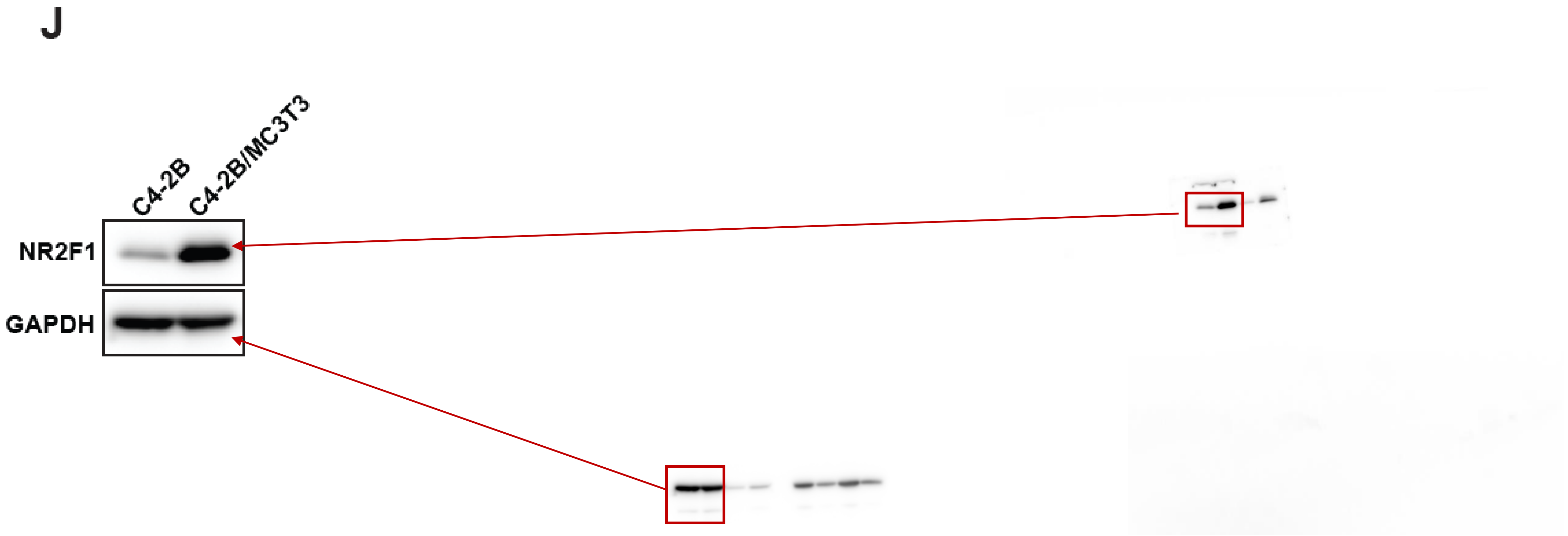

Figure 5

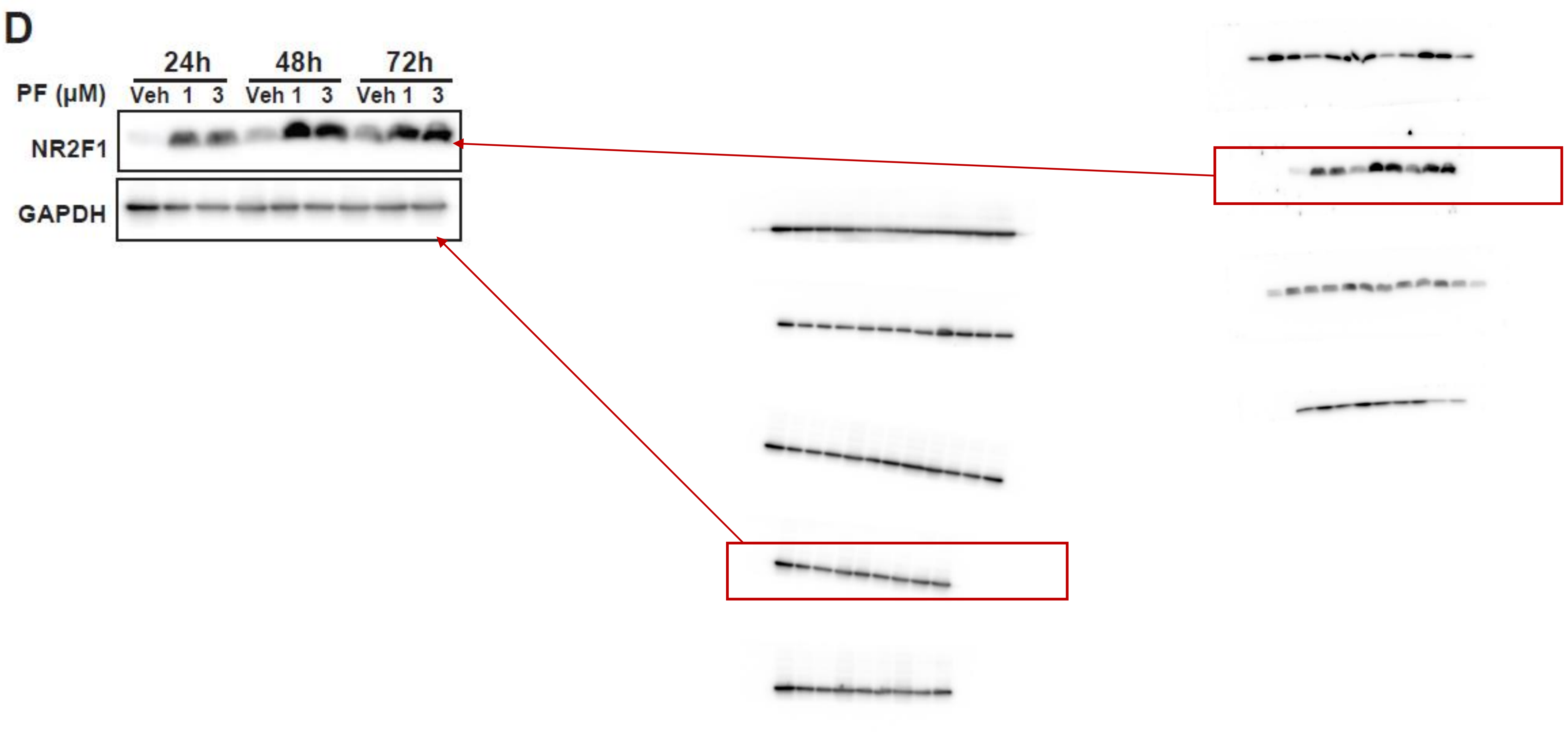

Figure 6

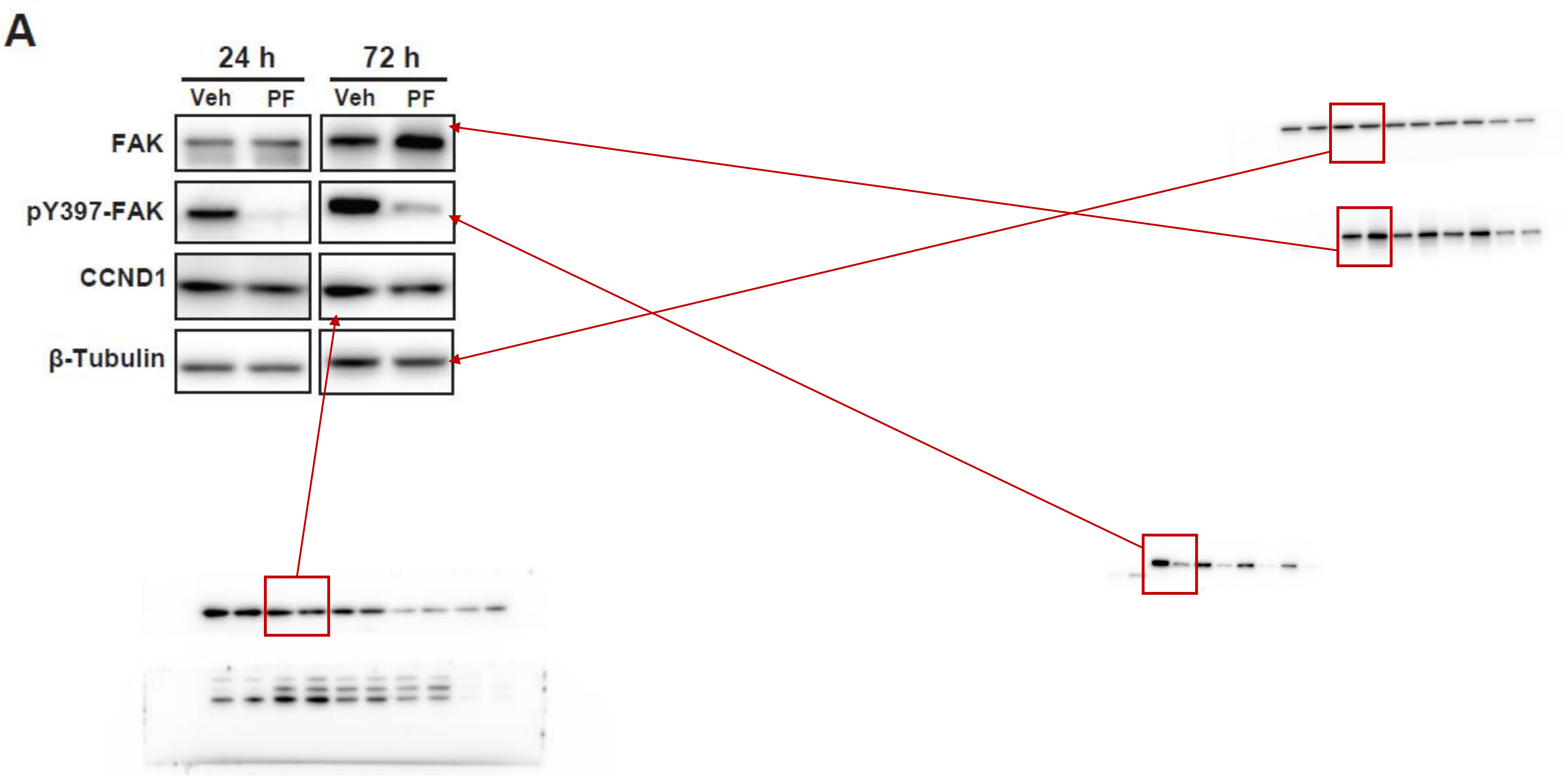

Figure 6

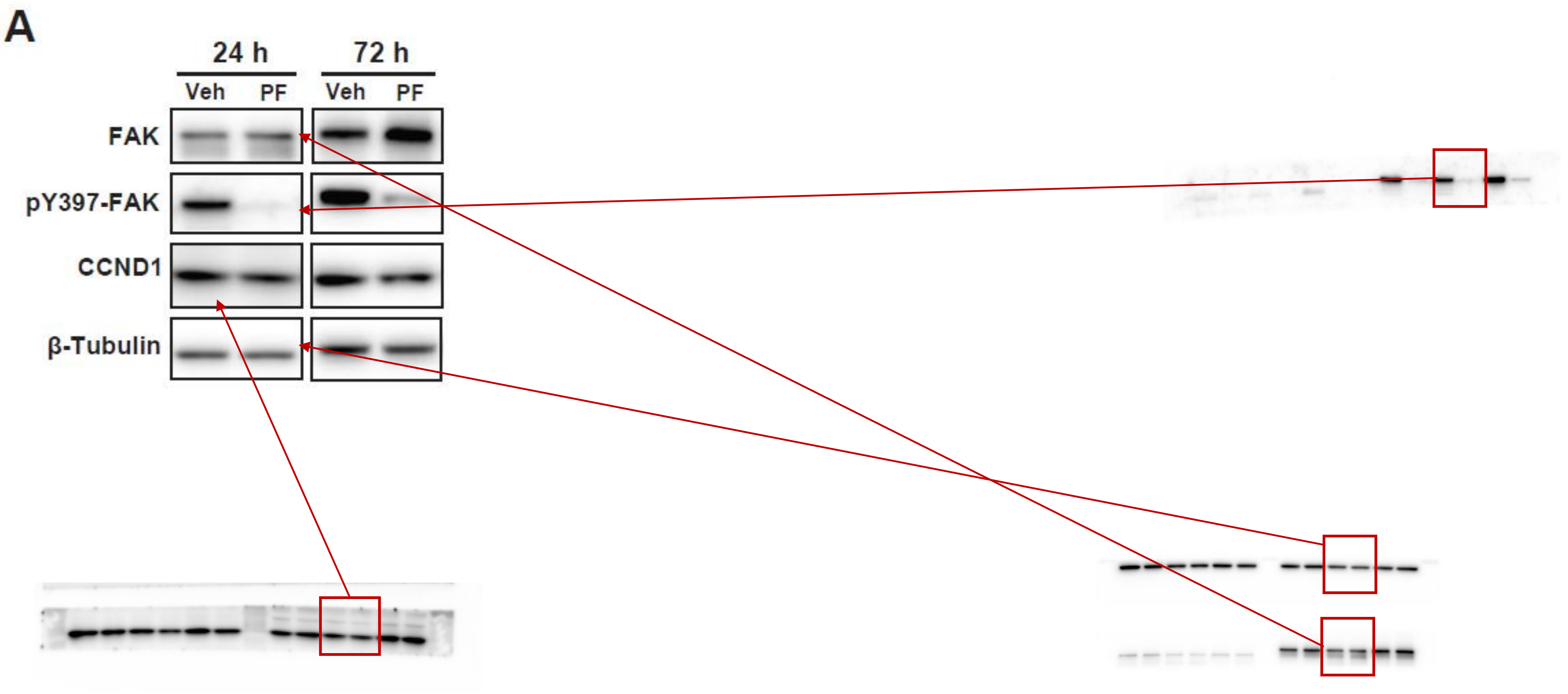

Figure 6

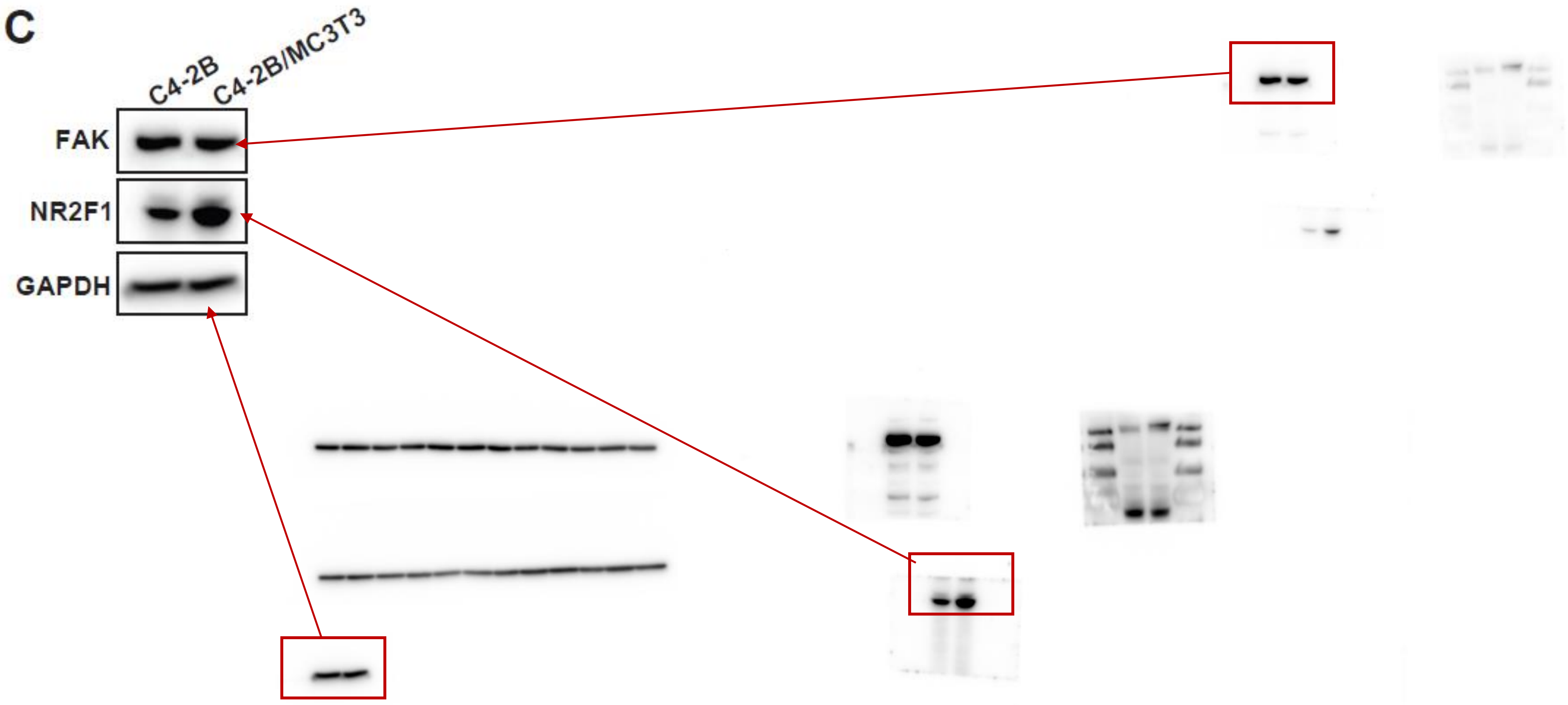

Figure S1

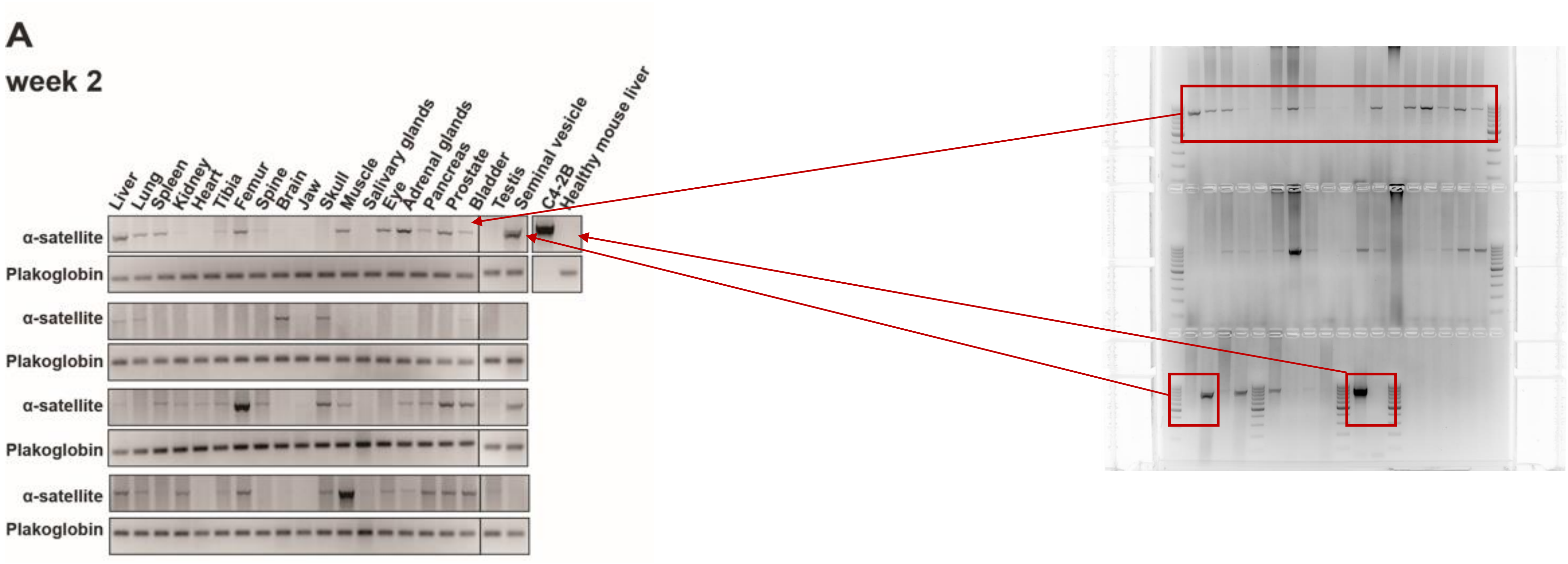
A
week 2
Liver
Lung
Spleen
Kidney
Heart
Tibia
Femur
Spine
Brain
Jaw
Skull
Muscle
Salivary glands
Eye
Adrenal glands
Pancreas
Prostate
Bladder
Testis
Seminal vesicle
C4-2B
Healthy mouse liver
α-satellite
Plakoglobin
α-satellite
Plakoglobin
α-satellite
Plakoglobin
α-satellite
Plakoglobin

Figure S1

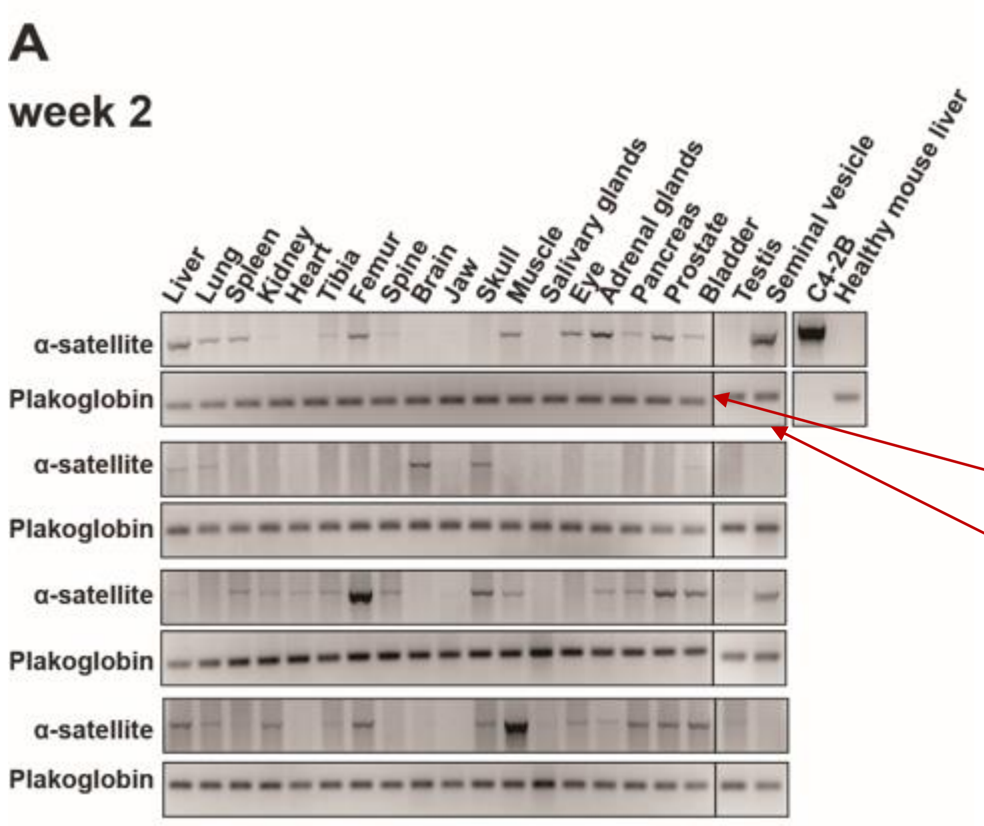
A
week 2
Liver
Lung
Spleen
Kidney
Heart
Tibia
Femur
Spine
Brain
Jaw
Skull
Muscle
Salivary glands
Eye
Adrenal glands
Pancreas
Prostate
Bladder
Testis
Seminal vesicle
C4-2B
Healthy mouse liver
α-satellite
Plakoglobin
α-satellite
Plakoglobin
α-satellite
Plakoglobin
α-satellite
Plakoglobin

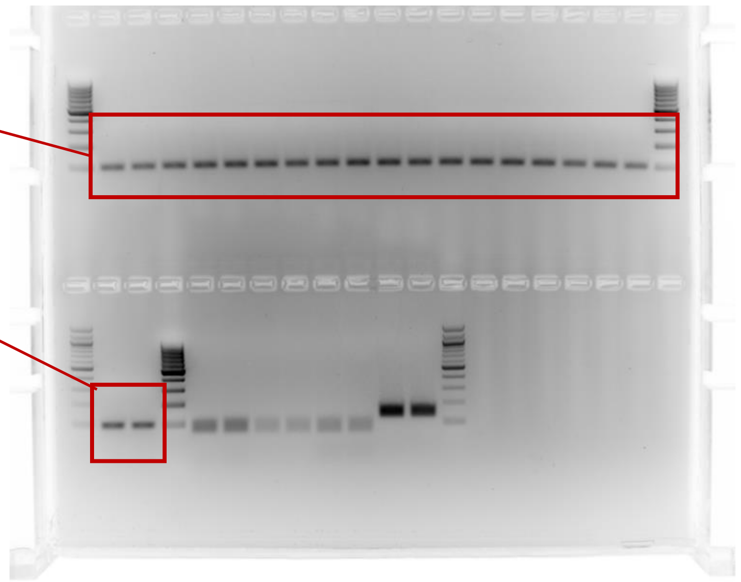

Figure S1

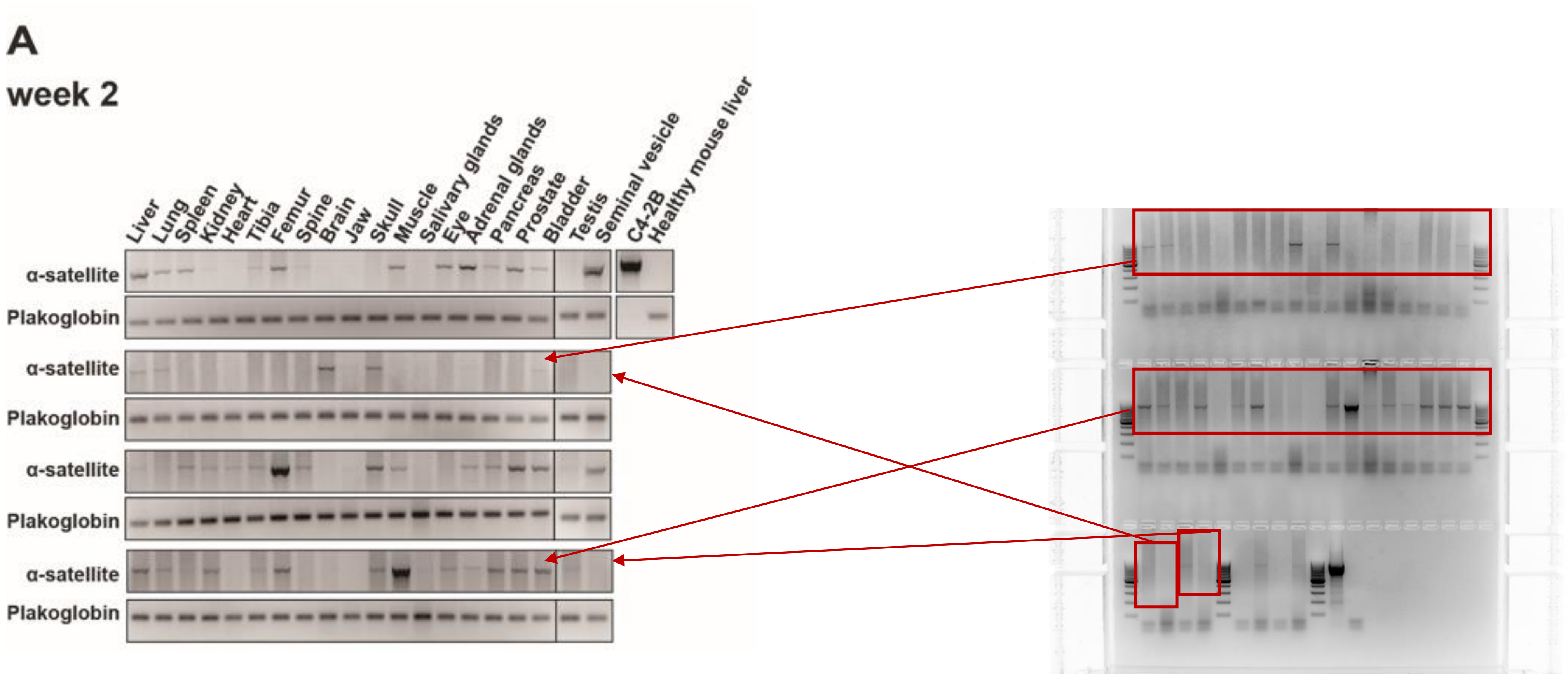
A
week 2
Liver
Lung
Spleen
Kidney
Heart
Tibia
Femur
Spine
Brain
Jaw
Skull
Muscle
Salivary glands
Eye
Adrenal glands
Pancreas
Prostate
Bladder
Testis
Seminal vesicle
C4-2B
Healthy mouse liver
α-satellite
Plakoglobin
α-satellite
Plakoglobin
α-satellite
Plakoglobin
α-satellite
Plakoglobin

Figure S1

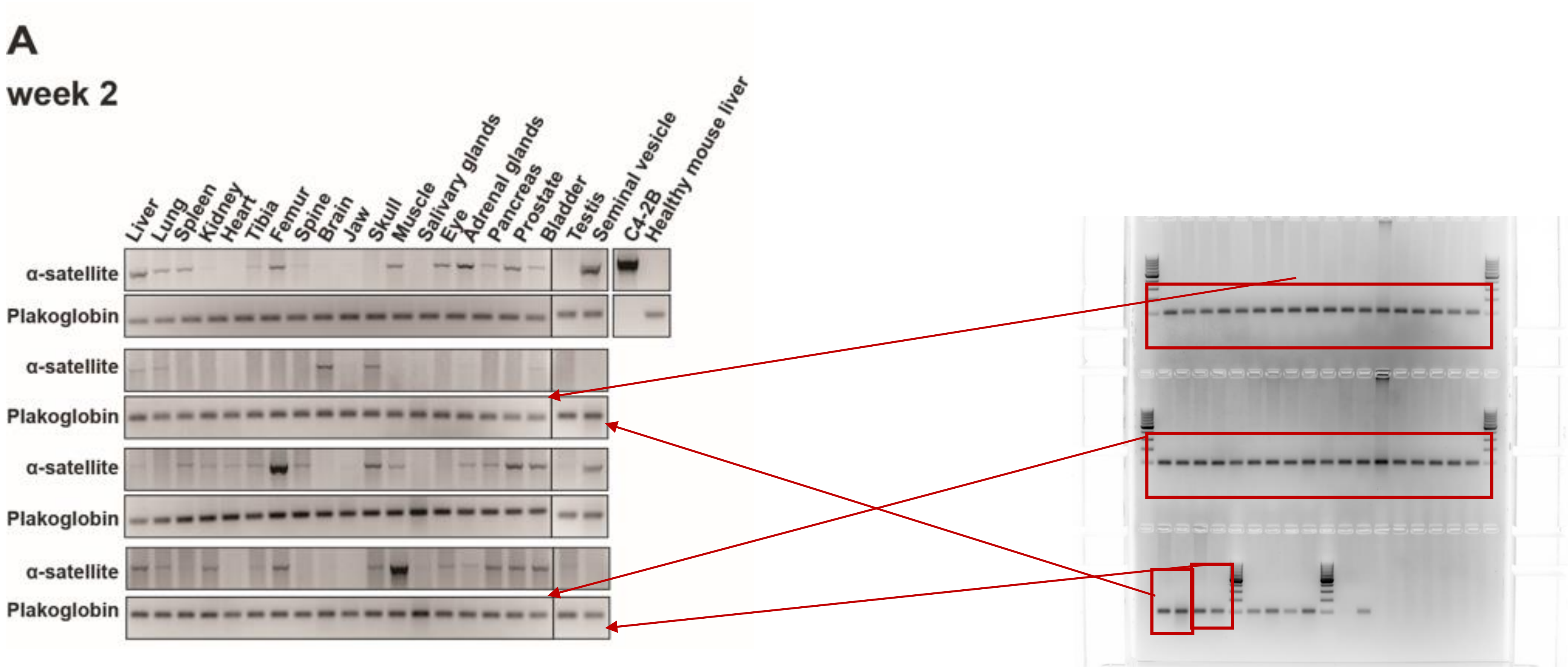
A
week 2
Liver
Lung
Spleen
Kidney
Heart
Tibia
Femur
Spine
Brain
Jaw
Skull
Muscle
Salivary glands
Eye
Adrenal glands
Pancreas
Prostate
Bladder
Testis
Seminal vesicle
C4-2B
Healthy mouse liver
α-satellite
Plakoglobin
α-satellite
Plakoglobin
α-satellite
Plakoglobin
α-satellite
Plakoglobin

Figure S1

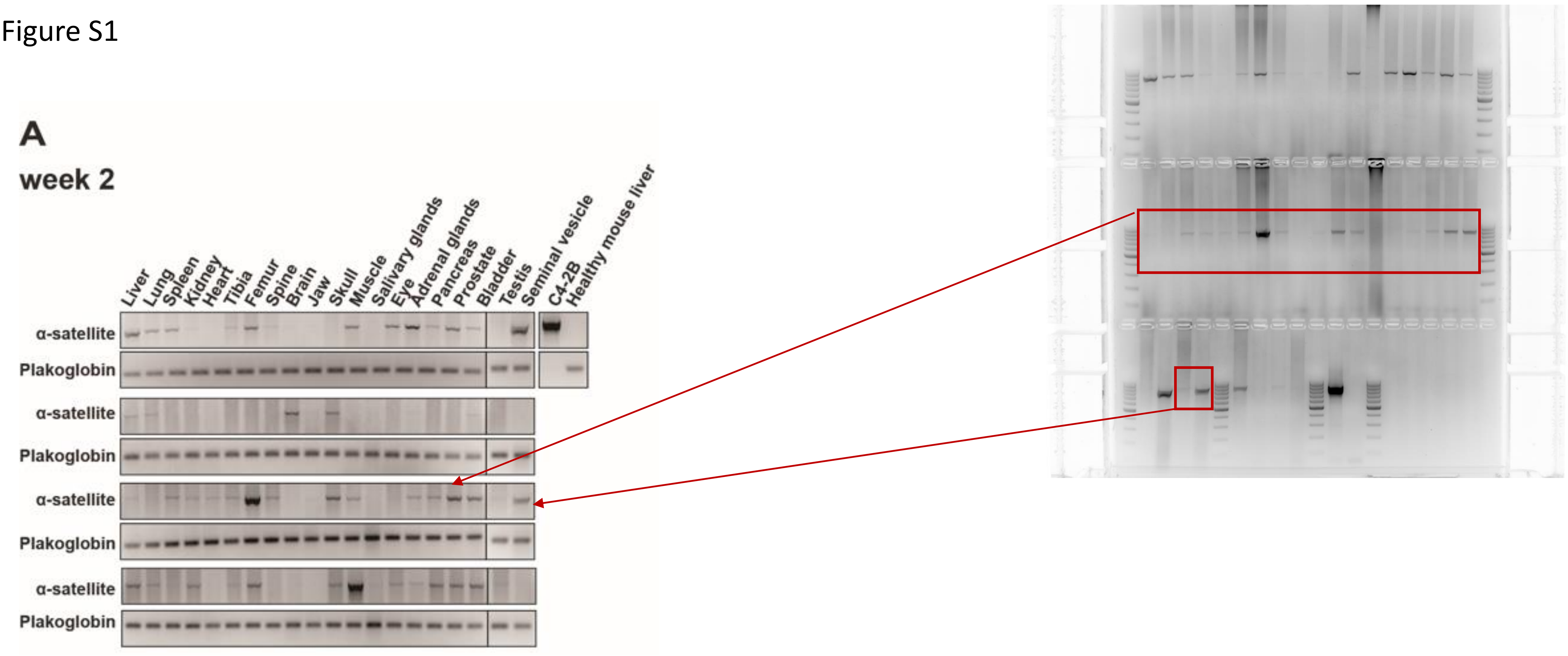
A
week 2
Liver
Lung
Spleen
Kidney
Heart
Tibia
Femur
Spine
Brain
Jaw
Skull
Muscle
Salivary glands
Eye
Adrenal glands
Pancreas
Prostate
Bladder
Testis
Seminal vesicle
C4-2B
Healthy mouse liver
α-satellite
Plakoglobin
α-satellite
Plakoglobin
α-satellite
Plakoglobin
α-satellite
Plakoglobin

Figure S1

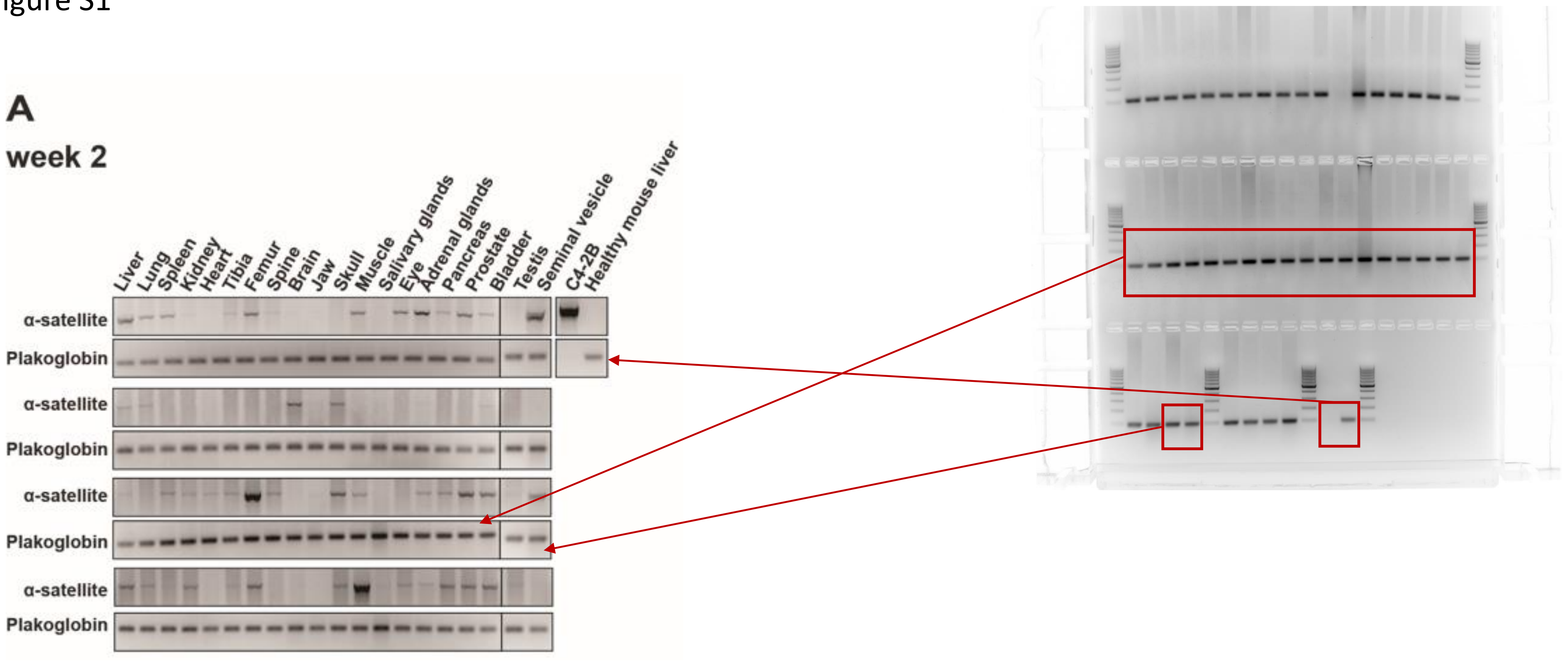
A
week 2
Liver
Lung
Spleen
Kidney
Heart
Tibia
Femur
Spine
Brain
Jaw
Skull
Muscle
Salivary glands
Eye
Adrenal glands
Pancreas
Prostate
Bladder
Testis
Seminal vesicle
C4-2B
Healthy mouse liver
α-satellite
Plakoglobin
α-satellite
Plakoglobin
α-satellite
Plakoglobin
α-satellite
Plakoglobin

Figure S1

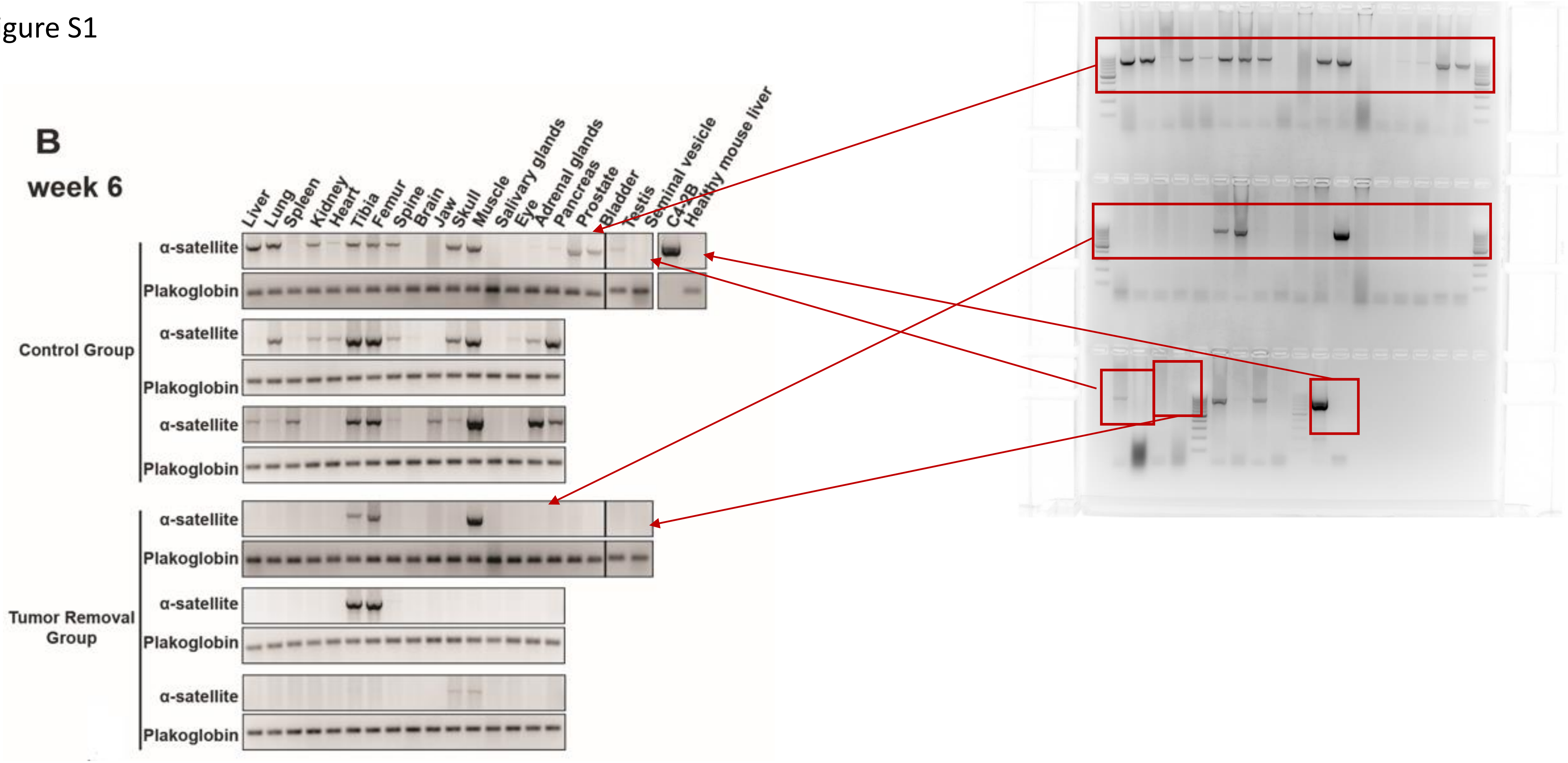
B
week 6
Liver
Lung
Spleen
Kidney
Heart
Tibia
Femur
Spine
Brain
Jaw
Skull
Muscle
Salivary glands
Eye
Adrenal glands
Pancreas
Prostate
Bladder
Testis
Seminal vesicle
C4-2B
Healthy mouse liver
α-satellite
Plakoglobin
Control Group
Tumor Removal Group

### Figure S1

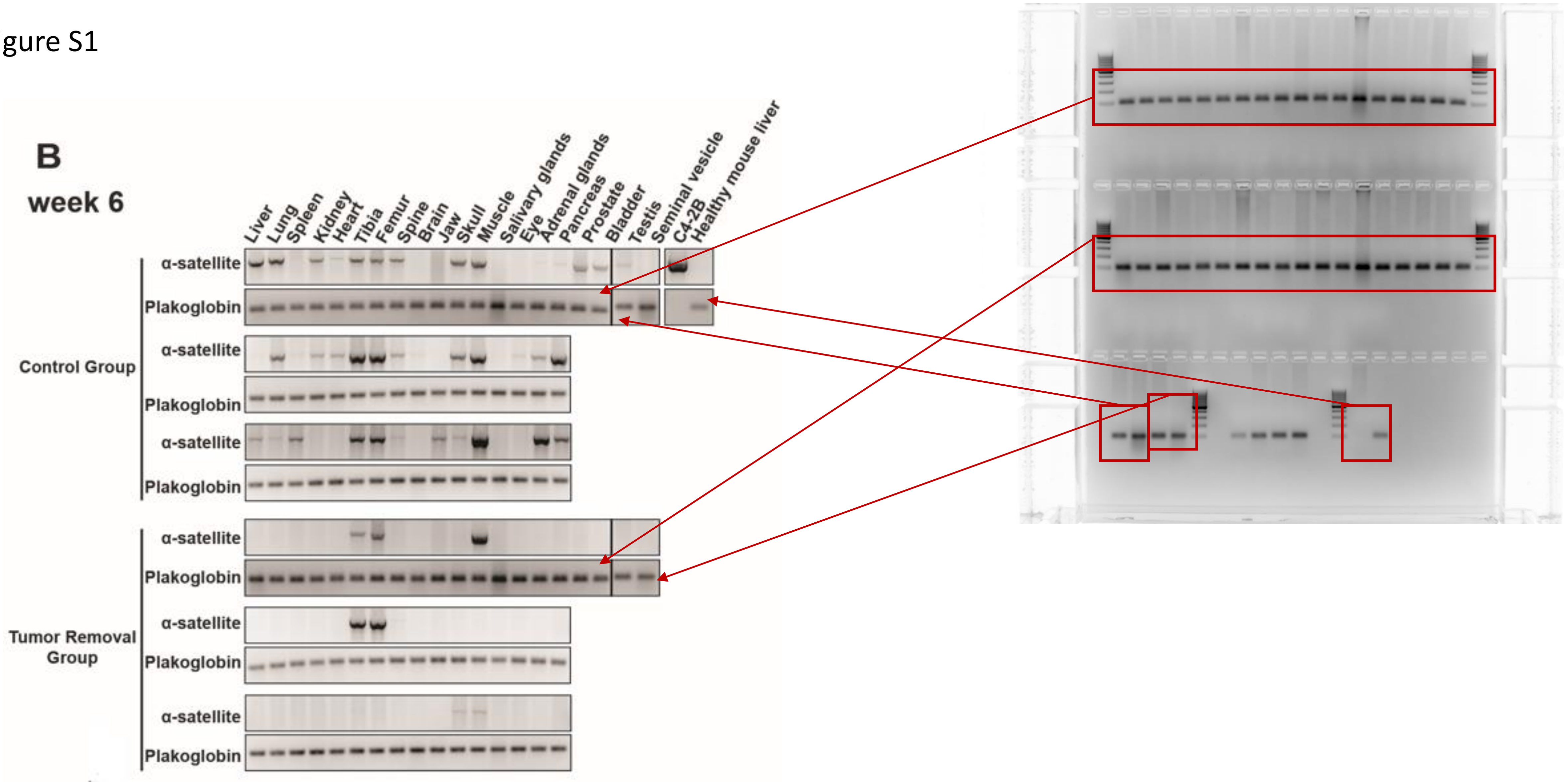
B
week 6
Liver
Lung
Spleen
Kidney
Heart
Tibia
Femur
Spine
Brain
Jaw
Skull
Muscle
Salivary glands
Eye
Adrenal glands
Pancreas
Prostate
Bladder
Testis
Seminal vesicle
C4-2B
Healthy mouse liver
α-satellite
Plakoglobin
Control Group
Tumor Removal Group

Figure S1

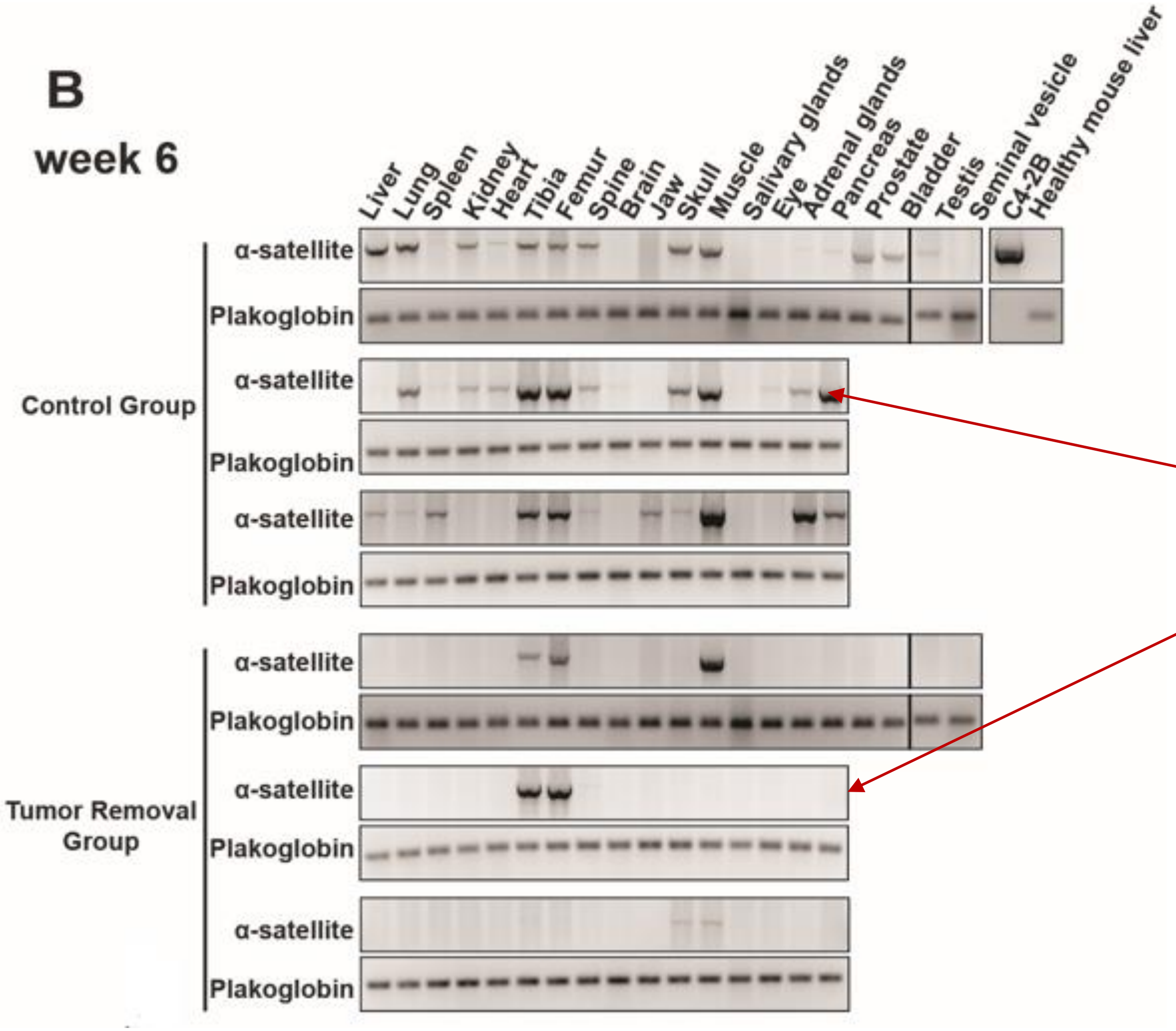

B
week 6
Liver
Lung
Spleen
Kidney
Heart
Tibia
Femur
Spine
Brain
Jaw
Skull
Muscle
Salivary glands
Eye
Adrenal glands
Pancreas
Prostate
Bladder
Testis
Seminal vesicle
C4-2B
Healthy mouse liver
Control Group
Tumor Removal Group
α-satellite
Plakoglobin


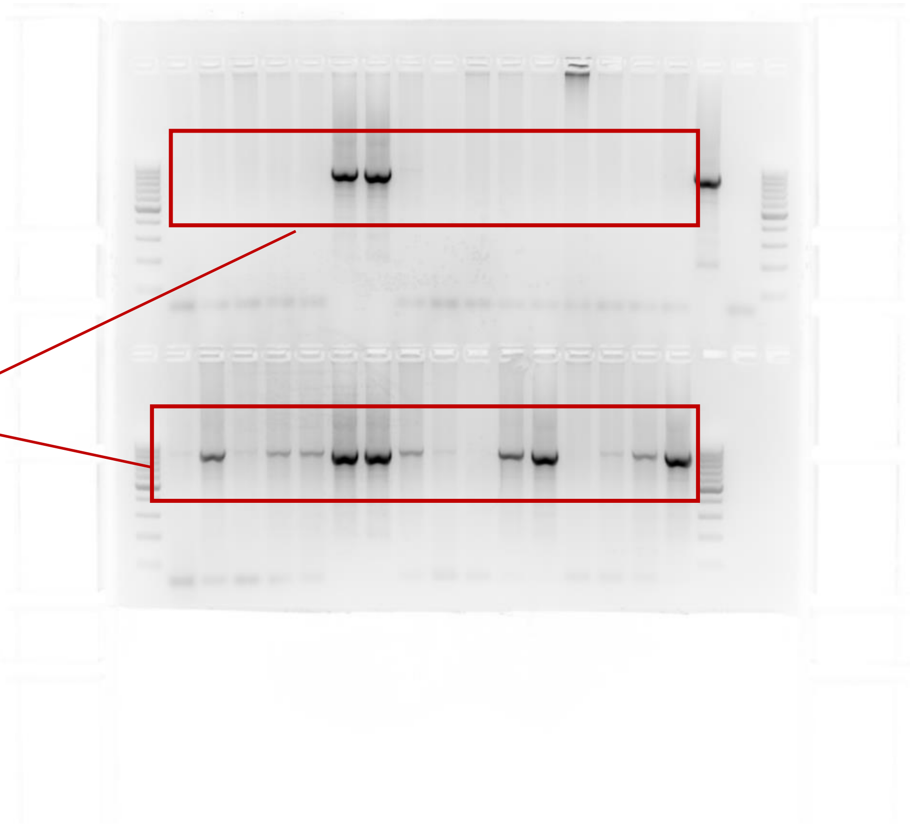

Figure S1

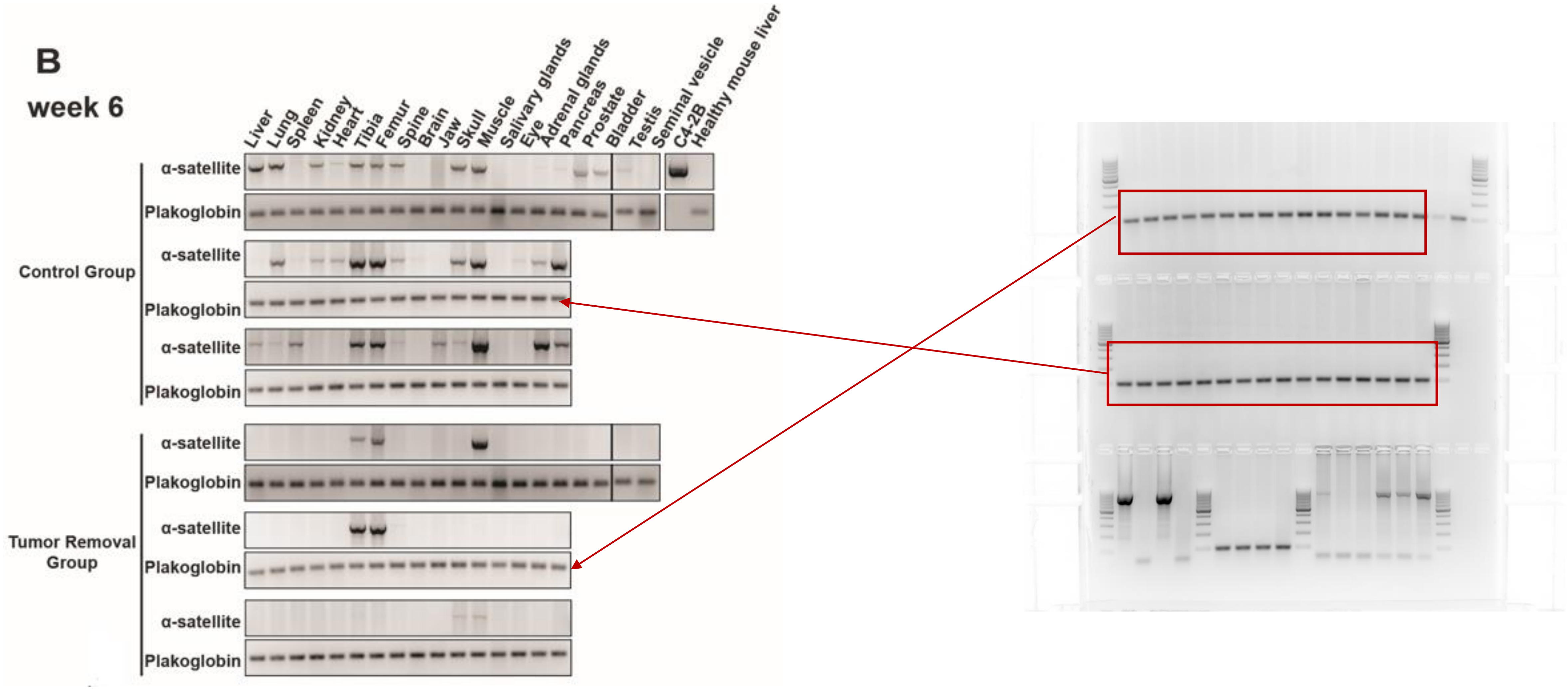
B
week 6
Liver
Lung
Spleen
Kidney
Heart
Tibia
Femur
Spine
Brain
Jaw
Skull
Muscle
Salivary glands
Eye
Adrenal glands
Pancreas
Prostate
Bladder
Testis
Seminal vesicle
C4-2B
Healthy mouse liver
Control Group
Tumor Removal Group
α-satellite
Plakoglobin

Figure S1

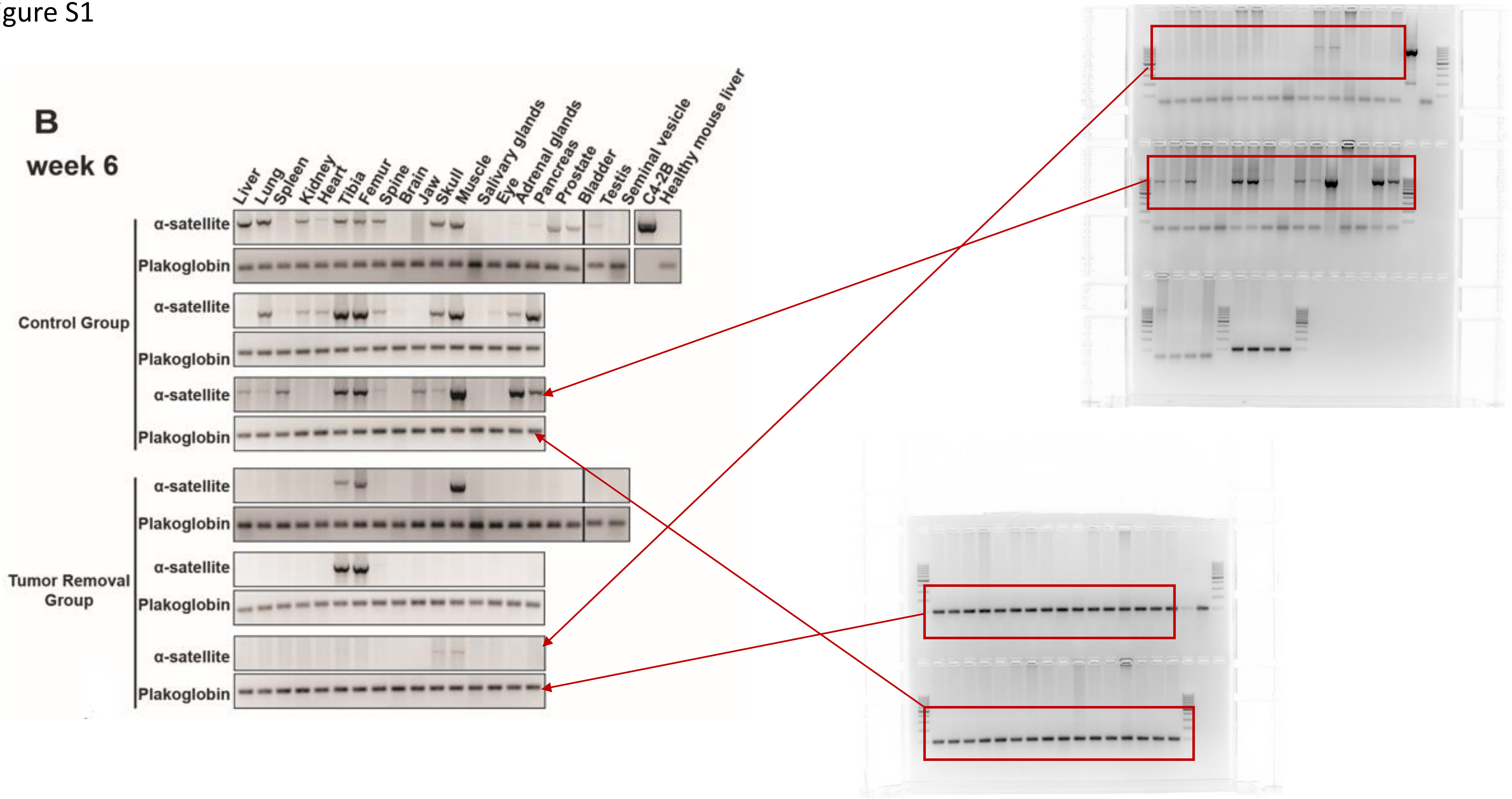

B
week 6
Liver
Lung
Spleen
Kidney
Heart
Tibia
Femur
Spine
Brain
Jaw
Skull
Muscle
Salivary glands
Eye
Adrenal glands
Pancreas
Prostate
Bladder
Testis
Seminal vesicle
C4-2B
Healthy mouse liver
Control Group
Tumor Removal Group
α-satellite
Plakoglobin

Figure S1

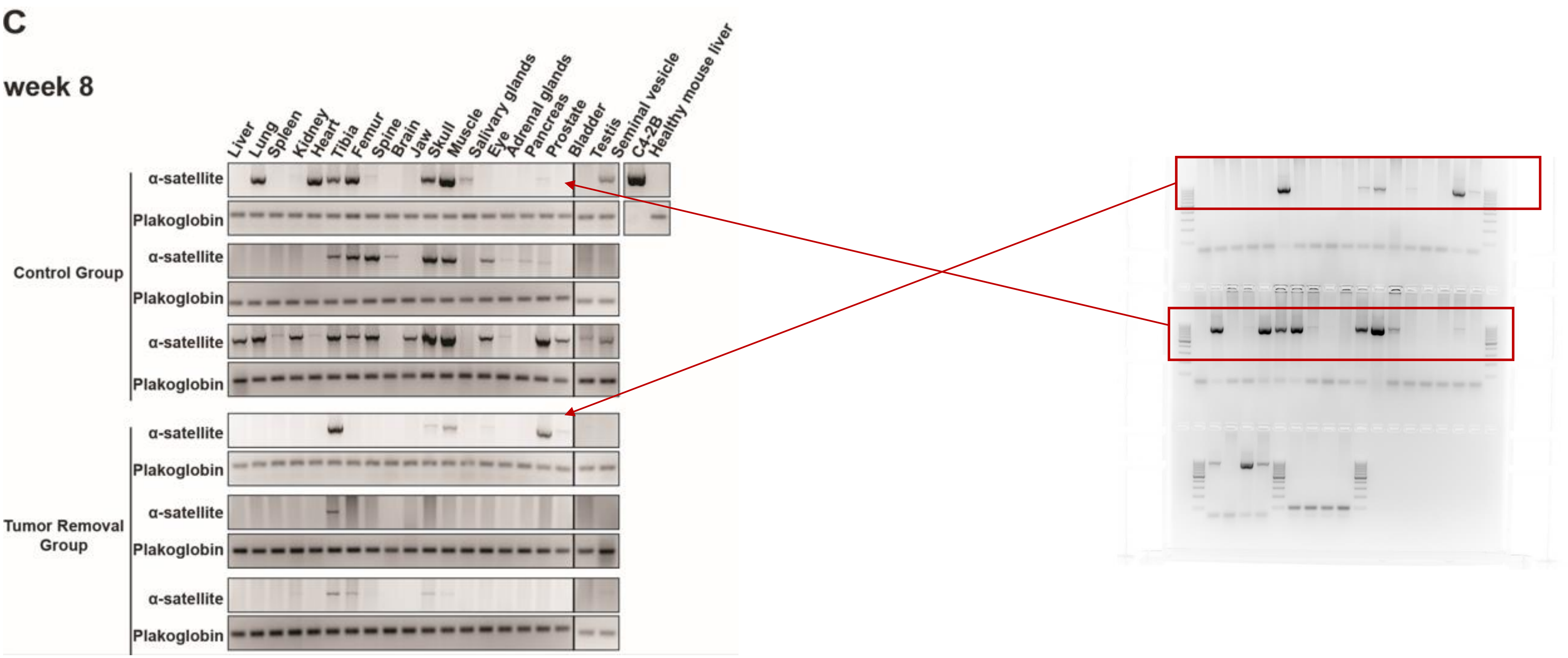
C
week 8
Liver
Lung
Spleen
Kidney
Heart
Tibia
Femur
Spine
Brain
Jaw
Skull
Muscle
Salivary glands
Eye
Adrenal glands
Pancreas
Prostate
Bladder
Testis
Seminal vesicle
C4-2B
Healthy mouse liver
α-satellite
Plakoglobin
Control Group
Tumor Removal Group

Figure S1

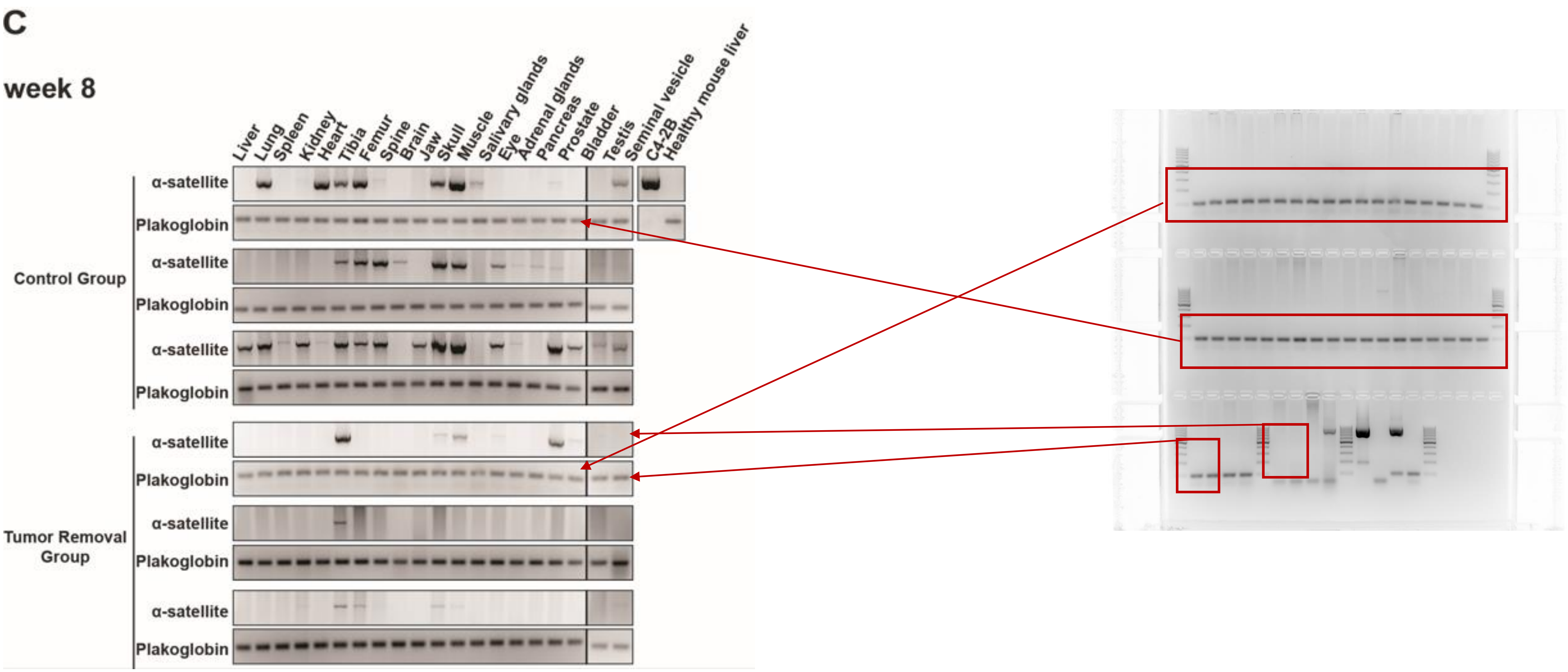
C
week 8
Liver
Lung
Spleen
Kidney
Heart
Tibia
Femur
Spine
Brain
Jaw
Skull
Muscle
Salivary glands
Eye
Adrenal glands
Pancreas
Prostate
Bladder
Testis
Seminal vesicle
C4-2B
Healthy mouse liver
α-satellite
Plakoglobin
Control Group
Tumor Removal Group

Figure S1

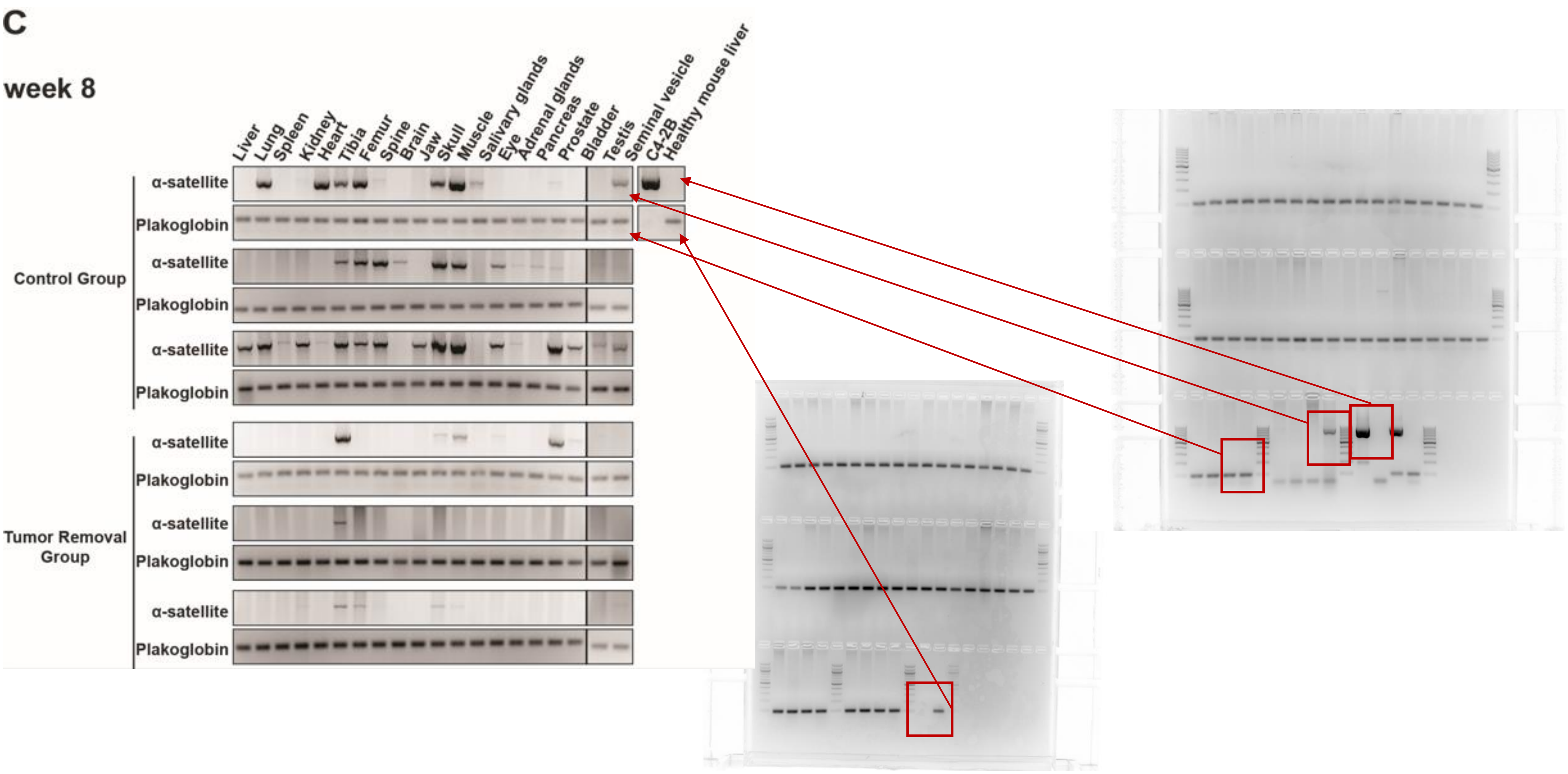
C
week 8
Liver
Lung
Spleen
Kidney
Heart
Tibia
Femur
Spine
Brain
Jaw
Skull
Muscle
Salivary glands
Eye
Adrenal glands
Pancreas
Prostate
Bladder
Testis
Seminal vesicle
C4-2B
Healthy mouse liver
α-satellite
Plakoglobin
α-satellite
Plakoglobin
α-satellite
Plakoglobin
Control Group
α-satellite
Plakoglobin
α-satellite
Plakoglobin
α-satellite
Plakoglobin
Tumor Removal Group

Figure S1

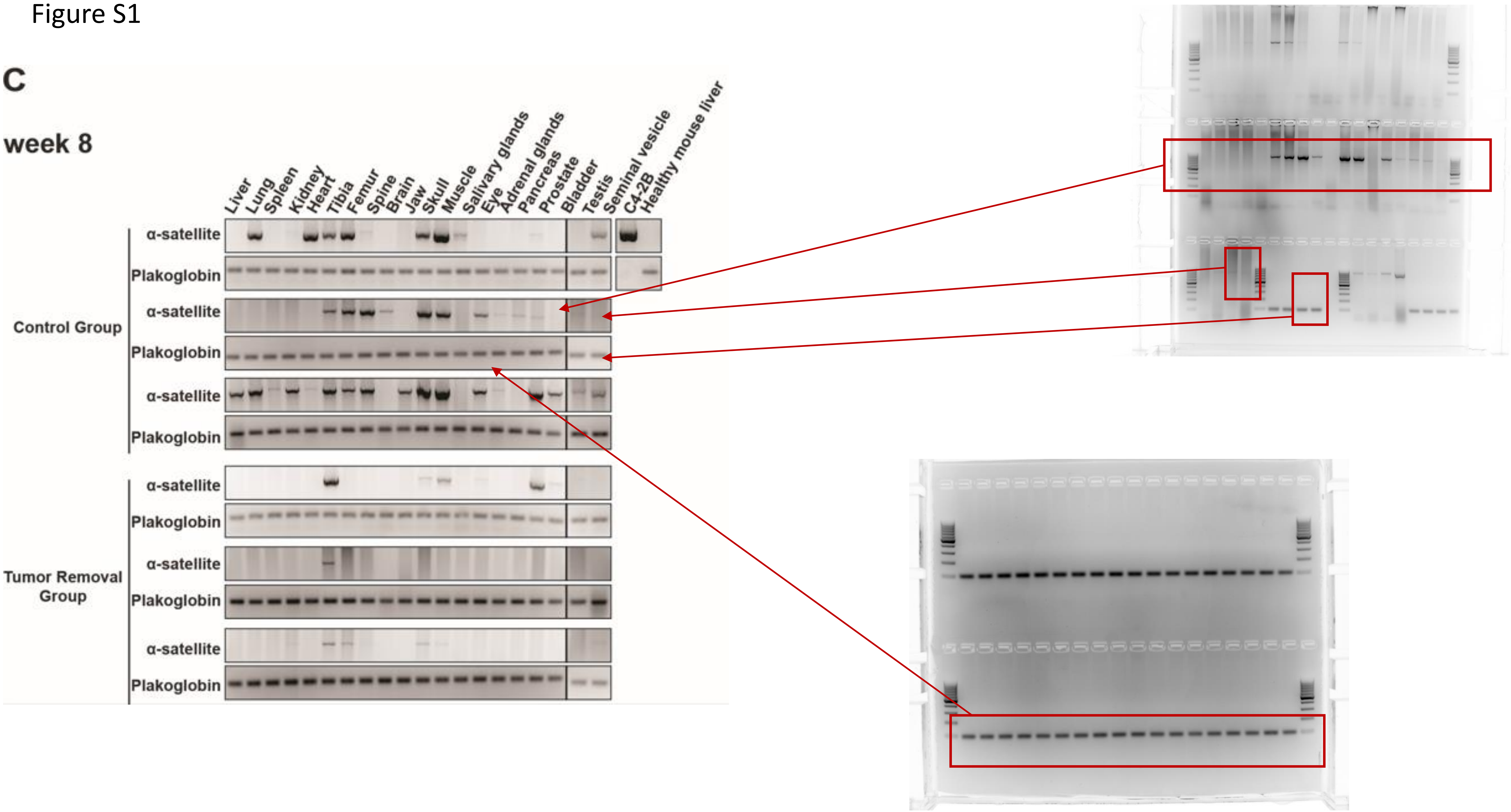
C
week 8
Liver
Lung
Spleen
Kidney
Heart
Tibia
Femur
Spine
Brain
Jaw
Skull
Muscle
Salivary glands
Eye
Adrenal glands
Pancreas
Prostate
Bladder
Testis
Seminal vesicle
C4-2B
Healthy mouse liver
Control Group
Tumor Removal Group
α-satellite
Plakoglobin

Figure S1

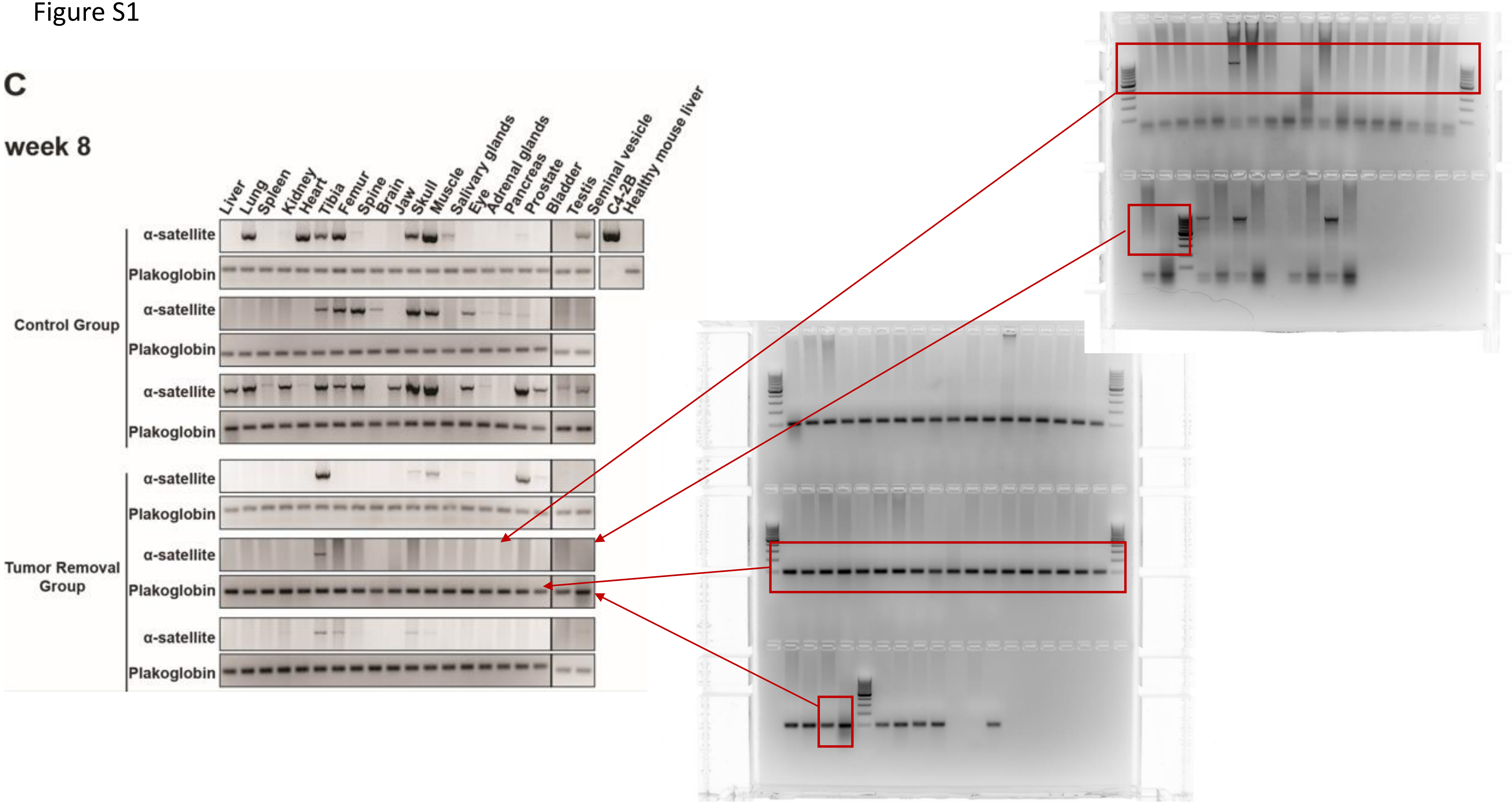

C
week 8
Liver
Lung
Spleen
Kidney
Heart
Tibia
Femur
Spine
Brain
Jaw
Skull
Muscle
Salivary glands
Eye
Adrenal glands
Pancreas
Prostate
Bladder
Testis
Seminal vesicle
C4-2B
Healthy mouse liver
Control Group
Tumor Removal Group
α-satellite
Plakoglobin

Figure S1

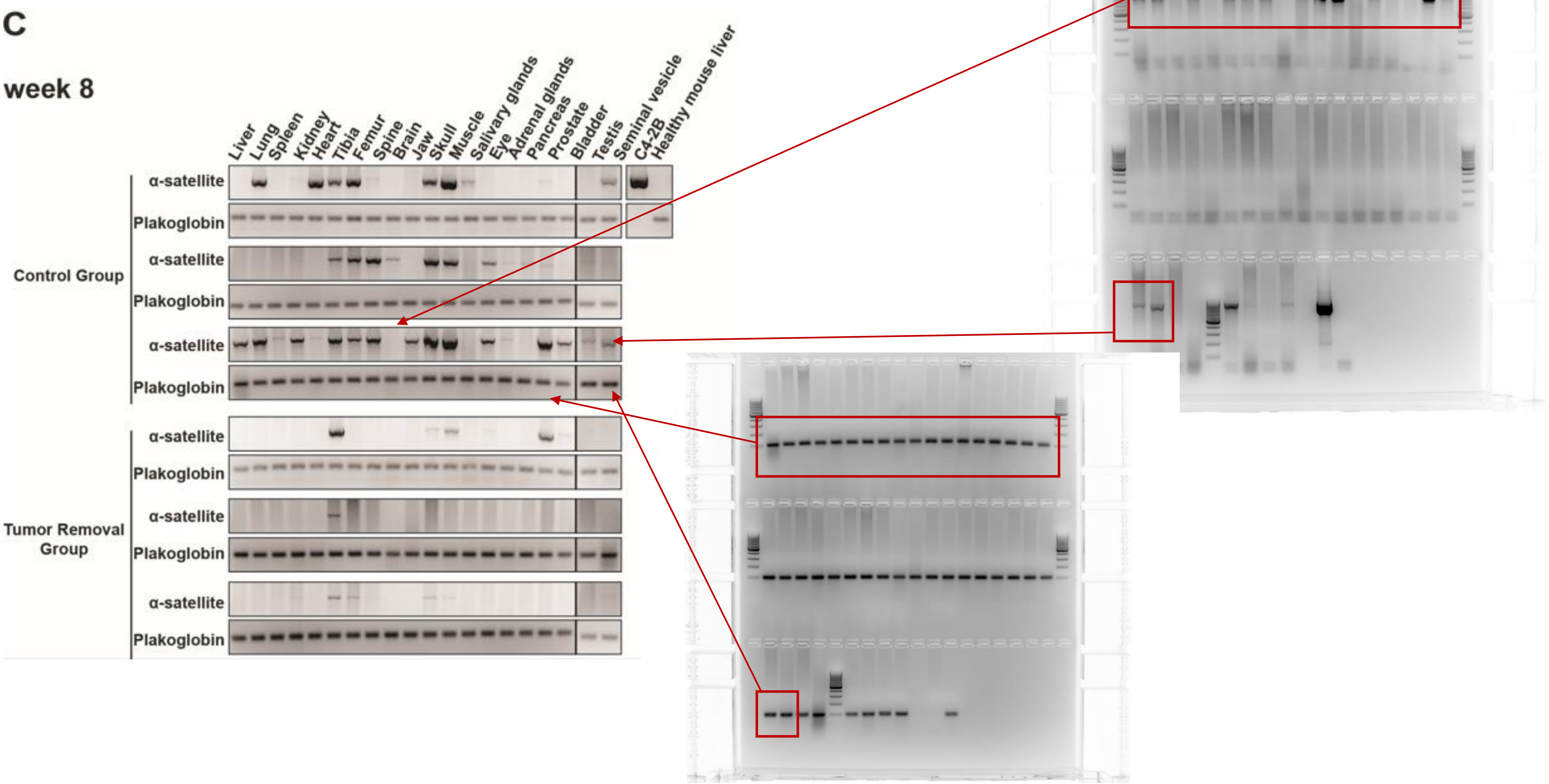
C
week 8
Liver
Lung
Spleen
Kidney
Heart
Tibia
Femur
Spine
Brain
Jaw
Skull
Muscle
Salivary glands
Eye
Adrenal glands
Pancreas
Prostate
Bladder
Testis
Seminal vesicle
C4-2B
Healthy mouse liver
Control Group
Tumor Removal Group
α-satellite
Plakoglobin

Figure S1

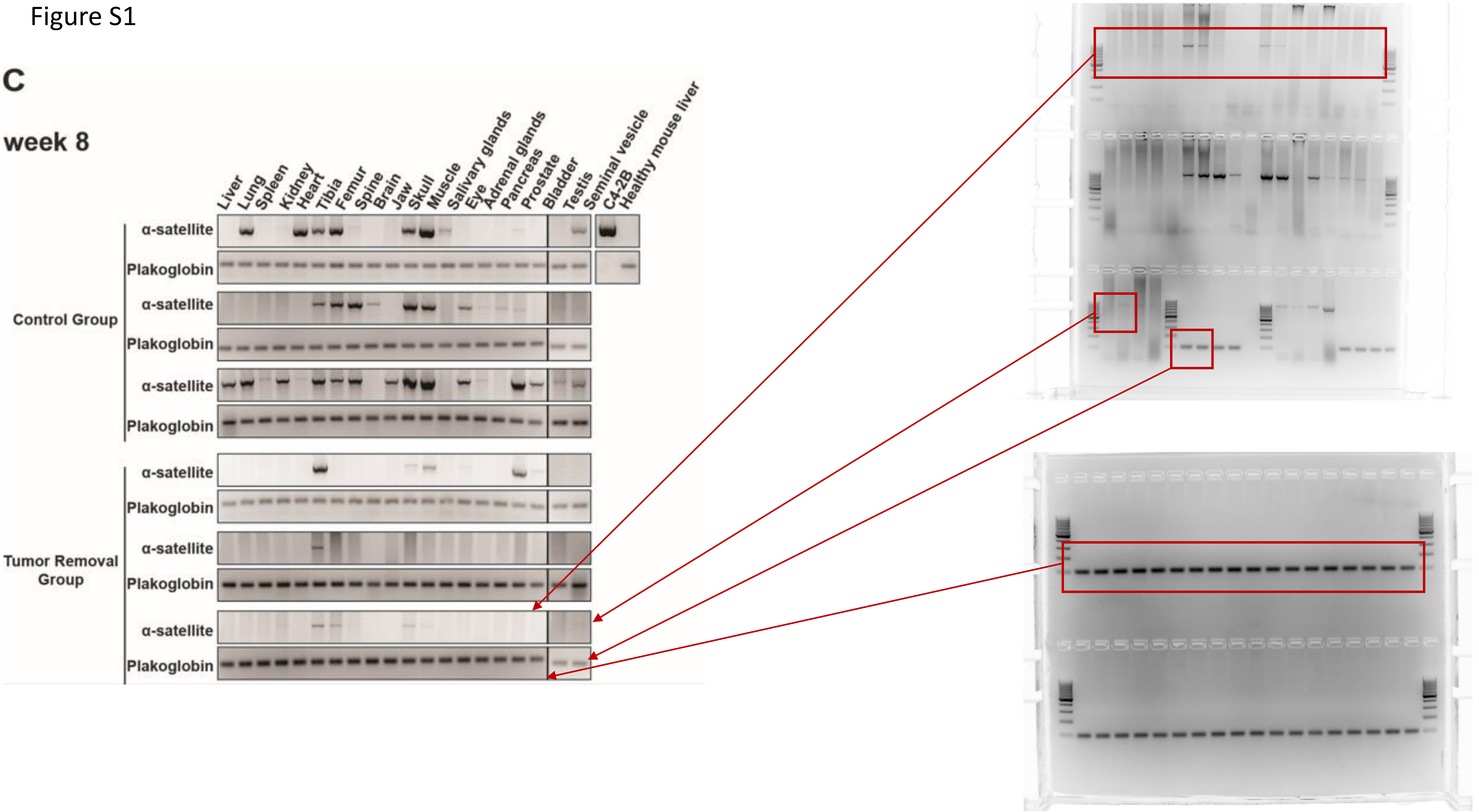
C
week 8
Liver
Lung
Spleen
Kidney
Heart
Tibia
Femur
Spine
Brain
Jaw
Skull
Muscle
Salivary glands
Eye
Adrenal glands
Pancreas
Prostate
Bladder
Testis
Seminal vesicle
C4-2B
Healthy mouse liver
α-satellite
Plakoglobin
Control Group
Tumor Removal Group

Figure S1

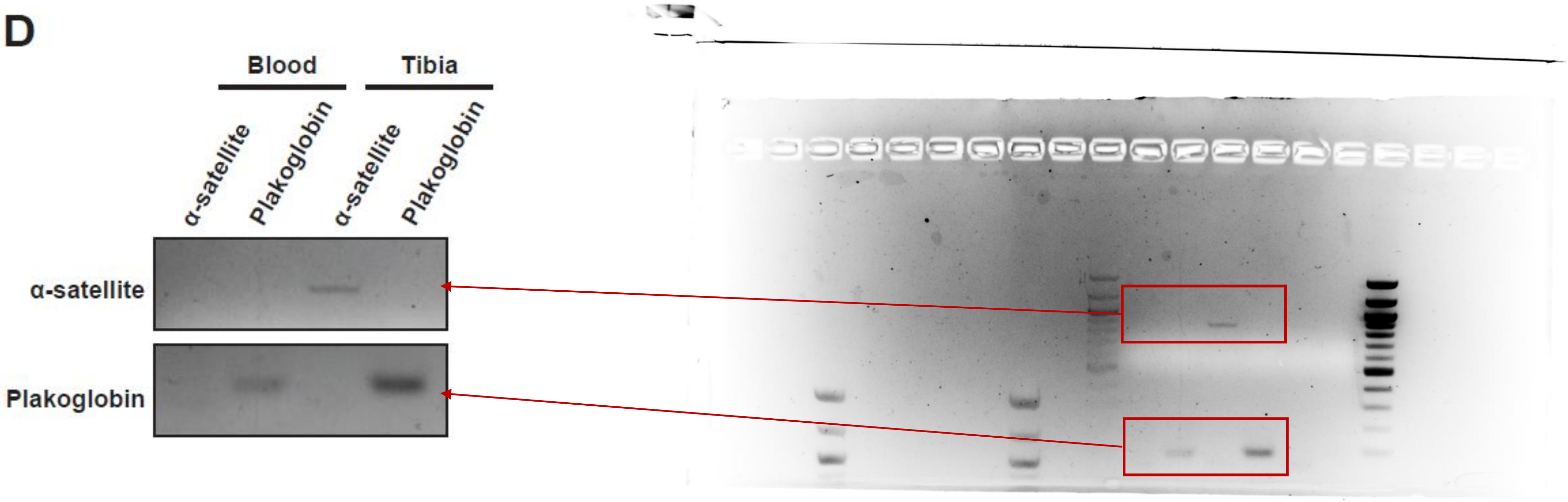
D
Blood
Tibia
α-satellite
Plakoglobin
α-satellite
Plakoglobin
α-satellite
Plakoglobin

Figure S3

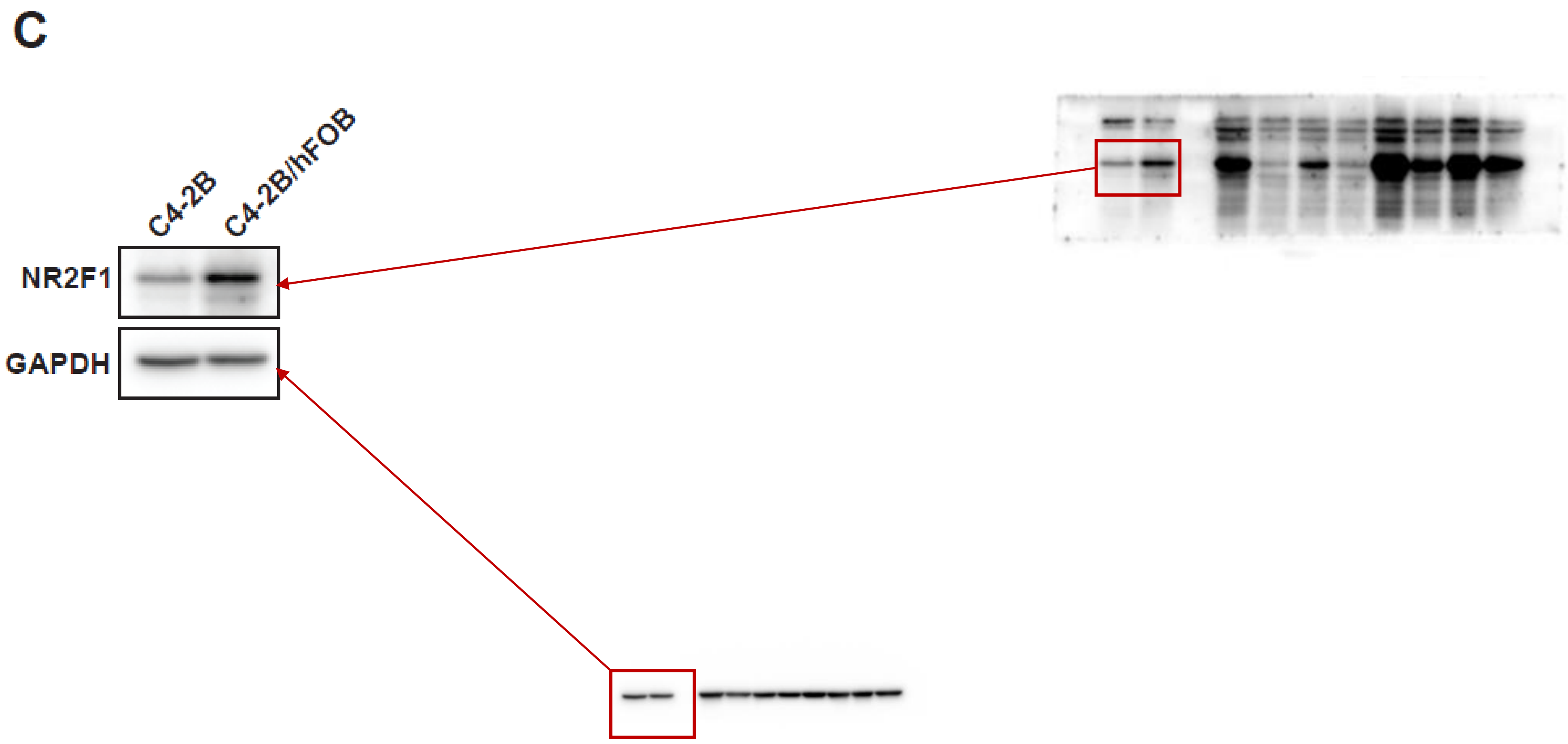

Figure S4

C

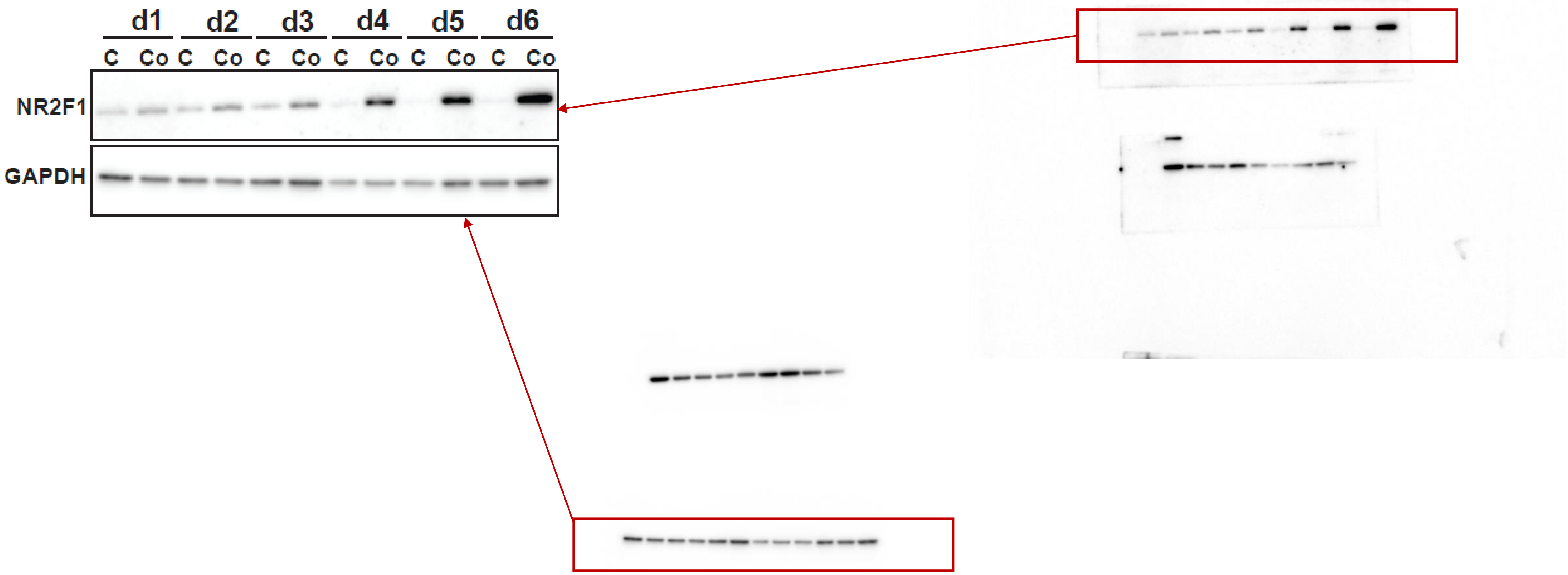

Figure S8

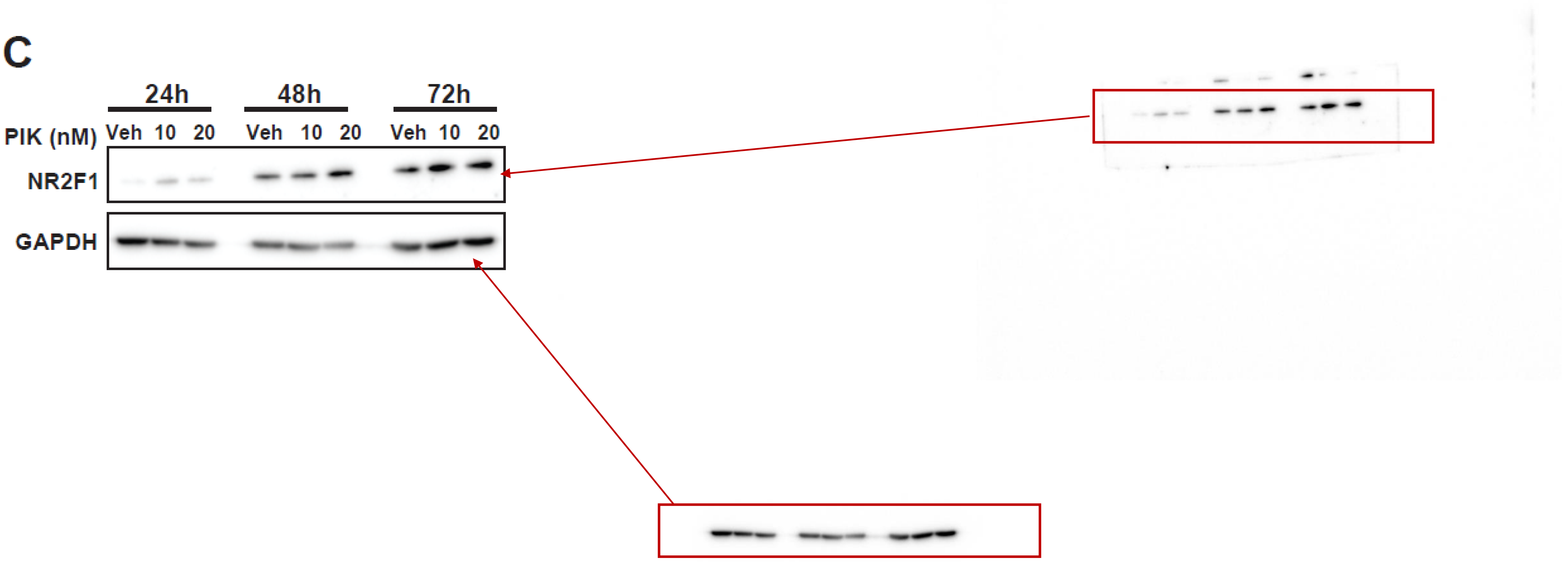

Figure S9

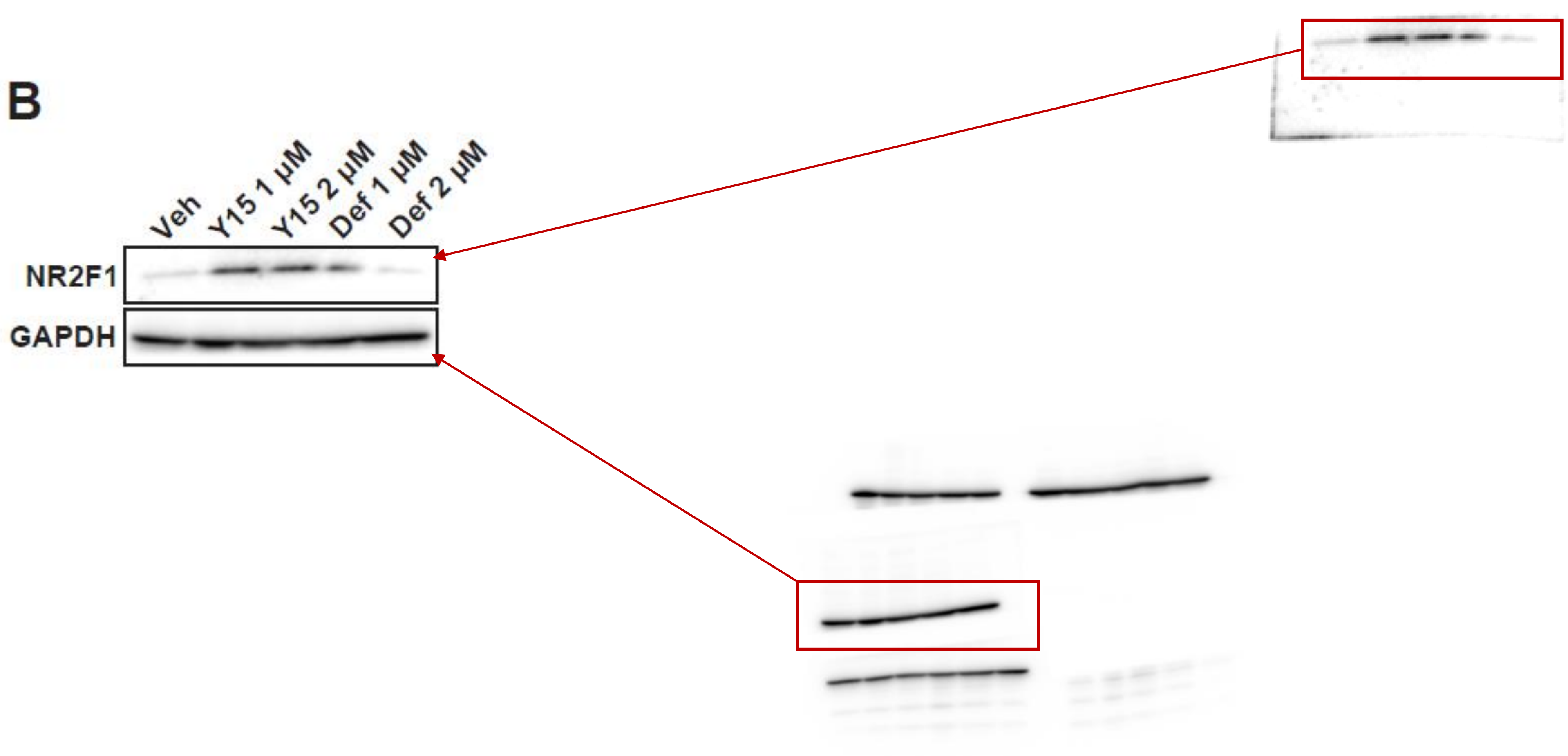

Figure S10

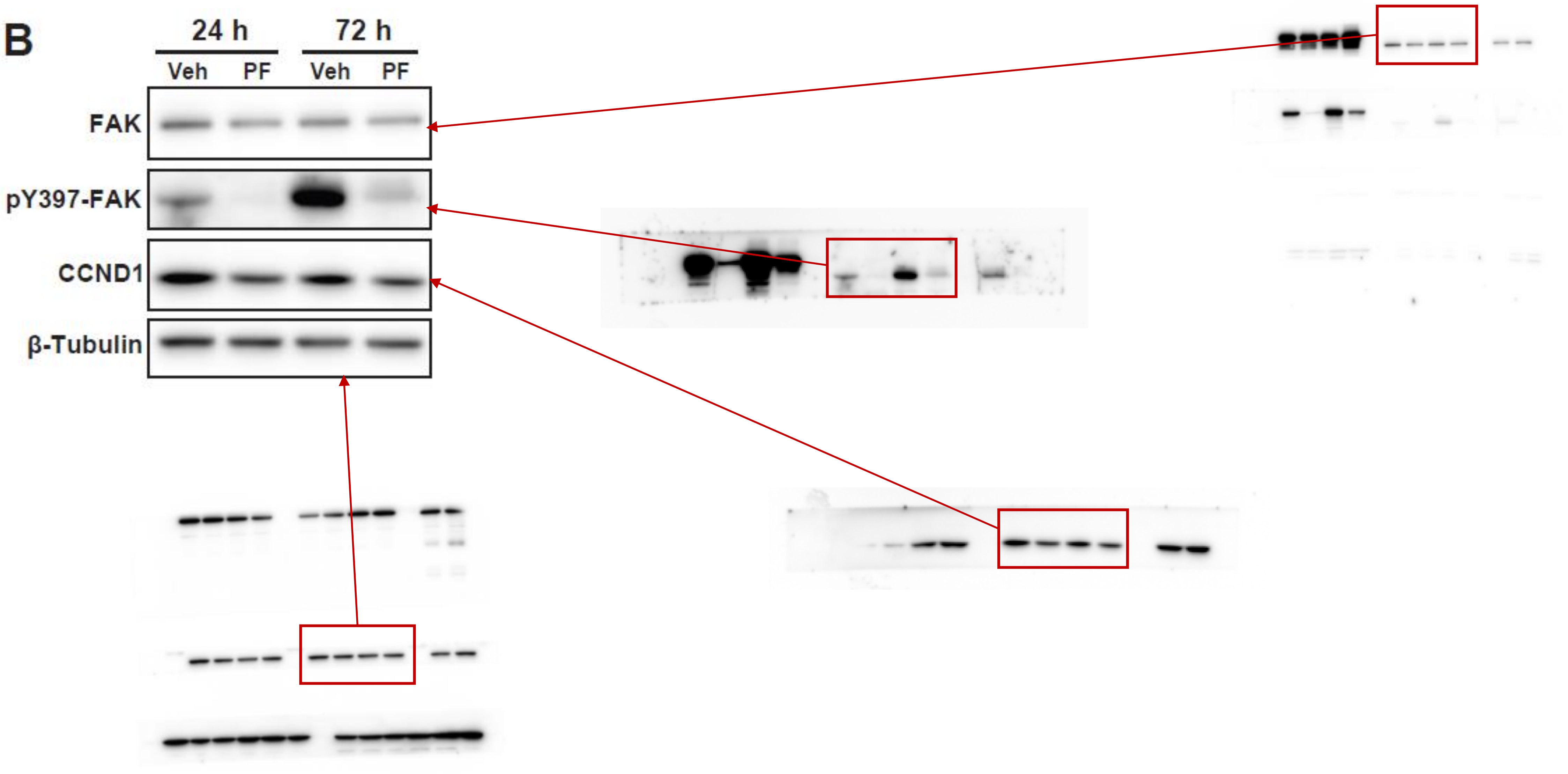

Figure S11

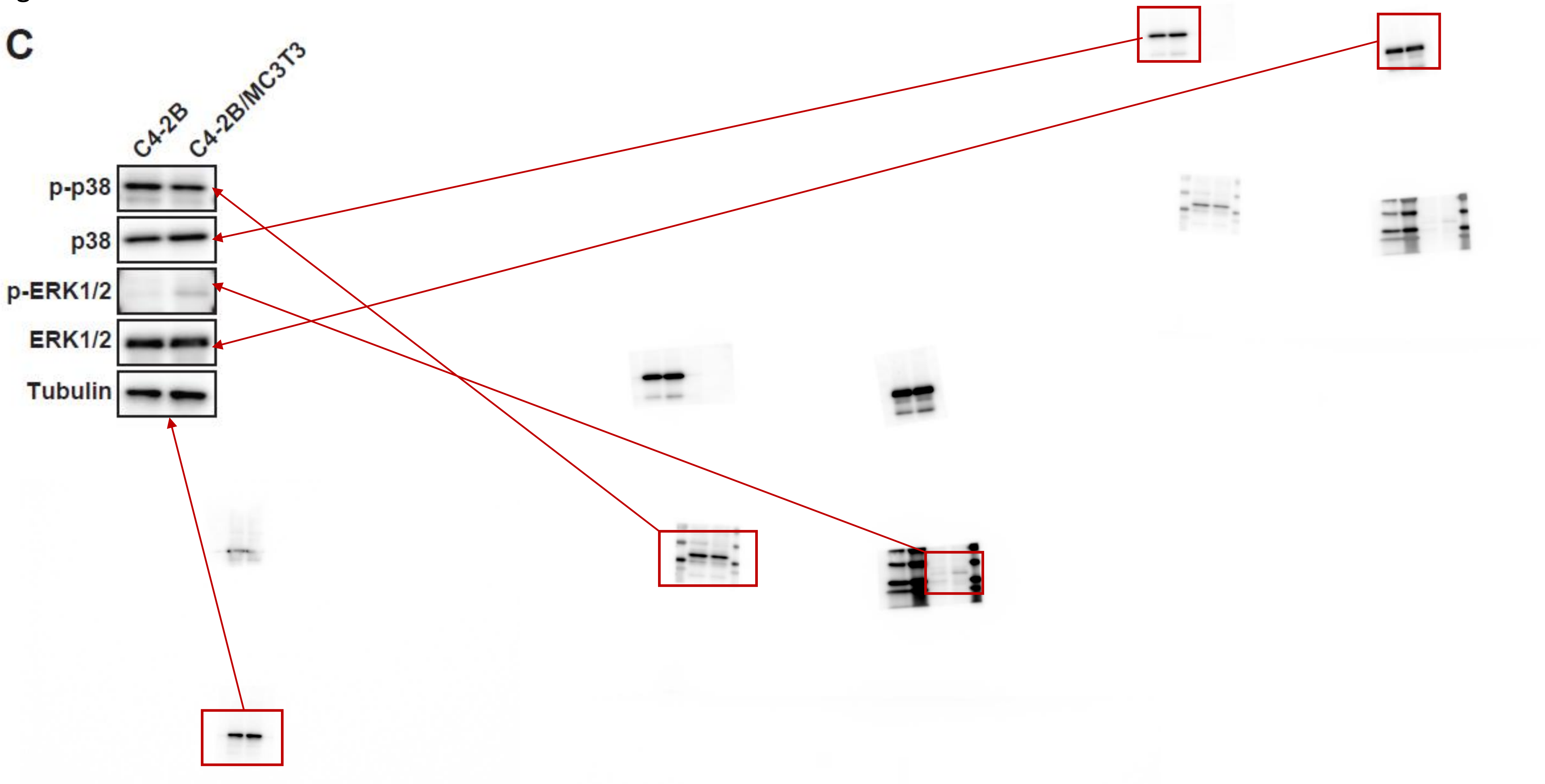

Species specificity Primer

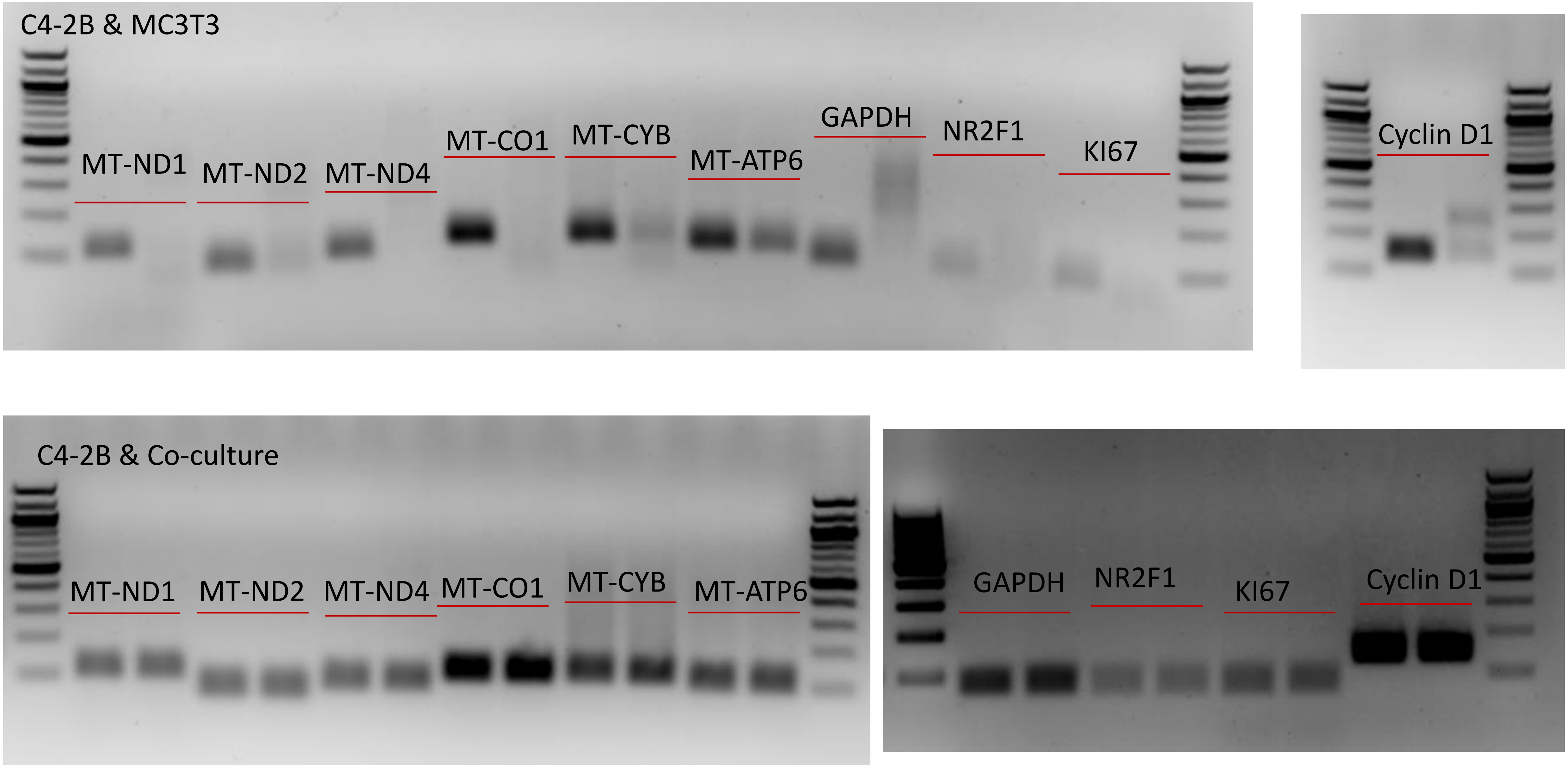

Species specificity Primer

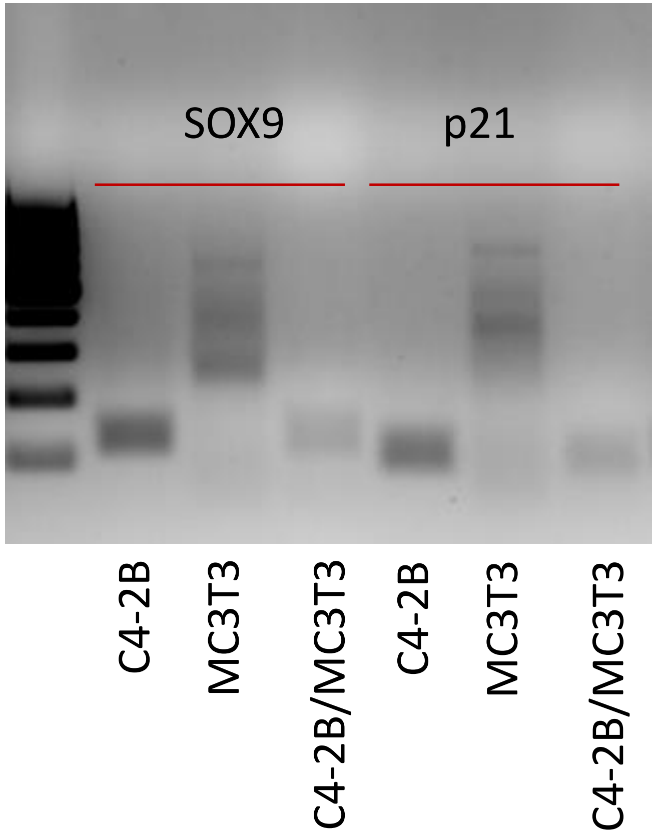
